# Ancient DNA reveals the origins of the Albanians

**DOI:** 10.1101/2023.06.05.543790

**Authors:** Leonidas-Romanos Davranoglou, Alban Lauka, Aris Aristodemou, Zoltán Maróti, Gjergj Bojaxhi, Ardian Muhaj, Ilia Mikerezi, David Wesolowski, Brian D. Joseph, Alexandros Heraclides

## Abstract

The origins of the Albanian people have long been debated, as they first appear in historical records in the 11^th^ century CE, and their language is not closely related to any surviving Indo-European branches. To elucidate the origins of the Albanians, we analysed over 6,000 ancient West Eurasian genomes and 74 newly sequenced present-day ethnic Albanians. We detect remarkable continuity of West Balkan Late Bronze and Iron Age ancestry in Albania during the Early Medieval period, a pattern distinct from neighbouring Balkan regions. Utilising a wide range of population genetics methods, including an enhanced protocol to detect identity-by-descent (IBD) segments between ancient and present-day individuals, we reveal that present-day Albanians predominantly descend from Albania’s Early Medieval inhabitants, who were present in Albania as early as 800-900 CE, preceding their historical attestation. Additionally, we observe geographically structured admixture with Medieval East European-related groups, averaging 10-20% across Albanians. Our findings provide unprecedented insights into the historical and demographic processes shaping present-day Albanians and locates the origins of this population into the Central-Western Balkans.

## Introduction

During the Iron Age (1100 BCE–150 CE), the Balkans were characterised by remarkable cultural and linguistic heterogeneity^1–6^. In the western Balkans, “Celtic” cultures interacted for centuries with groups referred to as the “Illyrians” and “Dalmatians“^7,8^. Deeper in the Balkan heartland, populations named by classical authors as “Dacians”, “Dardanians”, “Moesians”, and “Paeonians”^1,7–9^ bordered nomadic cultures from the Pontic-Kazakh steppe known as the “Scythians”^10^, while the southeastern part of the peninsula was inhabited by “Thracians” and “Greeks”^2,11^. Balkan peoples also expanded beyond the confines of the peninsula, with “Messapians” migrating to southeast Italy at least since 600 BCE^12^. The linguistic and cultural diversity of the Balkans was considerably homogenised during the Hellenistic^2,9^ and Roman^7–9^ eras, and especially following the Migration Period, when Germanic and Slavic-speaking groups massively settled in the region^2,3,13,14^. These events ultimately led to the extinction of all palaeo-Balkan languages except Greek and Albanian. The latter is a distinct branch within the Indo-European language family, inasmuch as it is attested only from the 15^th^ century CE, with the first substantial text not coming until the mid-16^th^ century, its prehistory and its possible relations with other branches and other languages within the family are somewhat enigmatic^2,15,16^.

Tracing the origins of the Albanians is challenging, as none of the available historical sources mention an Albanian-speaking population from the territory of present-day Albania during the transition from classical antiquity to Medieval times (500-1000 CE)^17–19^. Incursions and settlements by speakers of Slavic languages in what is now Albania are mentioned in the 7^th^-8^th^ centuries CE^18,20^, and the past presence of such population groups is mirrored by numerous Slavic place names^21^. The southwestern part of present-day Albania had a loose association with the Eastern Roman Empire, but the language(s) and emic identities of its inhabitants are shrouded in mystery^22,23^. The urbanised Medieval populations of northwest present-day Albania, referred to by contemporary historians as the Romani (Ῥωμᾶνοι)^17^, may have spoken a variant of vulgar Latin known as West Balkan Romance^20,24,25^ that persisted at least until the 13^th^ century CE^26^. It is only in the 11^th^ century CE that Albanians appear in the historical record, while the earliest surviving written documentation of their language, a one-line baptismal formula, dates to 1462 CE^27^.

However, it is evident that the ancestors of present-day Albanians lived in Albania much earlier than historical sources suggest. Linguistic hypotheses propose that an Albanian-speaking group may have inhabited high-altitude areas spanning present-day northern Albania, Kosovo, southeastern Serbia, and northern North Macedonia at least since the 5^th^ century CE^28,29^. Furthermore, the split between the two main dialects of modern Albanian – Gheg in the north and Tosk in the south (separated by the Shkumbin River in central Albania) – likely took place prior to or during the early phases of Albanian-Slavic contact^29,30^.

The affinities of the Albanian language itself have also been hotly debated, yet no definitive conclusions have been drawn. The most prominent, mutually exclusive hypotheses can be divided into those arguing for a local West Balkan origin from an Illyrian^30^ or Messapic background^16,31^ (which may or may not have been distinct languages^7,31,32^), and those proposing an origin from adjacent Daco-Moesian-Thracian groups^2,16,33^, whose speakers entered Albania after 400 CE^20,28,32,34^. The validity of these hypotheses is hard to test, as these ancient languages are poorly recorded, being known only from fragmentary inscriptions, toponyms, and a handful of historical sources^2,7,35^. Furthermore, all of the ethnonyms of ancient Balkan peoples, such as “Illyrian” and “Thracian”, are likely artificial constructs coined by both ancient and present-day authors^36^, and may encompass diverse groups of related languages with unclear geographical boundaries, mutual intelligibility, and emic identities of their speakers^8,9,32^. Recent linguistic hypotheses propose a sister-group relationship of Albanian to Greek^15,37,38^, which firmly places the origin of the language in the Balkans but does not pinpoint the location of the proto-Albanian homeland within the peninsula and its potential affiliation to historically attested populations.

Due to the challenges associated with linking archaeological, literary, and linguistic evidence, an archaeogenetic approach is critical in terms of providing novel insights into the origin of the Albanians, their biological relationships to ancient people, and the affinities of their language. Indeed, recent years have witnessed a surge in the palaeogenomic sampling of the Balkan peninsula^3–6,11,13^, which identified three major events that shaped the ancestry of modern Balkan peoples. During the early Roman era (1-250 CE), Anatolian and Near Eastern-related ancestry became pervasive throughout the Empire, including the Balkans^3,6,39,40^. Between the 3^d^ and 4^th^ centuries CE, the Balkans received an influx of individuals from the frontier regions, mainly Germanic and nomadic steppe groups^3^. The third and most significant migration occurred by 700 CE, with Slavic-speaking groups settling throughout the Balkans, and contributing 30-60% of Eastern European-related ancestry to present-day Balkan peoples^3^.

A previous study shows that the ancestors of Albanians participated in the demographic processes described above, as they derive approximately 25% of their ancestry from a local Bronze Age-Iron Age source, with additional contributions of 45% from Anatolian-related and 30% from Eastern European-related proxies, respectively^3^. However, this broad approach does not allow the identification of the likeliest Roman, Medieval, and Post-Medieval populations who contributed to the ancestry of present-day Albanians, while the ancestry of the pre-Roman samples from Albania was not examined. Additionally, that study relies on a limited set of present-day Albanian samples (n = 6), sequenced in Tirana and of unknown provenance, potentially failing to capture the full range of Albanian genetic diversity. Another study based entirely on present-day populations suggests that Albanians descend from a small population that experienced a major bottleneck 1,500 years ago^41^. Despite significant advances in the use of genetics to study Balkan history, the origins of Albanians remain a complex and debated question.

To elucidate the origins of the Albanians, we conduct a 4,000-year palaeogenomic transect of the inhabitants of the territory of present-day Albania and the Balkans. We analysed more than 6,000 previously published ancient genomes from western Eurasia (including 22 from Albania) (Fig. 1A) and 74 newly sequenced present-day Albanians, covering all dialectal groups (Fig. 1B). We interrogate this dataset using group-based analyses (f-statistics, *qpAdm*-based admixture modelling^42,43^, and Distribution of Ancestry Tracts of Evolutionary Signals - DATES^44^), principal components analysis (PCA), and large-scale algorithms that quantify human mobility^45^. Furthermore, we apply an optimised version^46^ of a previously published^47^ protocol to study identity-by-descent (IBD) segments – large genomic tracts co-inherited from recent common ancestors. Additionally, we analyse publicly available uniparental data from over 4,000 ancient and present-day Balkan samples, in order to trace migrations into and out of present-day Albania. Our multifaceted approach allows us to trace key demographic events that shaped the modern Albanian genome, and to identify the likely geographic and temporal origins of Albanians.

**Fig. 1.**
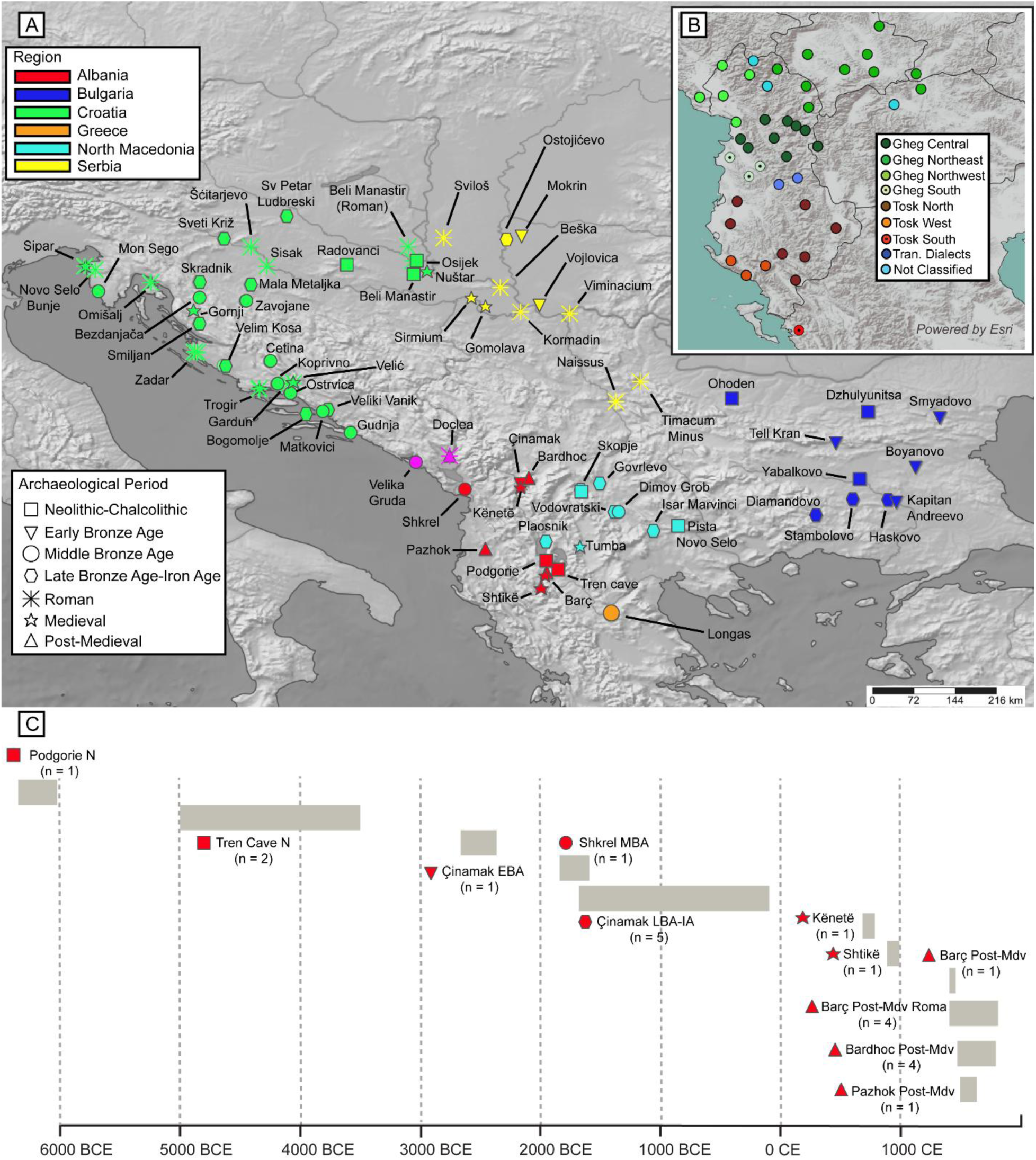
Geographical distribution and dating of the ancient and present-day Balkan populations examined. A) Archaeological sites annotated by period. B) Inset showing locations of newly sequenced present-day Albanian subpopulations. Note that the total number of sampling sites (43) does not correspond to the total number of samples (74), as multiple individuals were sequenced from some locations. The inset map of Albania was created with QGIS 3.40.0.^48^ using Esri World Terrain Base Map, ArcGIS Online^49^. C) Analysed ancient individuals from the territory of present-day Albania arranged by archaeological site and date (radiocarbon or archaeological chronology).

## Results

### Estimating the arrival of Indo-European languages in Albania

To elucidate the origins of the Albanians, we aimed to capture the timeframe during which their linguistic ancestors arrived in the Balkans. During the Early Bronze Age (ca. 2600-1800 BCE), populations related to the putatively Indo-European-speaking Yamnaya steppe pastoralists arrived in the Balkans, significantly altering the region’s linguistic, cultural and genetic landscape^4,11,50–53^. Only a single male individual (I14689) is known from the territory of modern Albania during this transitional period, specifically from the northeastern site of Çinamak, dating to 2663-2472 BCE^6^ (Table S1). To visualise the genetic affinities of this individual, we performed principal components analysis (PCA), where we projected previously published ancient individuals from the Balkans and adjacent regions onto present day West Eurasian populations genotyped with the Human Origins array^54^ (Fig. S1).

Mirroring the findings of a previous study^6^, the PCA indicates remarkable ancestry shifts in the Balkans during the Early Bronze Age. While Neolithic-Chalcolithic Balkan populations cluster on the bottom left corner of PC2 with Early European Farmers (EEF), Early Bronze Age populations show a significant genetic shift, plotting towards the Yamnaya on the top left corner of PC2 (Fig. 2). The Albania_Çinamak_EBA individual clusters closely with the Yamnaya (Fig. 2), with our *qpAdm* and unsupervised ADMIXTURE tests both indicating that approximately 70% of his ancestry derived from the Pontic-Caspian steppe, and the remainder from presumably local EEF populations (Figs. S2-3; Table S2). This individual carried the Y-chromosome haplogroup R1b-M269^6^, a primary indicator of paternal ancestry from the Pontic-Caspian steppe^6,52,53^. Our further analysis using *qpAdm*, f3-statistics, and f4-statistics, suggests that steppe-derived admixture in Albania Çinamak EBA likely derives from largely unadmixed Yamnaya groups rather than EEF- and-European hunter-gatherer-admixed Corded Ware populations (Figs. S5A, S6A; Tables S4-S6). This suggests that the first wave of Indo-European-related ancestry arrived in Albania in largely unadmixed form directly from the Pontic-Caspian steppe. To estimate the approximate admixture date between the EEF-related and steppe-related ancestors of Albania_Çinamak_EBA, we used the DATES method^44^. Our analysis yields an admixture date of 4 ± 2.5 generations before the age of Albania_Çinamak_EBA (Table S7), suggesting that his ancestors were recent arrivals from the steppe, possibly at ca. 2700 BCE. Indeed, contemporary archaeological artefacts from tumuli in Shkodër, located 100 km west of Çinamak, showed cultural links with the northern Adriatic and present-day Hungary^55^, suggesting possible routes for the introduction of steppe-related ancestry into Albania. While the relationship between the languages spoken in EBA Albania and modern Albanian remains unknown, this evidence offers a plausible timeframe for the introduction of Indo-European languages into the region.

**Fig. 2.**
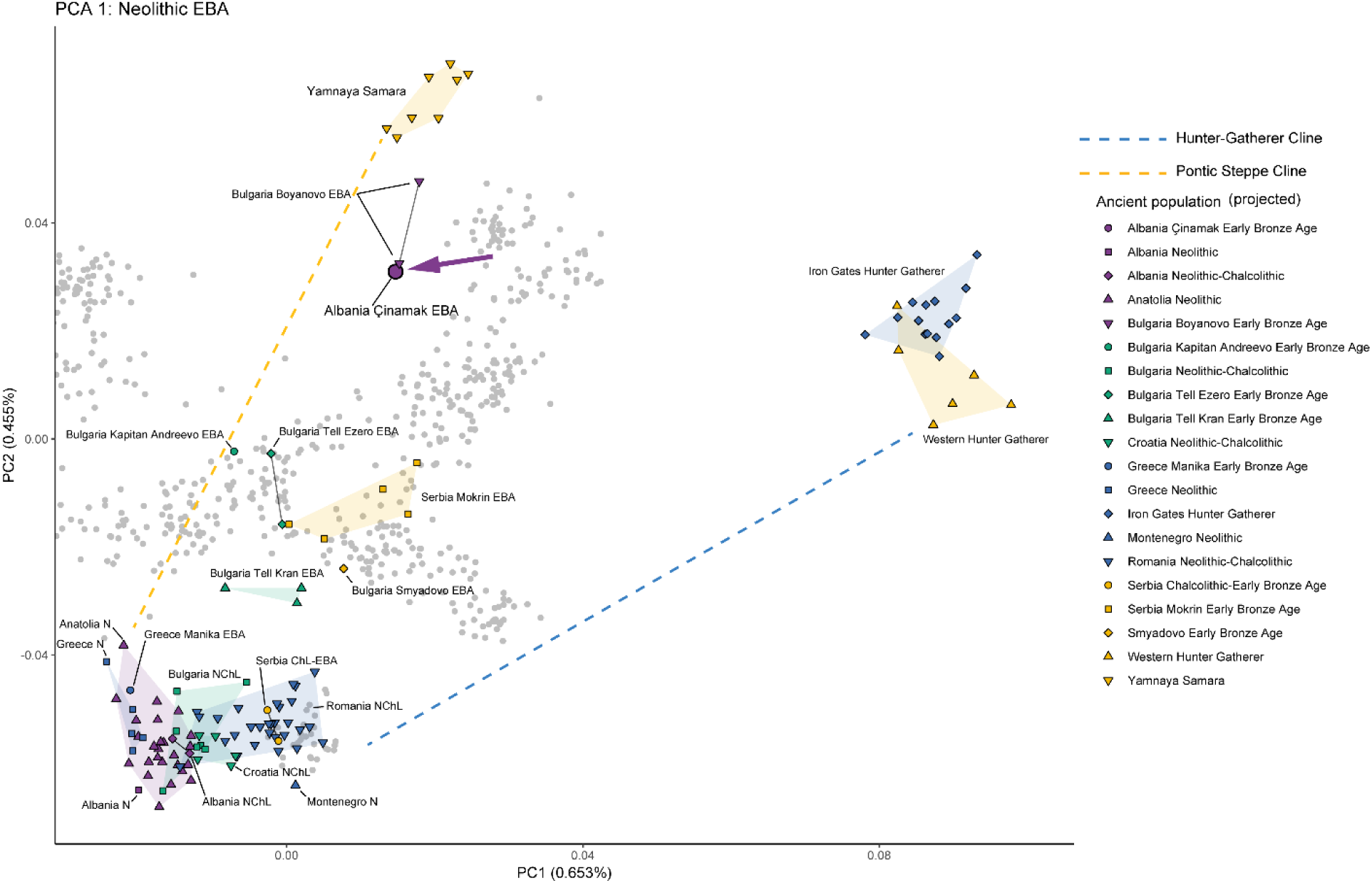
PCA of present-day West Eurasian samples (grey points) with projection of ancient Neolithic-Early Bronze Age individuals from the Balkans. The purple arrow shows the position of the Early Bronze Age sample from Çinamak in Albania. Dashed lines indicate the two poles of observed ancestry shifts from the Neolithic onwards – one generated by the assimilation of hunter-gatherers (blue line), and another by admixture with incoming Yamnaya-related populations from the Pontic-Caspian steppe (yellow line). Samples clustering within the PCA space enclosed by HG, EEF, and Yamnaya derive variable amounts of ancestry from these populations.

### The origins of the classical era inhabitants of Albania

To investigate the ancestry of “Illyrians”, who were the predominant cultural group in classical era Albania^8,30,35^, we examined the projections of Middle-Late Bronze Age (BA) and Iron Age (IA) Balkan populations upon the principal components presented above (Fig. 2), including BA-IA individuals from Çinamak (Fig. 3). During this period, the ancestry of Balkan populations had become less heterogenous, exhibiting a north-to-south cline on the PCA that broadly reflects geography (Fig. 3A). Accordingly, the Albania_BA_IA individuals from Çinamak, an “Illyrian” region^1,7,8^, overlap with contemporary populations from Croatia, Montenegro, North Macedonia and northern Greece on PC1 and PC2 (Fig. 3B). To test whether this genetic shift is attributable to changes of steppe versus EEF ancestry, we ran *qpAdm* models using Mesolithic-Neolithic source populations (Table S3). These analyses indicate homogenisation of EEF ancestry across the sampled Central-West Balkans sites and one site in northern Greece (with inhabitants of uncertain linguistic affiliation) compared with the preceding EBA, as most samples derive around 60% of their ancestry from EEFs, 0-5% from Neolithic Iran (Iran_N), and 30-40% from steppe populations (Table S3). In the southern end of the Balkan genetic cline, populations of central-southern Greece and Bulgaria have considerably higher EEF (75-80%) and Iran_N (5-10%) ancestry, and a much lower proportion of steppe ancestry (15-20%) (Table S2). It should be noted that a previously published^6^ *qpAdm* model that utilised different source populations found similar ancestry proportions for all abovementioned groups.

**Fig. 3.**
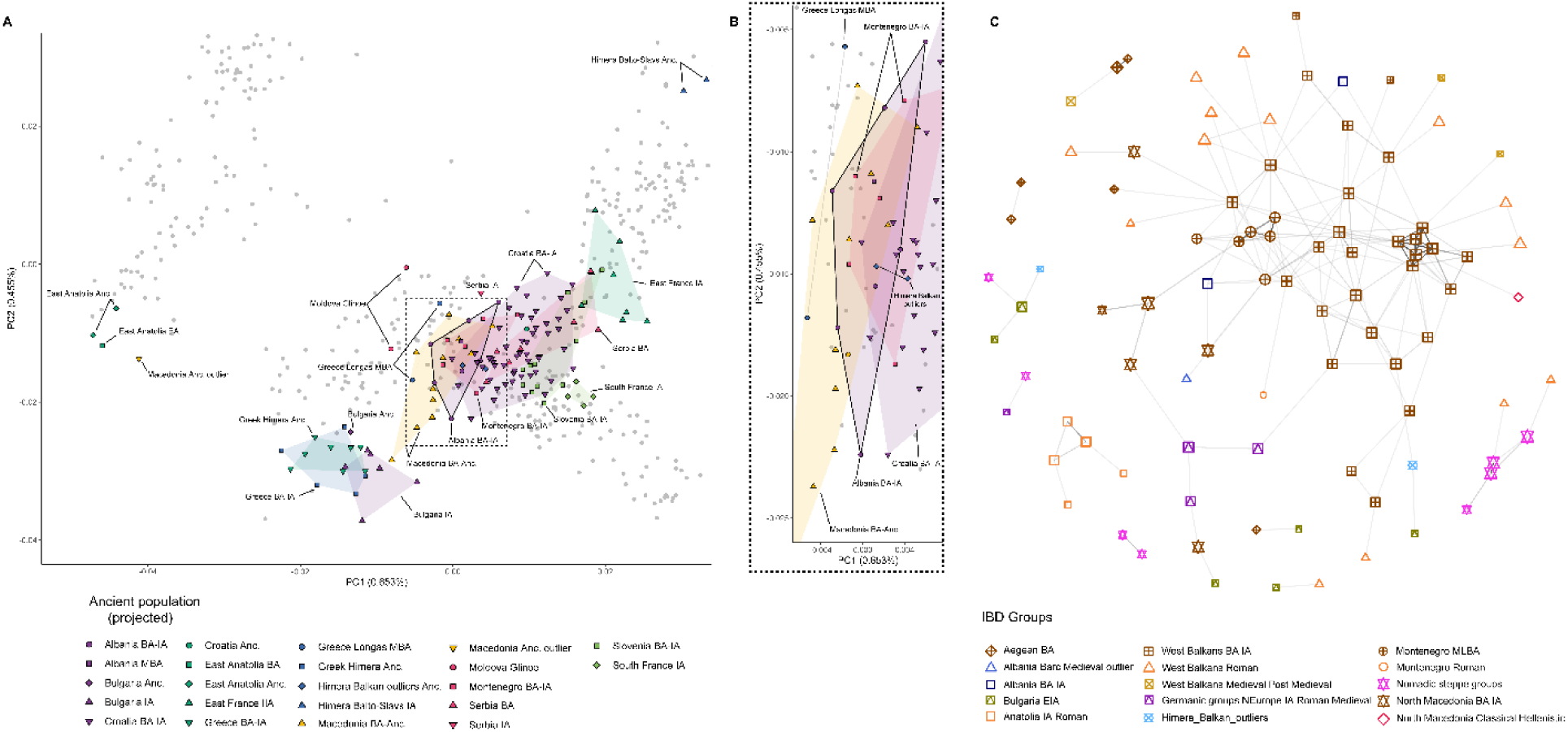
PCA of present-day West Eurasian samples (grey circles) with projection of ancient Bronze Age and Iron Age individuals from the Balkans. A) PCA of all tested populations, showing a north-to-south cline that mirrors geography. B) Detail of region enclosed in the dotted rectangle in panel A, showing the clustering of populations from Albania (enclosed in solid line). C) Intergroup Identity-by-Descent (IBD) sharing among selected ancient genomes from West Eurasia, focusing on matching patterns at 8 cM in Bronze Age and Iron Age individuals. The genomes are categorized by geographical regions and archaeological periods (refer to Table S2 for detailed information). Lines indicate the presence of a shared IBD segment.

However, the genetic affinities of Albania_BA_IA to contemporary populations have not been examined previously. Using a *qpAdm* model that tested a battery of West-Central Balkan BA-IA, Greece MBA-BA, and Bulgaria_EIA-related base set of references, we recover the Albania_BA_IA group as cladal with the adjacent Montenegro_MLBA and North_Macedonia_BA populations (Table S4), an affinity further supported by *f3* and *f4*-statistics (Fig. S5C, S6; Tables S5-S6) and Mobest analysis (Fig. S7), which employs an algorithm for spatiotemporal mapping of genetic profiles using bulk aDNA data^45^. To test the strength of our *qpAdm* models, and to investigate the recent genealogical connections of Albania_BA_IA, we performed IBD analyses using an optimised protocol of ancIBD^46^ (Fig. 3C; Tables S8-S17). We selected and imputed 330 ancient genomes from Western Eurasia (ranging from the BA to the Post-Medieval period and focusing on the Balkans), implementing an approach that enables the identification of segments of at least 8 centimorgans (cM) in length (see details in Methods). We reveal genealogical connections of Albania_BA_IA with both northern (Mon Sego) and southern (Gudnja cave, Veliki Vanik) populations of Croatia_BA, Montenegro_MLBA, and eastern North_Macedonia_IA (Valandovo), with shared genomic segments of ca. 8-10 cM in length (Fig. 3C). These findings demonstrate that the population of Albania_BA_IA was a core part of the “Illyrian” world that spanned the Adriatic^8^, but also maintained connections with groups from North Macedonia known as “Paeonians”, who spoke an Indo-European language of uncertain affinities^2,9^. The absence of intragroup IBD-sharing in Albania_BA_IA may stem from the low sample size and large chronological range of the examined individuals (1700-400 BCE; Table S1).

### Albania as a refugium of BA-IA West Balkan ancestry during the Medieval period

Previous studies have shown that the genetic landscape of the Roman Empire transformed significantly during the Imperial period due to large-scale immigration from the Eastern provinces, introducing post-Neolithic West Asian ancestry throughout the Empire^3,13,39,40^. More profound ancestry shifts occurred during the Migration Period (approximately 3^d^-8^th^ centuries CE) as initially Germanic and nomadic steppe groups, followed by Slavic-speaking groups migrated and settled in various parts of the Roman Empire^3,14,39,40^. These migrations led to significant changes in the genetic makeup of the Balkans^3^. This is evident in the PCA of Roman and Post-Roman samples (Fig. 4A), where nearly all populations have shifted from their Bronze Age and Iron Age positions towards West Asian and Eastern European clusters, forming apparent Anatolian and East European clines, respectively.

**Fig. 4.**
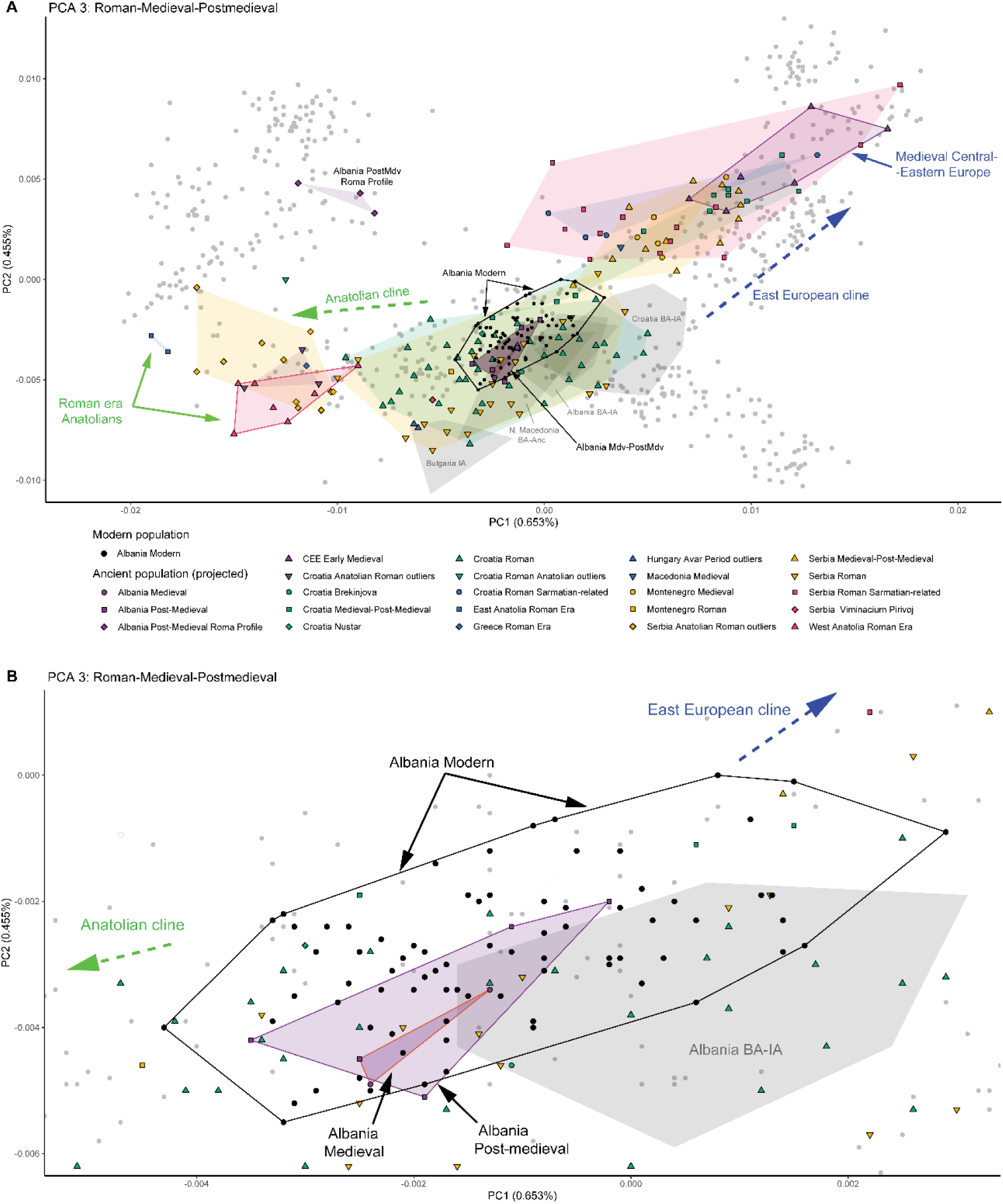
PCA of present-day West Eurasian samples (grey circles) with projection of ancient Roman, Medieval, and post-Medieval individuals from the Balkans. A) PCA of tested populations, highlighting the shift in the PCA coordinates of Roman and post-Roman groups towards the Anatolian (green dashed line) and East European (blue dashed line) clines, compared to their counterparts from the same regions in the preceding Bronze Age and Iron Age (faded grey polygons). B) Detail of region in panel A, showing the tight clustering of populations from Albania over the past 3,000 years, including 74 newly sequenced present-day Albanians (black polygon). For simplicity, Roman-era populations within and close to this cluster have been removed and can be seen in the previous panel.

To quantify the impact of Eastern European migrations into the Balkans, we used *qpAdm* tests with source populations dating to the Iron Age, Bronze Age and the Roman era, where CEE_Medieval (Early Medieval samples from western Hungary, Czechia, eastern Austria, and western Slovakia) served as a proxy for unadmixed Eastern European-related ancestry (following^3^). Our *qpAdm* models uncovered remarkably high Eastern European-related ancestry in the late Roman and Post-Roman populations of Croatia (75-86%), Montenegro and Serbia (ca. 65%), and North Macedonia (ca. 40%) (Table S4). These admixture proportions were nearly identical to those proposed by a study on present-day Slavic-speaking populations using different methods, which identified a core Slavic component in present-day South Slavs ranging from 55-70%, with the remainder of their ancestry originating from the pre-Slavic inhabitants of the Balkans ^56^.

We next sought to model the ancestry of the two available samples from Medieval Albania, which originate from South-Eastern (Shtikë, 889-989 calCE) and North-Eastern (Kënetë, 773-885 calCE) Albania (referred to as Albania_Medieval). These samples showed only a minor shift in their PC position from the Bronze Age-Iron Age (BA-IA) to the Migration Period (Fig. 4A, B), suggesting large-scale genetic continuity for over 2,500-3,000 years. We ran a *qpAdm* model that compared a local Albania_BA_IA source population against an Eastern Balkan proxy (Bulgaria_EIA), and populations with Southeast Anatolian ancestry (Table S4). The results indicated that 68-84% of Albania_Medieval’s ancestry came from an Albania_BA_IA-related population, with additional admixture from Anatolian (16-32%) or Bulgaria_EIA-related (11-32%) sources (Table S4). The Anatolian-related ancestry ranged from 16-21% with a Southeast Anatolian proxy and up to 32% with West Anatolian Romans or Anatolian-admixed Roman Balkan samples (Table S4). It is possible that Anatolian and Bulgaria_EIA-related sources were conflated by *qpAdm*, due to both deriving most of their ancestry from EEFs (Table S3).

The Bulgaria-EIA-related ancestry showed lower p-values (p = 0.067) and higher standard errors (0.101) which may indicate statistical uncertainty but cannot be used to reject this model^42^. Southeast Balkan-related ancestry could not be excluded, as *f4*-statistics of the form *f4(Cameroon_SMA, palaeo-Balkan sources; Albania_Medieval, Anatolian-admixed samples)* indicated very high allele sharing with Balkan populations when compared to Anatolian-admixed individuals (Fig. S6; Table S6). In turn, *f4*-statistics suggested that Bulgaria_EIA-related sources from the Roman period (Roman_Balkans_Thracian_profile) exhibited slightly higher allele sharing with Albania_Medieval when compared against West Balkan individuals dating from the Bronze Age to the Roman period (Fig. S6; Table S6).

To formally exclude the presence of East European-related ancestry in Albania_Medieval, we further tested its ancestry using a *qpAdm* model that rotated among Roman era West and East Balkan source populations, East European (CEE_Medieval), and Anatolian (West_Anatolia_Roman) proxies. This analysis recovered Albania_Medieval once more as 100% Roman West Balkan (Table S4), showing that the ancestors of this population were largely shielded from the demographic upheavals of the Migration Period.

### The formation of the present-day Albanian gene pool

We next sought to investigate the ancestry of post-Medieval samples from central (Pazhok, 1527-1660 calCE) and north-eastern (Bardhoc, 1400-1700 calCE) Albania. The place-name Bardhoc, of Albanian origin^34^, appears in contemporaneous Ottoman registers^57,58^, indicating the individuals might have been Albanian speakers. Remarkably, post-Medieval Bardhoc individuals (Albania_Bardhoc_Post_Medieval) cluster with Medieval samples from Kënetë and Shtikë (Fig. 4A, B), except for one outlier clustering with the Pazhok sample, both of which are shifted towards the East European cline.

We used a two-way *qpAdm* model to examine if the PCA pattern reflects shared ancestry. Albania_Medieval served as a local source population, with other source populations serving as proxies for unadmixed East European-related ancestry (CEE_Medieval), and palaeo-Balkan-admixed South Slavic-related populations from the Medieval and Post-Medieval periods (Montenegro_Medieval, Croatia_Medieval_Post_Medieval, Serbia_Medieval_Post_Medieval) (Table S4).

One-way models for Albania_Bardhoc_Post_Medieval using Albania_Medieval as source populations received high statistical support (Fig. 4A, B; Table S4), suggesting continuity from the preceding Medieval period. However, a one-way *qpAdm* model for the East European-shifted outliers from Bardhoc and Pazhok was rejected (Table S4), indicating more complex ancestry in these individuals. These outliers were instead recovered as deriving their ancestry from a two-way mixture between a local source (Albania_Medieval) and unadmixed (ca. 21%) or admixed (ca. 25-32%) East European-related groups (Table S4). The individual from Pazhok also carried an East European-derived Y-DNA lineage, R1a-CTS1211>BY33436. The presence of East European ancestry in two early modern samples from different regions in Albania suggests that it was already being diffused among Albanians during the Middle Ages, and that currently unsampled adjacent subpopulations may have had varying levels of West Balkan_IA, West Anatolian and East European ancestry components, which became homogenised in later periods.

Using the insights gained from the analysis of the Post-Medieval samples from Bardhoc and Pazhok, we next attempted to model the ancestry of 74 newly-sequenced present-day Albanians, covering all major dialectal groups. As can be seen from the PCA (Fig. 4A, B), present-day Albanians plot with or adjacent to the Medieval and Post-Medieval inhabitants of the country, with some individuals displaying varying degrees of affinity towards the East European cline.

For our *qpAdm* analyses on present-day Albanians, we aimed to capture West Balkan genetic variation beyond what the two Medieval samples from Shtikë and Kënetë provide, and to maximise the number of SNPs used. Thus, we created a new category called *West_Balkan_Roman_Medieval*, which includes Albania_Medieval and Montenegro_Doclea_Roman. These geographically adjacent individuals share a similar PCA position (Fig. 4A) and nearly identical proportions of West Balkan Iron Age ancestry (Table S4). Additionally, we included a Roman-era Central Balkan proxy (Serbia_Roman_Naissus) from the city of Naissus (an area likely influenced by the linguistic ancestors of Albanians^59,60^), along with various East European-related proxies (CEE_Medieval, Montenegro_Medieval, Croatia_Medieval_Post_Medieval).

Our analysis indicated remarkable continuity of present-day Albanians from the ancient West Balkans, as many sampled individuals (34/74, or 46%) were recovered as cladal with West_Balkan_Roman_Medieval (Table S4). We also found significant variation in East European-related admixture among present-day Albanians (Fig. 5). Depending on the proxy used – either largely unadmixed (CEE_Medieval) or paleo-Balkan-admixed (Montenegro_Medieval, Croatia_Medieval_Post_Medieval) – averages of present-day Albanian subpopulations showed 4-16% or 8-32% of this ancestry, respectively (Fig. 5; Table S4). However, some individuals from TN and two outliers from GNW exhibited 23-50% CEE_Medieval or 36-73% Montenegro_Medieval ancestry, respectively. When we merged all present-day Albanian subpopulations into a single group, the average proportion of East European-related ancestry was 12% or 23%, depending on whether CEE_Medieval or Montenegro_Medieval was used as a proxy, respectively (Table S4). Based on *f4*-statistics, Montenegro_Medieval displayed excess allele sharing with present-day Albanians (Fig. S6), although this could be attributed to the significant Roman West Balkan ancestry of this population (Tables S4, S6).

**Fig. 5.**
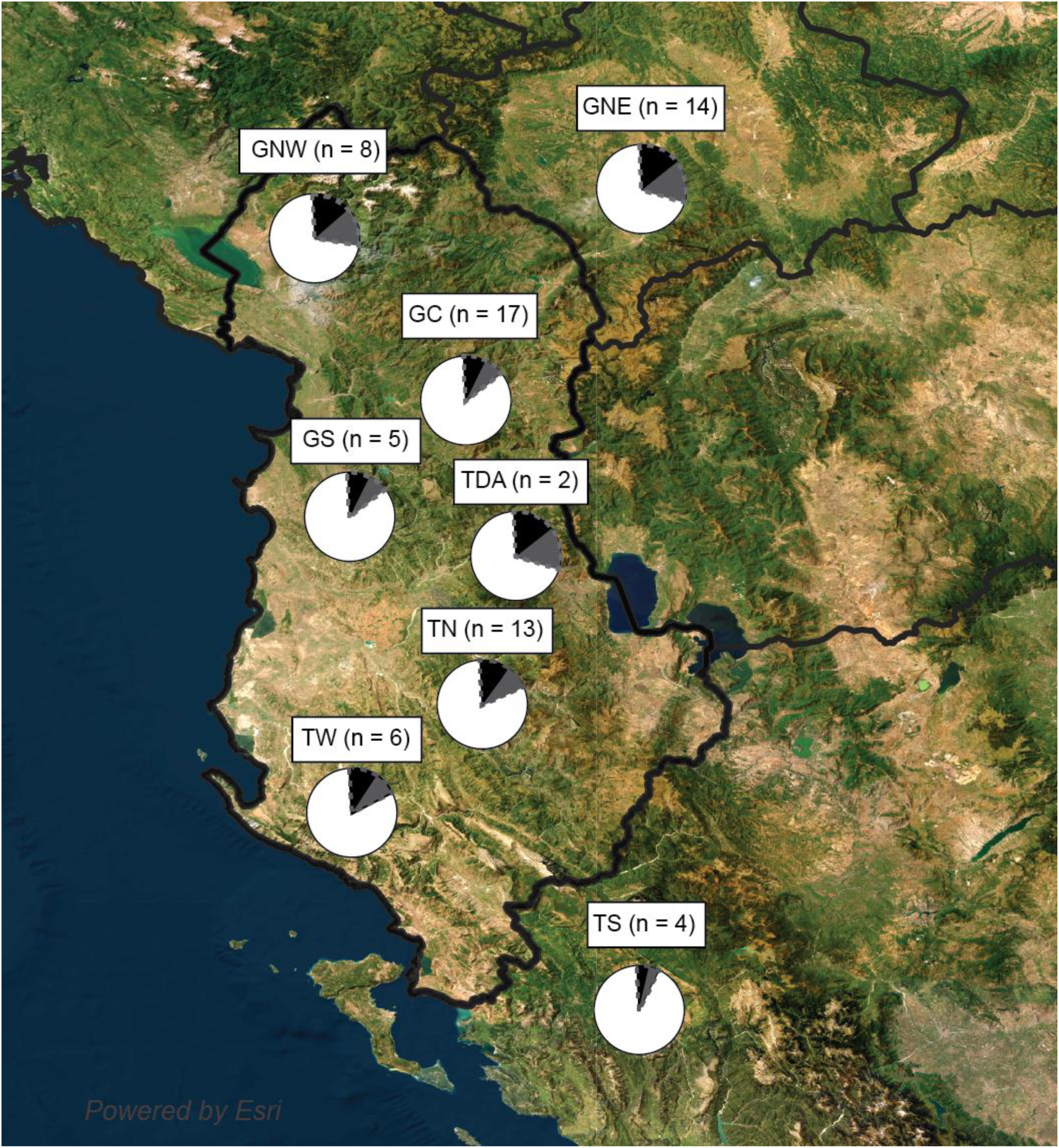
Ancestry proportions of present-day Albanians, utilising two proxies for East European-related ancestry: CEE_Medieval (black) and Montenegro_Medieval (dashed grey). Each proxy represents a lower and upper proportion of East European ancestry, respectively. The inset map of Albania was created with QGIS 3.40.0.^48^ using Esri Satellite (ArcGIS/World_Imagery)^61^.

Based on PCA and *qpAdm* models, we showed that present-day Albanians descend from earlier Medieval groups in Albania, which derived most of their ancestry from Late Bronze and Iron Age western Balkan individuals. Additionally, despite its relative isolation, post-Medieval Albania had a diverse cultural landscape, as evidenced by three individuals from Barç who plotted far from other Balkan populations on the PCA (Fig. 5B). Previously thought to be of Central Asian origin^13^, our *qpAdm* and *ADMIXTURE* analyses revealed that these individuals had South Asian ancestry, likely from the Roma people^62^ (Fig. S2; Table S4). This was further supported by Y-chromosome haplogroup J2a-Y18404 and mitochondrial haplogroup U3b1 found in these individuals, markers commonly associated with Roma populations. These lineages likely entered the founding Roma population during their journey in the Caucasus^63,64^. To our knowledge, this represents the first aDNA record of Roma people and the first line of evidence for their entry into Europe.

### The dynamics of East European-related admixture among the ancestors of present-day Albanians

The geography and chronology of the Albanian-Slavic contact has remained an open question^29,30,65^. The genetic results reported here are congruent with the findings of historical linguistics, showing the highest proportions of East European-related ancestry along the Albanian-Montenegrin border (GNW, 14-28%), northeast Albania and Kosovo (GNE, 16-30%), and near Lake Ohrid (TDA, 16-32%; individuals TN2, TN7, 23-46%) (Fig. 5; Table S4). These areas coincide with the suggested principal locations of the longest Albanian-South Slavic linguistic contact^21,66,67^, while Slavic toponymy also peaks in the northeast and especially the southeast of the country^68^.

Using DATES, we estimated the timing of East European-related admixture in the ancestors of present-day Albanians to 500-1400 years before present (Fig. S9; Table S7). These findings align with hypotheses on Albanian-Slavic linguistic contact, which postulate limited interactions during the Migration Period, increasing significantly from the 11^th^ century CE to the Ottoman period^21,65–68^. However, the dissemination of East European ancestry in Albanians likely involved South Slavic-speaking groups, as well as subsequent mixing between unadmixed Albanian groups and those with high levels of East European-related ancestry. The complexity of these admixture layers poses challenges in accurately capturing them using DATES and may account for the relatively wide range of admixture dates in our model (Fig. S9; Table S7).

A significant knowledge gap on the introduction of East European-related ancestry in the ancestors of Albanians involves its social and demographic patterns of ancestry transmission. It is not known whether individuals with elevated East European-related ancestry who mixed with Albanians were primarily male, female, or had an even sex ratio. To test for sex bias, we compared ancestry proportions on the X-chromosome with autosomes using *qpAdm*. The results were inconclusive, likely due to limited X-chromosome markers (4.6k SNPs vs. 1240k SNPs on autosomes) and high standard errors (S1 Text; Table S4).

To overcome this limitation, we examined Y-chromosome haplogroup frequencies in 2,272 present-day ethnic Albanian men using publicly available datasets^69–73^ (Table S18). We found that Albanians derive ca. 19% of their paternal ancestry from Migration Period-associated Y-chromosome haplogroups (R1a-M417, I2a-M423, I1-M253; Fig. S10), in contrast to present-day South Slavic peoples, where these lineages reach 36-70%^74–76^. However, Y-DNA haplogroup frequencies may be prone to genetic drift due to the Y-chromosome’s reduced effective population size compared to autosomes (NeY-chr = 1/4 NeAutosomes)^77,78^.

For this reason, we also examined mtDNA haplogroups in 375 present-day Albanians (Table S19, using^64,79^). Data from mtDNA haplogroups may be of lower resolution for population genetics compared to Y-chromosome haplogroups due to higher mutation rates and smaller genome size, leading to backmutation and saturation^80^. Additionally, historical gene flow between Mediterranean and northeastern European populations^13,39,40^ has blurred the historical phylogeography of many mtDNA lineages.

Considering these factors, we identified putative Migration Period-related mtDNA haplogroups in 16% of the present-day Albanian population (Table S19). This may be an underestimate, as incoming Migration Period populations absorbed Balkan Romans early on^3^, suggesting that assimilation of palaeo-Balkan and West Asian mtDNA haplogroups may partially mask the East European-related autosomal signal. Indeed, in East European-admixed Balkan samples from the Medieval and Post-Medieval periods, we find 26% and 38% frequencies of palaeo-Balkan and Migration Period-related mtDNA haplogroups, respectively (Table S19), contrasting the larger autosomal (40-86%) and Y-DNA (64%) contributions of East European-related ancestry (Tables S4, S19).

Overall, autosomal, Y-DNA, and mtDNA data indicated strikingly similar levels of Migration Period-related ancestry in present-day Albanians (ca. 10-20%, 19%, and 16%, respectively). This suggests that East European ancestry diffusion in Albanians occurred roughly evenly through both sexes and at much lower rates than those observed in surrounding present-day Balkan populations.

Our *qpAdm* analyses revealed the long-term persistence of West palaeo-Balkan ancestry in Albanian populations from Medieval times to the present day. However, *qpAdm* cannot ascertain whether West palaeo-Balkan ancestry in Albania originated from local descendants or related groups migrating from adjacent regions such as the northern Adriatic coast or the Balkan interior.

To investigate genealogical connections from the Bronze Age to the present day in Albania, we used an optimised version of ancIBD^46^ for Identity-by-Descent (IBD) analysis. This analysis included historically and geographically relevant samples from across West Eurasia. We used segments greater than 8 cM to identify more recent IBD relationships (Fig. 6A), while we employed smaller segments (5-6 cM) to locate deeper intra- and interpopulation relationships (Fig. 6B). We applied a rigorous filtering protocol to detect false positive matches (see Methods), which suggested that 5-6 cM segments are as informative as larger IBDs, with only slightly false positive matches (0.64% and 0.11%, respectively; Table S9).

**Fig. 6.**
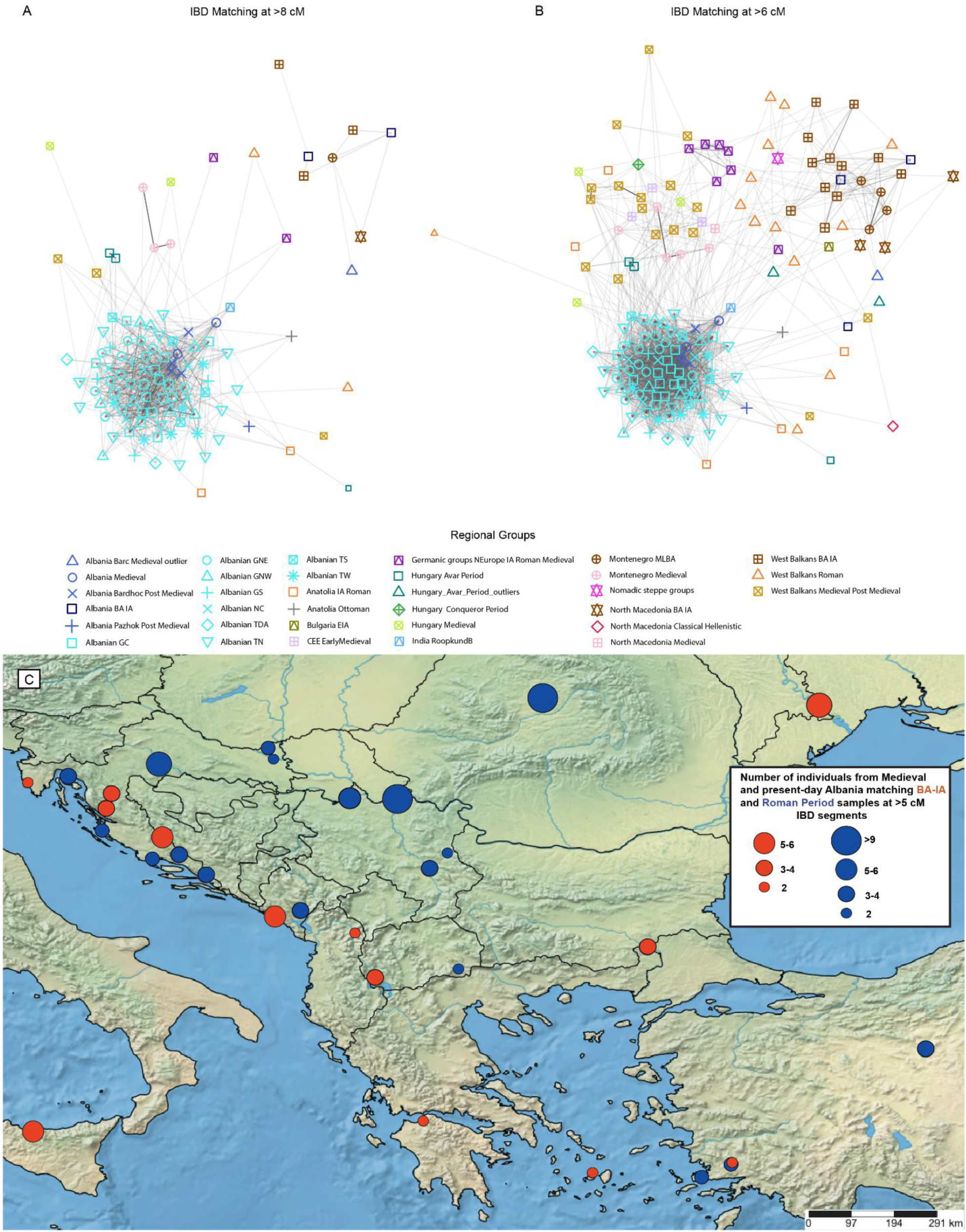
Identity-by-Descent (IBD) sharing among present-day Albanians and selected ancient genomes from West Eurasia. A) IBD matching patterns at >8 cM, showing close genealogical connections of Medieval, Post-Medieval, and present-day individuals from Albania. B) IBD matching patterns at >6 cM, revealing deep-time links of individuals from Albania with populations from the Balkans and West Eurasia more broadly. C) Number of Medieval and present-day individual having >5 cM IBD connections with populations from the Bronze Age to the Roman period.

In a remarkable case of interperiod connectivity, Albania_Medieval shared very large IBD segments with all Post-Medieval samples from Bardhoc (23 cM), as well as present-day Gheg (9.5-21 cM) and Tosk (9-11 cM) Albanians (Fig. 6A; Table S17). The two medieval samples also shared two IBD segments with each other, totaling close to 11cM (Table S20), indicating shared ancestry but not very recent common descent. Together with results from *qpAdm* and PCA (Fig. 4, Table S4), the IBD matching patterns provide conclusive evidence that all present-day Albanians directly descend from indigenous populations that lived in both north and south Albania over 1,200 years ago. This confirms linguistic hypotheses that suggested the presence of Albanian-speaking populations in southern Albania long before their first historical attestation^30,33^. Moreover, all populations from Albania, spanning Medieval times to the present day, share a genetic connection with a 17^th^-20^th^ century CE individual from Roopkund Lake in India, who clusters with present-day mainland Greeks^81^ (Fig. 6A; Table S16). This suggests that the individual from Roopkund Lake may have descended from Albanian-speaking communities in Greece, which persist to this day^15,71^, and may be the reason behind the high IBD-sharing between many present-day Greeks and Albanians^82^.

Crucially, the high affinity of Albania_Medieval to present-day Albanians was not the result of bias from the high coverage and large sample size of our modern population dataset. The two individuals comprising Albania_Medieval shared 69% and 77% of their IBD matches greater than 5 cM with Albania_Bardhoc_Post_Medieval and present-day Albanians (Table S20). In contrast, Late Roman and Early Medieval individuals from neighbouring regions (Montenegro, Serbia, and North Macedonia) were primarily linked with non-Albanian Balkan groups from earlier periods, with individuals from Medieval, Post-Medieval, and present-day Albania representing only 4-30% of their total matches (Table S20).

Similarly to Albania_Medieval, Post-Medieval samples showed IBD affinity to modern Albanians (Table S17). Even among the IBD matches of the outlier Pazhok sample, 82% include Albania_Bardhoc_Post_Medieval and present-day Albanians. The lower number of total matches of this sample suggests this individual could represent a profile within the Albanian genetic continuum which persisted in relative isolation, a possibility supported by a close IBD connection (>21 cM) with a present-day Albanian from nearby Tirana (Tables S17, S20).

### Links of the inhabitants of Albania to the broader Balkan sphere and the possible location of the proto-Albanian homeland

The genealogical links of Albania_Medieval with individuals from neighbouring Roman era Zadar in Croatia (ca. 11 cM), and Doclea in Medieval Montenegro (ca. 8.5 cM) (Fig. 6A; Table S16) suggest an origin close to the Adriatic^30^. However, Albania_Medieval is also genetically connected with roughly contemporaneous populations from more distant geographical locations, such as West Anatolian Romans, Gepids from Romania, and Avar-era commoners from Hungary (Fig. 6A; Table S16). The links with the individuals from Hungary suggest that possible trade networks between the inhabitants of Medieval Albania and the Avar Khaganate (based on archaeological artefacts^8,20,23^) may have also involved genetic exchange. Given that Albania_Medieval lacks East European and Central Asian-related ancestry (Table S4), it is likely that IBD matching with Avar era and Medieval Montenegrin populations is either due to a shared palaeoBalkan-Roman background, or that individuals from within the same mating pool as Albania_Medieval migrated and admixed with those populations.

Present-day Albanians display IBD matching patterns complementary to those of Albania_Medieval, specifically with an Iron Age Roman Period Scandinavian-related Germanic group from Poland, West Anatolian Romans, Medieval Hungarians, post-Medieval individuals from Serbia, and Ottomans who cluster with Balkan populations (Fig. 6A). Intriguingly, a Gheg individual from the coastal Kurbin area in Albania shares 8.3 cM with a Roman individual from Kormadin in Serbia (I27297) (Table S16), who had a mixed West and East Balkan profile (Table S4). This finding provides further support for linguistic theories suggesting a proto-Albanian homeland within the territory of present-day Albania, Kosovo, southern Serbia, and N. Macedonia^28,29^.

We identified genealogical connections by analyzing individuals from Medieval to present-day Albania with the Medieval-Post-Medieval population of Gornij in Croatia (7.5-11 cM) (Fig. 6A). A 16^th^-17^th^ century CE individual from Gornij (I35008) is characterized by mtDNA haplogroup T1a1l (Table S19). This haplogroup comprises 2% of the total maternal lineages of present-day Albanians and is almost exclusively associated with this ethnic group. It was also found in a Post-Medieval individual from Bardhoc (Table S19). Given that Albania_Medieval and Albania_Bardhoc_Post_Medieval individuals lack East European-related ancestry (Table S4), it is likely that a migration from an Albanian-related population introduced genealogical links and haplogroup T1a1l to Gornij, rather than vice versa. This connection may be related to the long-term presence of Albanian-speaking communities in various parts of present-day Croatia^27^, or perhaps earlier, historically unattested migrations. We interpret IBD matching between present-day Albanians and Medieval populations from Montenegro (Fig. 6A; Table S16) as likely stemming from shared palaeo-Balkan ancestry, as well as admixture during and after the Migration Period, as evidenced by the variable levels of East European-related ancestry in Albanians.

Using a filtered, lower IBD threshold (5-6 cM) (Tables S10-S11), we document relationships between populations from Albania spanning the Bronze Age to the Early Modern periods (Fig. 6B). We should stress that these segments may not represent true recent genealogical connections, but rather ancient population structuring that could help elucidate the deep-time ancestors of present-day Albanians. Remarkably, both Medieval and present-day inhabitants of Albania exhibit IBD matches with various Bronze and Iron Age populations from the West (Croatia and Montenegro, totaling 29 matches), Central (North Macedonia, 4 matches), and Eastern Balkans (Bulgaria, 7 matches), as well as West Balkan-shifted individuals from the Battle of Himera in Sicily (8 matches) and a Scythian from Moldova with a southern-European profile (6 matches) (Fig. 6B, C, Tables S10-S11).

Regarding IBD matching between present-day Albanians and Roman period individuals, we find frequent links with samples from Croatia, Montenegro, and Serbia with a West Balkan-Anatolian-admixed genetic profile (R3481, R2041, R9669, R2053, R3685, R2041; 29 matches; Fig. 6C; Table S20). An individual from Roman Sisak (R2041) in Croatia with a mixed West Anatolian-West Balkan autosomal profile (Table S4) displays a high proportion of matches with present-day Albanians (6 out of 12, or 50%) (Table S20). Sample R2041 is also characterised by Y-chromosome haplogroup J-Z39727>J-FT151551; lineage J-Z39727* is also known in present-day Albanians^64,73^.

Beyond West Balkan connections, there is also a notable increase in matching individuals from Roman Serbia with partial (I28447, I27297, R6764; 13 matches) or full East Balkan-related ancestry (I15510, I15504; 14 matches) (Fig. 6B, C; Table S20). In absolute numbers, sample I15510 from Roman Viminacium has the highest number of matches with Albania_Medieval, AlbPostMdv and present-day Albanian samples, who make up 11 out of his 48 total matches (23%) (Table S20). Importantly, sample R6764 from Roman Naissus, also has a high proportion of Albanian matches (4/13 for 30%), and his Y-chromosome lineage (E-BY15412) is present today in low percentages in both Ghegs and Tosks^73^.

Furthermore, *f4*-statistics indicate some degree of allele sharing between these southeast Balkan-shifted individuals and Albania_Medieval and present-day Albanians (Fig. S6; Table S6). Together with the IBD-matching patterns observed here, this suggests that Albanians may also derive ancestry from a Central Balkan population with either an eastern-shifted, or an intermediate autosomal profile between West and East Balkan populations. However, we cannot exclude that the IBD signal with several samples from Roman Serbia may be due to a non-palaeo-Balkan component of their ancestry, such as Anatolian-related admixture (Table S4).

Overall, these results pinpoint the West and Central Balkans as the broad region from which the Albanians’ main ancestral sources derived in Antiquity, in accordance with the *qpAdm* analyses (Table S4) and their position on the PCA (Fig. 5A, B).

### Demography of the first Albanians, and the emergence of Ghegs and Tosks

Estimating the effective population size (Ne) of the proto-Albanians would allow us to understand their historical evolution and explain the strikingly high IBD-sharing between present-day Albanians and the Medieval individuals from Albania. To this end, we used Colate^83^, which infers coalescence rates for unphased, low-coverage genomes.

By plotting the inverse coalescence rates between different individuals within each present-day Albanian subpopulation, as well as the Medieval and Post-Medieval samples from Albania, we estimated that ethnic Albanians descend from an ancestral population of approximately 8,000-11,000 individuals (Fig. 7A). Plotting the inverse coalescence rates between and within Ghegs and Tosks resulted in an identical Ne (Fig. 7B), suggesting that both dialectal groups have an identical, shared population history. Comparing the inverse coalescence rates between present-day Albanians and Medieval-Post-Medieval samples results in slightly higher Ne (8k-18k) (Fig. S11). These different estimates may have arisen from a mismatch between fully and partially genotyped present-day and ancient individuals, respectively. Overall, our findings are supported by previous studies that have suggested that Albanians originated from a small, cohesive population around 1,500 years ago^41,82^.

**Fig. 7.**
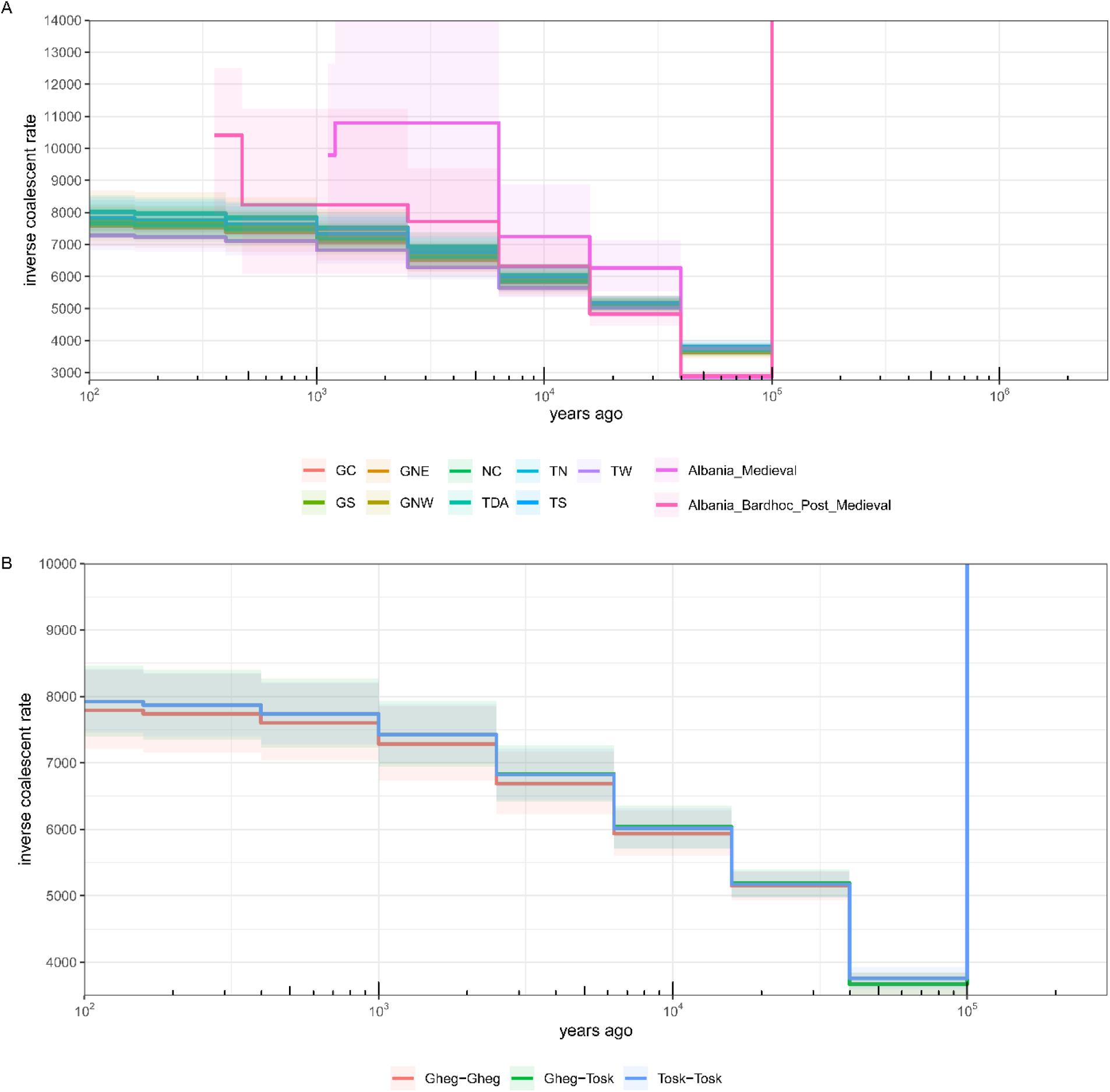
Effective population size estimation of present-day Albanians and ancient samples from the territory of present-day Albania using Colate. A) Inverse coalescence rates between different individuals within present-day Albanian subpopulations and ancient samples from Albania. B) Estimation of inverse coalescence rates between and within present-day Ghegs and Tosks.

To determine whether the founding population of Albanians experienced inbreeding due to its small size, we studied runs of homozygosity (ROH) – continuous stretches of DNA where alleles inherited from both parents are identical, indicating descent from a recent common ancestor^84^. Using the hapROH tool^85^, we tested all ancient and present-day individuals from Albania for ROH (Figs. 8A, B, S12; Table S21). We found that none of the present-day ethnic Albanian subpopulations displayed an excess of long ROHs consistent with inbreeding (Fig. 8B), possibly due to their exogamous social organization^86^. Exceptions included two individuals (GS3, TW6) from regions where exogamous traditions are less strictly enforced^86^, whose ROH is consistent with first cousin unions (Fig. S12). Crucially, none of the ancient post-Roman samples, including Albania_Medieval showed evidence of inbreeding (Fig. 8A), aligning with a small founding population that managed to maintain a mating pool that was sufficiently large to prevent excess homozygosity or endogamy. However, the Post-Medieval Roma-related individuals displayed significant inbreeding (Fig. 8A), consistent with previous work on present-day Roma populations suggesting a founder effect and subsequent bottlenecks in their demographic history^87^. Our study indicates that the endogamous nature of the Roma was firmly established by the 15^th^-18^th^ centuries CE, the period these samples date from.

**Fig. 8.**
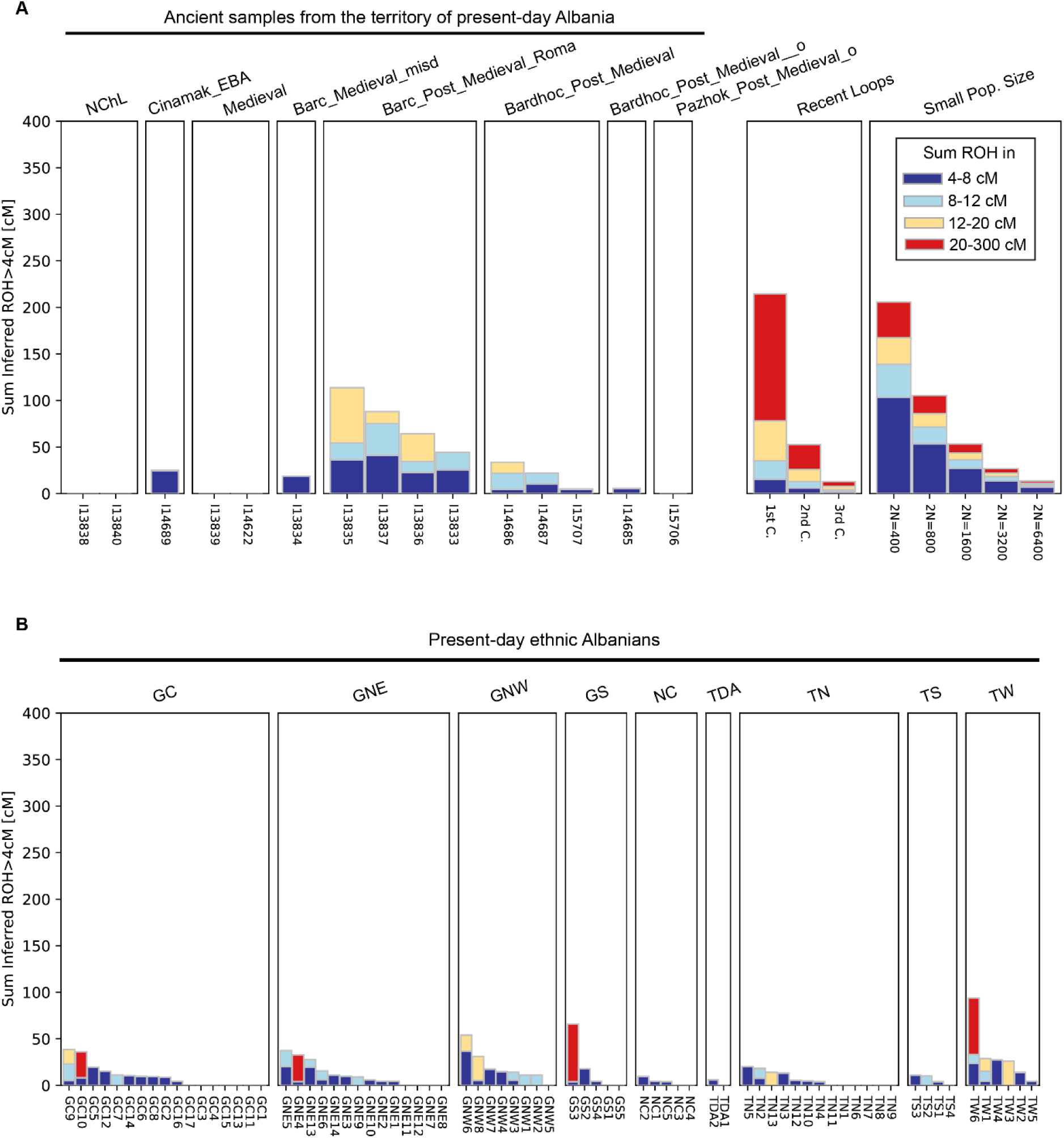
Runs of homozygosity (ROH) in populations from Albania over a 6,000-year transect, using hapROH. A) ROH in ancient individuals. The left-hand stacked plot illustrates RoH across various length categories. In contrast, the plots on the right present the anticipated total lengths of ROHs for cases that involve close relatives (“Recent Loops”) or small population sizes. B) ROH patterns across present-day ethnic Albanian subpopulations.

The small Ne of the founding Albanian population explains the exceptionally high IBD-sharing among present-day Albanians, with intra-subpopulation values ranging from 11-29 cM and both intra- and interdialect sharing at 9-13 cM (Table S16). This contrasts starkly with the 2-3 cM shared among average Europeans^41^. Combined with the high IBD-sharing between modern Albanians and the country’s Medieval inhabitants from 1,200 years ago (Fig. 6A; Tables S16), the results from Colate and hapROH are consistent with a very small founding Albanian population – before the emergence of Albania_Medieval. As the two Albania_Medieval samples were not closely related, we can hypothesise that the founding proto-Albanian population may have lived at least a few centuries before them, roughly during Late Antiquity or the very early Middle Ages. We should also note that wars, epidemics and mass migrations to Italy and Greece in the Late Middle Ages^15,30,65,88,89^ may have further reduced the genetic diversity of present-day Albanians compared to their Medieval ancestors. The results presented here, may therefore reflect only a subset of the founding proto-Albanian population.

The dialectal and geographical split between Gheg and Tosk Albanians is considered one of the earliest events in Albanian history, potentially occurring before or during the Migration Period^15,29,30,65^. To examine whether this dialectal division influenced genetic structure, we visualized all dialectal groups on a PCA (Fig. 9A). Despite long-standing dialectal divergence, Ghegs and Tosks and all of their subpopulations overlapped on the PCA and occupied the same PC with Medieval and Post-Medieval samples from Albania. Only more admixed individuals from different regions plotted towards the East European cline (Fig. 9A). Impressively, modern individuals from each subdialectal region, deriving from locations as distinct and distant as the Albanian Alps, the Presheva valley, the Himara coastline, the Korça plateau, and the Çamëria countryside matched both samples from Albania_Medieval, despite the latter originating from two regions separated by more than 250 km. The lack of substructure between Ghegs and Tosks, combined with their overlap with Albania_Medieval on the PCA, and the results of our *qpAdm* and ADMIXTURE analyses (Figs. 5, S2; Table S4), suggest that both dialectal groups derive from the same Medieval population.

**Fig. 9.**
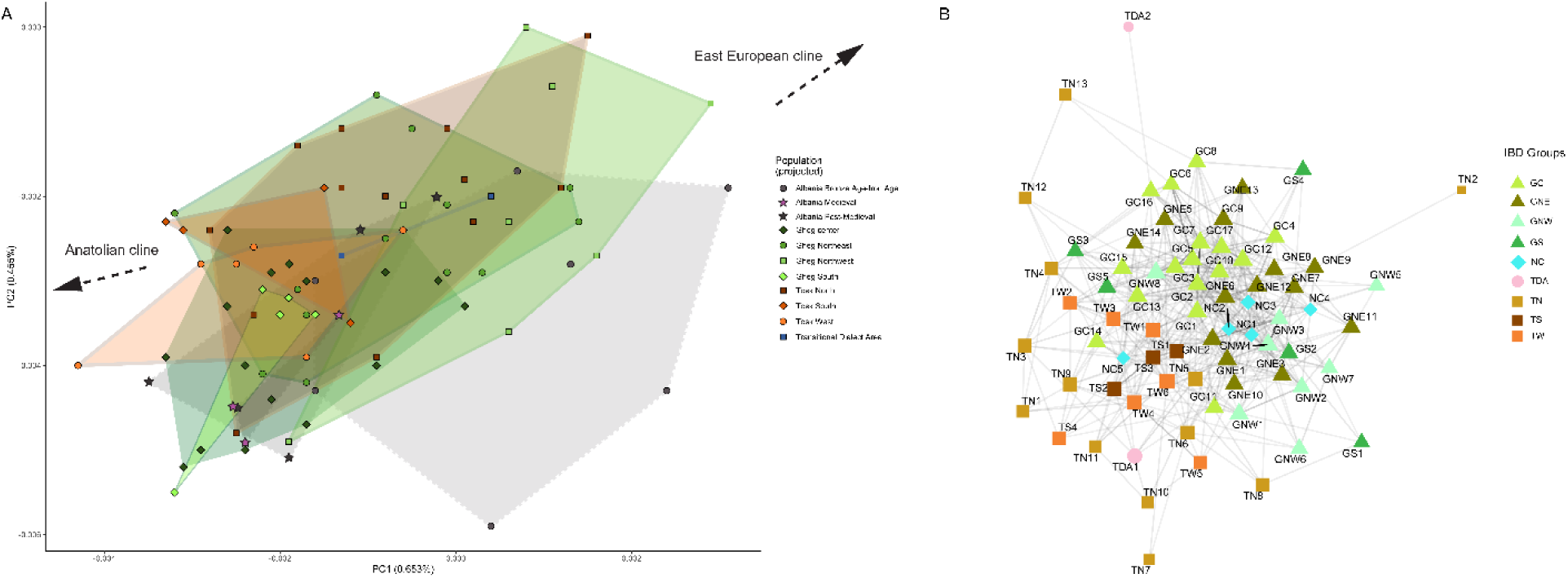
PCA and IBD-sharing patterns of present-day Albanians by dialect. (A) PCA of present-day West Eurasian samples (grey circles) with projection of present-day ethnic Albanians and their Bronze Age to Post-Medieval counterparts (faded grey polygons). Note the overlap between Medieval and Post-Medieval populations from Albania, and the shift of PCA coordinates of certain Gheg, Tosk, and Transitional Dialect individuals towards the East European cline (blue dashed line). (B) Intra- and interpopulation IBD-sharing patterns of present-day Albanians (>8 cM), showing a distinct separation between Gheg and Tosk populations.

Despite these similarities, inter-dialectal IBD-clustering showed that the two dialectal groups are distinctly separated from each other (Fig. 9B), which is consistent with mating networks that have been separated for a long period of time. This is further supported by the number of matching pairs within subdialectal and dialectal areas. At low IBD matching thresholds, pairs from different dialectal areas matched similarly to those within the same dialect, but at higher thresholds, matches were more common within the same dialect or subdialectal area (Fig. S13). This suggests distant shared ancestry for all Albanians, but that more recent ancestral ties are locally clustered. Interestingly, the two samples from the Transitional Dialect zone, displayed different IBD matching patterns; TDA1 only matched a Gheg Albanian from nearby North Macedonia, whereas TDA2 matched both GC and various Tosk subgroups, in line with the transitional nature of their dialect and geographical location^15^.

A peculiar finding concerns the differential IBD-sharing patterns between Albania_Medieval and the dialectal groups of present-day Albanians. In Ghegs, IBD-sharing with Albania_Medieval ranges from 10-21 cM, while in Tosks and Transitional Dialects, it ranges from 9.5-11 cM and 8.5 cM, respectively (Table S17). This occurs despite the Medieval sample with the largest number of matches to present-day Albanians originating from the southern locality of Shtikë. This may suggest that the Medieval individual from Shtikë was a migrant from more northern localities and accordingly shares larger IBD segments with present-day Ghegs. Conversely, Tosks and Transitional Dialects may have experienced different historical processes, such as kinship structures (e.g., endogamy), drift, or even admixture with currently unsampled palaeo-Balkan-derived populations, resulting in smaller IBD segments shared with Albania_Medieval.

### Large-scale persistence of pre-Migration Period haplogroups in present-day Albanians

Given that utility of uniparental markers in studying migrations and the transmission of languages^50,90^, we interrogated ancient and present-day Balkan Y-chromosome and mtDNA haplogroups to further refine our understanding of demographic changes in the region. Remarkably, we find that 80% of the paternal ancestry of present-day Albanians stems from pre-Migration Period populations of the Balkans and the Eastern Mediterranean (Fig. S10). We focus on the primary palaeo-Balkan lineages of present-day Albanians – haplogroups E-V13 (27-35%), J2b-L283 (17%), R1b-PF7562 (ca. 3%), and R1b-BY611>Z2705 (ca. 15%) (Fig. S10, Table S18) and report their regional history and phylogeny in the context of the formation of present-day Albanian populations.

Haplogroup J2b-Z638 branched off its parent lineage J2b-L283>J-Z622 around 3500-3000 BCE^64,91^. Current sampling suggests that the earliest J2b-L283 arose in pre-or-early Yamnaya-admixed groups from the north Caucasus^92^, and it appears abruptly on the Balkan aDNA record in the Bronze Age of Moldova (2900-2500 CE)^93^ and Serbia (2100-1800 BCE)^5,94^. In Serbia, J2b-L283 was found in a Maros cultural context, alongside an ancestral, upstream subclade of R1b-BY611 (Figs. 10A, S14). This places two of the most frequent paternal lineages of the Albanians (Fig. S10) in the Central-West Balkans by the EBA (Fig. 10A, B, S14). Haplogroup J2b-L283 experienced a major founder effect and diversification in the ancient populations of the Adriatic coast (Albania, Croatia, Montenegro), where it accounts for 50-70% of all paternal lineages during the BA-IA (Fig. 10A), and has been found in samples associated with major West Balkan archaeological cultural expressions, most notably in Maros, Glasinac-Mat, Cetina, Japodian and Liburnian contexts (Table S22)^5,94^. Coupled with its remarkably local distribution in pre-Roman times (Fig. S15), J2b-L283 may represent a reliable indicator of distant Bronze Age-Iron Age West Balkan paternal ancestry. The distributional expansion of J2b-L283 in northern and western Europe in Roman and post-Roman times (Figs. 9A, B, S15) is not surprising, as the West Balkans supplied the Empire with mercenaries, soldiers, and Emperors for centuries^7–9^. Within an Albanian context, J2b-Z638 subclades found in BA-IA, Roman and Medieval Albania (Bardhoc, Çinamak), Montenegro (Doclea, Velika Gruda), and Southern Croatia (Gardun, Gudnja cave), have daughter or sister lineages in present-day Albanians (Table S24), and today reach peak subclade diversity in Mat, Dibër, and Mirditë^73^, suggesting significant paternal continuity from ancient south-west Balkan populations identified as “Illyrians” by classical authors (Fig. S16). aDNA samples from Roman Serbia (Viminacium) and Late Avar-Medieval Hungary (Alattyán, Sárrétudvari) belong to J2b-L283 lineages related to those of present-day Albanians (Table S23), indicating an ultimately south-west Balkan paternal origin for these individuals, which corroborates inscriptional and historical evidence for transplantations of “Illyrian” soldiers along the Limes^8,9^.

**Fig. 10.**
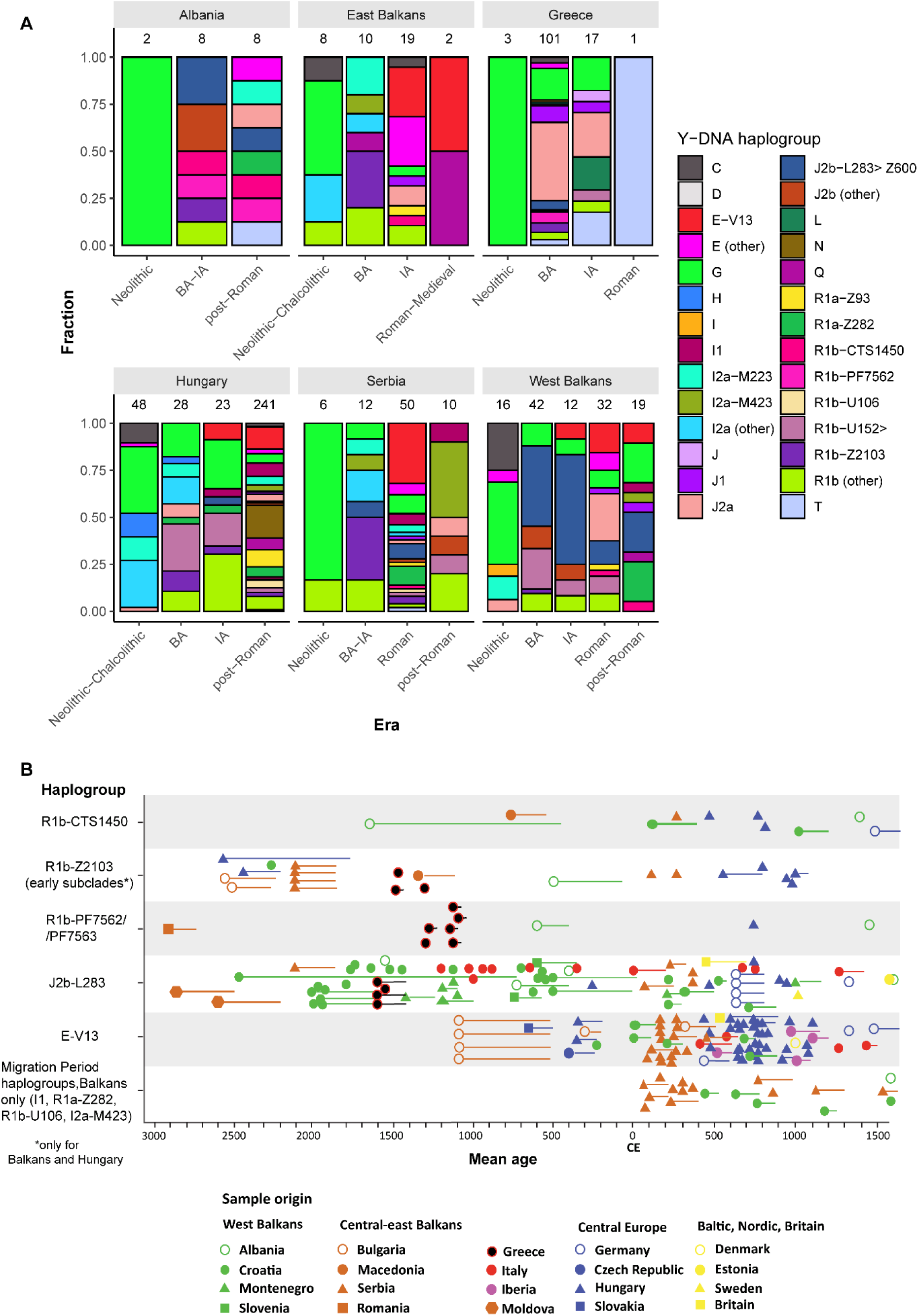
Haplogroup transformations of the Balkans and Hungary from the Neolithic to the Early Modern Period. (A) Y-DNA transect of the Balkans and Hungary over the past 8.000 years. Sample sizes shown above bars. (B) Temporal distribution of the principal Central-West Balkan-derived Y-chromosome haplogroups in European populations. Horizontal lines extending from some datapoints represent the archaeologically or radio-carbon-determined time range in which a particular individual lived. Compiled from Table S22.

The earliest recorded steppe-related paternal lineage to reach the Balkans is R1b-PF7562, which was first found in an EBA Yamnaya context in Romania (2893-2674 BCE)^93^ (Fig. 10). This haplogroup was subsequently found in BA-IA Albania and Greece (Fig. 10), most likely having arrived with the populations who first introduced Indo-European languages in the region. In the present-day Balkans, R1b-PF7562 is found almost exclusively among ethnic Albanians (at ca. 3% frequency; Table S18), where it is particularly rich in subclades (R1b-Z29758, R1b-Y83965)^64^. Based on the aDNA record (Fig. 10B) and the phylogenesis of Albanian-specific subclades (ca. 4,300 ybp for R1b-Z29758)^64^, R1b-PF7562 represents the survival of a palaeo-Balkan lineage that has persisted in the present-day territory of Albania since the BA-IA.

The most frequent paternal lineage among the Yamnaya, R1b-Z2103^6,52^, is represented in the ancient Balkans by daughter haplogroup R1b-CTS7556, which was first found in a Maros cultural context in Serbia (Figs. 9-10, S14)^94^. In turn, the primary descendant clade of R1b-CTS7556 is R1b-CTS1450, whose various daughter lineages appear in Bronze and Iron Age populations of northeastern Albania (Çinamak) and North Macedonia (Figs. 9-10, S14). Importantly, R1b-CTS1450, an ancestral subclade to R1b-BY611>Z2705, was found in BA-IA Çinamak (Fig. S14C). Out of all palaeo-Balkan haplogroups, R1b-BY611>Z2705 is the one with the most uniform distribution across all areas where Albanian has been spoken historically^73^, and it was also found in the Albania_Bardhoc Post_Medieval sample with the most IBD-matches with our present-day Albanian dataset (Table S10)). Based on the above, R1b-BY611>Z2705 likely played a crucial role in the Albanian ethnogenesis. The presence of R1b-CTS1450 in Çinamak and in North Macedonia suggests that the R1b-BY611>Z2705 lineage was located in this region or nearby during BA/IA. Present-day R1b-Z2705 aligns with this distribution, with the oldest branches being found in Albania, Kosovo, and southern Serbia^64,73^.

Despite being one of the most frequent haplogroups in present-day Balkan populations^69,72,95–97^, the origins of E-V13 remain enigmatic. The earliest records of this haplogroup in the Balkans are in Early Iron Age Bulgaria within a “Thracian” cultural context, at a frequency of 26% (Figs. 9A, B, S16) and in a Scythian from Moldova that plots close to classical era Balkan populations (Fig. 3A). Later, it appeared in Iron Age populations from Croatia and Hungary at lower frequencies (8%). By the early Roman era, populations rich in E-V13 likely experienced significant and rapid demographic growth, with this haplogroup appearing more frequently in areas where it was previously rare (Croatia, 15%) or unsampled (Serbia, >30%) (Fig. 10A, B).

Our findings suggest that the increased frequency of E-V13 between the Iron Age the Roman period examined herein is frequently associated with samples having a fully or partially Bulgaria_EIA-related autosomal profile. Specifically, 39% of individuals carrying this haplogroup in Roman Serbia and Avar-era Hungary cluster with BA-IA populations from Bulgaria on the PCA (Fig. 5A) and are identified as cladal with a Bulgaria_EIA or Mycenaean_BA-related source population in *qpAdm* models and D-statistics (Fig. S17, Tables S4, S6). A further 33% derive 24-70% of their ancestry from such a proxy (Fig. S17, Table S4). These E-V13-carrying individuals, along with another Scythian from Moldova who clusters with Balkan Iron Age populations (Fig. 3A), also display IBD-sharing with Bulgaria_EIA (Table S17). This may support historical records of “Thracian” groups, known as the “Bessi,” and the “Moesians” present throughout the Balkans until the 6^th^ century CE^2,35,98,99^.

Furthermore, individuals from Roman Serbia cladal to Bulgaria_EIA show considerable diversity in E-V13 subclades (e.g., E-BY5490, E-FGC33621, E-BY5022, E-CTS9320; Table S22), suggesting either proximity to or inclusion in the epicentre of E-V13 diversity, or significant migrations from it. Notably, two individuals (15518, I15504) belong to subclade E-FGC33646 (Table S23), from which 3% of present-day Albanians descend^73^. The autosomal profile associated with these results suggests that some E-V13 subclades in modern Albanians may originate from Bronze and Iron Age Balkan populations with an autosomal profile fully or partially related to Bulgaria_EIA. Whether this autosomal profile was found beyond the territory of present-day Bulgaria in pre-Roman times, such as the Carpathian Mountains or the Danubian basin, remains an open question.

However, not all populations with E-V13 were characterized by a Bulgaria_EIA-related autosomal profile. For example, two E-V13-bearing mercenaries from Himera in Sicily plot with West Balkan populations on the PCA (Fig. 3A), may derive large parts of their ancestry from BA Kosovo or Serbia (S1 Text, Table S4); and share IBD segments with both “Illyrians” (8.5 cM) and “Thracians” (11.5 cM) (Table S17). It is likely that these mercenaries may have originated from a Central Balkan zone of linguistic contact between “Illyrian” and “Daco-Thracian” groups^1,7^, thereby displaying genetic affinities to both. Importantly, the E-V13 subclade found in one of the Himera mercenaries (E-CTS6377) is characterized by daughter branches from which another 7% of Albanians descend (Table S23). Additionally, two Roman individuals from Croatia (R3664, R3659) also harbored E-V13 (Table S22), despite the absence of Bulgaria_EIA-related ancestry. Furthermore, 50% of Roman period individuals with E-V13 are modelled as deriving 15-80% of their ancestry from West or Central Balkan Bronze and Iron Age populations (Fig. S17).

The presence of E-V13 in the Himera individuals may indicate either: (1) early arrivals from more eastern areas whose Bulgaria_EIA-related ancestry became diluted, or (2) the IA/Roman period spread of at least some E-V13 lineages from central/western regions, possibly with an autosomal profile intermediate between west and east Balkan groups. One candidate population are the “Dardanians”, situated in the central part of the Balkans, having been influenced by other “Illyrians” in the West and by “Daco-Thracians” in the East^1^. Supporting the second possibility, the BA/IA ties of modern Albanian E-V13 lineages generally point to regions to the north, rather than the east of the present-day Albanian speaking area (Table S23). Based on these findings, it is possible that populations from which many Albanian-specific E-V13 subclades descend, regardless of linguistic affiliation, were largely or fully West Balkan autosomally during the Roman period.

To gain insights into the ethnogenesis of present-day Albanians, we plot the mean Y-full TMRCAs of Albanian-specific subclades of palaeo-Balkan (E-V13, J2b-Z638, R1b-BY611, R1b-PF7562, I-M223) (Fig. 11), Roman (J1) and Migration Period-related (I1, R1a-Z282) haplogroups found in present-day Albanians. Remarkably, all palaeo-Balkan haplogroups show a sudden and steep increase in subclade diversity between 200-800 CE (Fig. 11A), aligning with linguistic and historical hypotheses on the origins of Albanians^28,33,34,98,100^, as well as our findings on the effective population size of the founding Albanian population (Fig. 7).

**Fig. 11.**
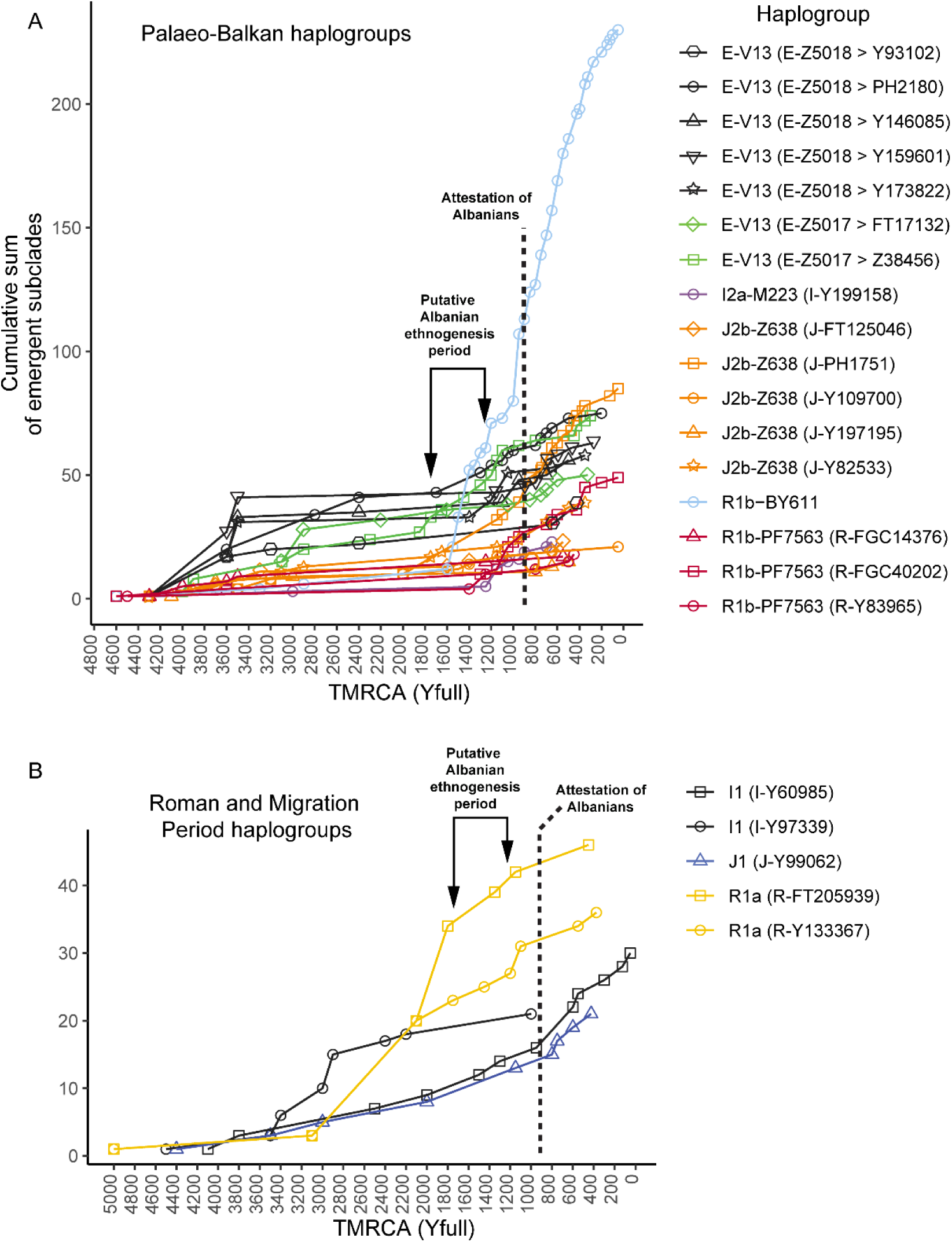
Graphical representation of clade formation (cumulative sum of new subclades) of the principal palaeo-Balkan and Migration Period-related haplogroups of present-day Albanians, using TMRCA estimates from Y-full. A) Plots for palaeo-Balkan-related haplogroups. B) Same, for Migration Period haplogroups. Plotted using data from Table S24.

Most paleo-Balkan haplogroups show signs of severe bottlenecking from the early Iron Age to Late Antiquity, while some E-V13 subclades exhibit a more continuous subclade diversification throughout the Iron Age (Fig. 11A). This trend suggests that some subclades of E-V13 may have followed a different demographic trajectory compared to those of J2b-Z638, R1b-BY611, R1b-PF7562, and I-M223, implying that they may have originated in different populations. Overall, the rate of diversification of all E-V13 Albanian-specific subclades increased significantly from 200 CE onwards, mirroring the pattern of other Albanian lineages (Fig. 11A). This may indicate that some E-V13 lineages merged and co-expanded with local populations during the Iron Age or early Roman Period, which could explain the absence of E-V13 in the aDNA transect of Albania, despite being the most common haplogroup in the present-day Albanian population (Fig. S10).

Our re-analysis of the Y-chromosome of a low coverage individual (I17623) from 9-10^th^ century Dukat in Albania, assigned his paternal lineage to haplogroup E-L539 (Table S22), possibly an indicator of E-V13 among early Albanians.

Regarding Roman (J1) and Migration Period (I1, R1a-Z282) haplogroups in present-day Albanians, these follow a drastically different trend compared to palaeo-Balkan haplogroups throughout the Iron Age and Roman period, but they start converging in the early Middle Ages (Fig. 11B). This suggests that they had merged with the proto-Albanian population by the later stages of the putative ethnogenesis period (200-800 CE), or shortly thereafter. Together with *qpAdm* and IBD data, the principal Y-chromosome haplogroups add further evidence for large-scale continuity of present-day Albanians from local groups. Continuity is also mirrored in rarer lineages, such as haplogroup T-Y206597, found in the Medieval individual from Kënetë (Table S23), which today comprises almost exclusively Albanians^64^.

Regarding the female ancestors of present-day Albanians, mtDNA also supports large-scale continuity from Bronze and Iron Age Balkan ancestors. Haplogroups at the level or directly upstream of H1e1b1c, H7c1a, H7c4a1, H11a2b, H12a*, H13b1-a, H-d9, H55-a, U3b1*, U3b1b1b, X2m1a1*, T2a1b1a1b2*, T2a1b1d, which collectively account for 22% of all Albanian matrilines, have been found in the Bronze and Iron Age populations of Bulgaria, Greece, and Montenegro (Table S19). Crucially, haplogroups found in Post-Medieval samples from Albania – namely T1a1l, H7c1a, J1e, U5a1a2a-a6 are also found almost exclusively in present-day Albanians (Table S19), further supporting their direct descent from this population.

## Discussion

Our palaeogenomic transect reveals all the major phases in the history and prehistory of the territory of present-day Albania. We detect the arrival of the first putative Indo-European speakers into Albania at approximately 2700 BCE, who likely came directly from the Pontic-Caspian steppe (Table S7). However, it remains uncertain whether the language spoken by this population influenced the linguistic landscape of later historical periods. By the Late Iron Age, the cultural and genetic formation of the Illyrians – Albania’s first historically documented inhabitants – appears to have been complete^7^. The Bronze and Iron Age population of Albania was part of the broader West Balkan continuum (Fig. 3). Indeed, our IBD analysis reveals connections of Albania’s Illyrians both with other peoples of the Western Balkans who may have spoken the same or related languages^7^, as well as with populations further to the east, perhaps with groups known as Paeonians (Fig. 3C). The Roman period represents a chronological gap in our transect.

However, our group-based, PCA, IBD, and uniparental analyses indicate that Albanians descend from a palaeo-Balkan population that experienced significant demographic increase approximately between 200-800 CE (Fig. 11A), perhaps after a population bottleneck that was caused by yet undetermined factors. We show that by the Early Middle Ages, autosomal profiles typical of many present-day Albanians were already formed (Fig. 4A, B). By the 8^th^-10^th^ centuries CE, the Early Medieval ancestors of Albanians were present in a broad geographical area, consistent with the available genetic and archaeological evidence.

Considering the overwhelming support of our study’s genetic analyses for a descent of all present-day Albanians from the country’s Medieval inhabitants, we propose that the samples from both Kënetë and Shtikë were part of the same founding population (who may have formed a few centuries prior) and in all likelihood speakers of proto-Albanian. If our interpretation is sound, it pushes the definitive presence of Albanians in Albania centuries earlier than their first historical attestation, highlighting a potential gap in Eastern Roman historical records regarding their acknowledgment or mention of this population^19^. The increased subsequent involvement of Albanians in Eastern Roman affairs (especially in a military context), may be the underlying reason why they attracted the attention of Greek-speaking historians only by the 11^th^ century^19^.

Based on the two samples from Kënetë and Shtikë – which may not capture the entire genetic diversity of Medieval populations from Albania, the Early Medieval ancestors of the Albanians, both in North and South Albania experienced no contribution from surrounding Slavic populations (Fig. 4A, B; Table S4) and maintained the highest levels of Bronze and Iron Age West Balkan ancestry observed in the present-day Balkans (68-84%). The only detectable post-Iron Age ancestry shift in the ancestors of Albania’s Medieval inhabitants is represented by a 16-32% contribution from an Anatolian or southeast Balkan-related population, which likely took place during the preceding Roman period, as happened elsewhere in the Balkans^3^. Remarkably, 1,200 years later, the genetic profile of Albania_Medieval represents on average 80-90% of the ancestry of present-day Albanians, depending on the East European-related proxy used in *qpAdm* analyses (Fig. 5).

Despite being largely unaffected by the demographic changes that took place during the Migration period, the historical Albanians did not emerge in isolation. At the peak of the Migration Period, the Medieval population of Albania displayed genetic links as far as Hungary and Romania (Fig. 6A-C; Tables S10-S17). Two post-Medieval (1400-1700 CE) samples from Albania exhibit significant admixture with East European-related populations (Table S4). Additionally, a substantial proportion of the Albanian-speaking diaspora in Southern Italy, known as the Arberesh, who settled from the 16^th^ century onward and originated primarily from the Peloponnesian Albanians^88^ carries East European-derived Y-DNA lineages (R1a-Z282, I-M423)^71^. This suggests that the diffusion of East European ancestry among Albanians had started already by the Late Medieval period. On the other hand, present-day Albanians display variable levels of this type of ancestry (Fig. 5, Table S4), indicating that this diffusion post-dated the initial stages of the Albanian ethnogenesis, in agreement with the absence of Eastern European ancestry in the Early Medieval ancestral population and the divergent expansion of East European derived lineages until the ethnogenesis period (Fig. 11B). This indicates complex historical interactions with East European-related populations, as suggested by toponymy and linguistics^21,66–68^.

Whether East European-related ancestry entered the ancestors of present-day Albanians in an unadmixed (CEE_Medieval) or Balkan-admixed (Montenegro_Medieval) form remains unclear, although a combination of the two is also likely. The Slavic migrations into the Balkans began in the 6^th^ century CE and likely involved largely unadmixed individuals, though we lack aDNA from this period. By the 7^th^-10^th^ centuries CE, individuals with Eastern European-related ancestry were admixed with Roman Balkan populations, as shown by a previous study^3^. Our models on present-day Albanians show lower p-values for CEE_Medieval (unadmixed Eastern European profile) compared to Montenegro_Medieval (Eastern European-Balkan-admixed profile), which is not an indicator of model rejection^42^. Given that the autosomal profile of the incoming Slavs remains uncertain until more extensive aDNA studies in present-day Albania are conducted, we consider CEE_Medieval and Montenegro_Medieval as minimal and maximal estimates of East European-related admixture in present-day Albanians.

The geographic and linguistic origins of the Albanians have been debated for centuries. Our analysis of uniparental markers reveal that a significant proportion of the paternal ancestry of present-day Albanians derives from groups ultimately descending from the BA-IA West Balkans (Table S23), including those traditionally known as “Illyrians” (Fig. 11A; S16), as well as Central Balkan groups, which reflects our findings on autosomal ancestry. Our IBD analysis is consistent with this scenario, as it reveals genealogical connections between Albanians and Roman era southern Croatia, Montenegro, and Serbia (Fig. 6). These data are in agreement with linguistic hypotheses^30,33^ that propose an original proto-Albanian homeland spanning mountainous regions in present-day northern Albania (Mat, Martanesh, Diber, Mirdite), southwest Kosovo, and part of North Macedonia. This area mirrors the contact zone between “Illyrians” and a potentially related Iron Age-Early Roman group known as the “Dardanians”^1^, both of which may have contributed to the ancestry of present-day Albanians. The presence of several E-V13 clades in present-day Albanians, which in the Roman period have been found in Serbia in individuals with a Central and southeast Balkan profile (Tables S4, S23), provides evidence for this hypothesis.

However, even though our analyses resolve the geographic and temporal origins of the Albanians, inferring the affinities of their language remains challenging, especially considering that there is no indication of the survival of “Illyrian” in the Balkans following the first centuries of Roman rule^7,8^. Furthermore, the Albanian language displays Latin loans from both the Western and Eastern Balkans^101^, which attests to linguistic influences in the northern parts of Albania and the region of Montenegro, where Western Balkan Romance is attested^101,102^. Considering the genetic affinities and the small population size of the proto-Albanian population (Fig. 7), it is likely that the founding population of the Albanians lived in a restricted geographical area that shielded them from the tumultuous events of the Migration Period. Future aDNA samples from the Roman era Central Balkans may further help pinpoint the precise geographical location of the ancestors of present-day Albanians.

We should also stress that although our palaeogenomic analysis provides novel insights into the demographic history and ancestral origins of present-day Albanians, we recognize that one cannot draw direct inferences about cultural or linguistic identities solely from genetic data. Our findings reveal genetic continuity between Early Medieval and present-day Albanian populations, yet this does not necessarily imply linguistic continuity. As ongoing debates surrounding the spread of Indo-European languages have shown, language transmission is a multifactorial process that may occur independently of large-scale genetic shifts. A notable example is the case of Basque– a non-Indo-European language isolate spoken by a population with high levels of steppe-related ancestry^50^. Therefore, any interpretation of the linguistic history of the Albanians based solely on genomic evidence must be approached with caution, acknowledging the influence of sociocultural, historical, and even environmental factors that fall beyond the scope of genetic data.

While the quest for the origins of the Albanian language will certainly continue, we expect that the findings from the present study will be pivotal in shaping these debates and providing the necessary framework for more extensive research on the genetic ancestry of the ancient and present-day inhabitants of Albania.

## Materials and Methods

### Datasets

#### Ancient Individuals

For admixture-based analyses, we used more than 660 previously published ancient genomes from the Allen Ancient DNA Resource (AADR) (version v54.1_1240K_public)^103^, and the studies of Antonio et al. (2024)^39^ and Reitsema et al. (2022)^104^, which include 22 ancient individuals from the territory of present-day Albania. We selected data from these datasets based on their genetic, geographical, and historical relevance to our study. We did not use genomes of low coverage (<15k SNPs), close relatives, or with high levels of contamination. We should note that on several occasions, our sample naming differs from that given in the original study of the corresponding sample, based on insights from the PCA, admixture modelling, and dating interpretation. A list of all the samples and our naming conventions used in the *qpAdm* models and PCA of this study can be found in Tables S1-S2.

#### Present-day individuals

Permission to collect whole genome sequences of present-day individuals was granted by the Council of Ethics of the Academy of Sciences of Albania (approval date 15.07.2024), in compliance with the guidelines specified by the Declaration of Helsinki. A total of 74 participants with confirmed ethnic Albanian ancestry at least four generations back, and who had previously tested with 30x direct-to-consumer whole genome sequencing (70 testers with Dante Labs, L’Aquila, Italy; participants GC1, GC4, GNE11 and GNE 14 tested with YSEQ GmbH, Berlin, Germany) were contacted and informed about the goals and methods of this study. Each participant gave informed consent for their sequencing data to be used for scientific inquiry. All experimental procedures were undertaken following the approved guidelines by the Council of Ethics of the Academy of Sciences of Albania. We selected 72 participants who have all four grandparents from their dialectal area of origin, except for two individuals (NC1, NC5) who had one grandparent from a neighbouring dialectal area. Due to our sampling strategy, the geographic range of the present-day individuals captures the genetic diversity of Albanian speakers with origins from Albania, Kosovo, Montenegro, northern Greece, North Macedonia, and Serbia.

DNA extraction, quality control, library preparation, and sequencing were performed by the sequencing providers (Dante-Labs, L’Aquila, Italy; YSEQ GmbH, Berlin, Germany), who undertook alignment with the human reference genome GRCh37.

### Statistical Analysis

#### PCA

We undertook Principal Component Analysis (PCA) on a subset of the HO^54^ and newly genotyped Reitsema et al. (2022)^104^ dataset using the ‘smartpca’ function (v16000) in EIGENSOFT (version 7.2.1)^105^ (Fig. S1). SNP datasets in EIGENSTRAT format were combined using the mergeit function included in EIGENSOFT, with default parameters, aside from allowdups: YES; outputformat: EIGENSTRAT; strandcheck: NO; hashcheck: NO. We converted .bam files to EIGENSTRAT format using the mpileup function from the SamTools software package (v1.16), with optional flags -R -B -q30 -Q30. The resulting pileup files were processed into EIGENSTRAT using the pileupCaller function from the SequenceTools software package (v1.5.2 - https://github.com/stschiff/sequenceTools), with default parameters and randomHaploid as calling method.

For compatibility with the HO dataset, SNPs within 1240K SNP datasets corresponding to those in the HO dataset were renamed prior to the datasets being combined using mergeit. We then projected ancient samples onto present-day individuals with “lsqproject:YES” and “shrinkmode:YES”, in three chronologically distinct datasets (Neolithic-Early Bronze Age, Bronze Age-Iron Age, Roman-Post-Roman), resulting in Figs 2, 4, 5 and 8. The final dataset of the present-day and ancient samples used for the PCA is provided in Table S2.

### ADMIXTURE

We used ADMIXTURE (v1.3.0) to analyse 1,118 present-day and ancient human samples. Most of the individuals were sourced from the Allen Ancient DNA Resource or AADR (v54.1_1240K_public) from the David Reich Lab. Sixteen of the samples came from Antonio et al. 2023^106^. The initial set of 1,150,639 autosomal SNPs was pruned for linkage disequilibrium in PLINK 1.90 with parameters --indep-pairwise 50 25 0.2, which resulted in a final set of 322,534 SNPs. All individuals with a genotyping rate of less than 5% were removed from the analysis. ADMIXTURE was run with the number of ancestral populations (K) ranging from 2 to 8. The results, particularly at K7 and K8, are similar to our formal *qpAdm* statistics-based analyses.

#### Outgroup *f3* and *f4*-statistics to assess shared genetic drift

We employed the ADMIXTOOLS v.2.0.0^42,43^ package in R, operated using RStudio (version 4.2.3), in order to estimate outgroup *f3*-statistics of the form *f3(outgroup; population A, population B)*, where outgroup = Cameroon_SMA, population A = the tested individual or metapopulation, and population B = selected ancient populations from our working dataset, respectively. Outgroup *f3*-statistics estimate pairwise genetic affinity via allele sharing^43^. The higher the value of the *f3*-statistic the higher is the genetic affinity between population A and the tested ancient population(s) B.

To gain insights into genetic distances between populations in comparative terms, we used f4-statistics, formulated as f4(outgroup, population A; population B, population C). The f4-statistics analysis enabled us to address critical questions about the genetic connections between populations, in particular, some of the following: 1) determining the most closely related European EBA populations for Albania_Cinamak_EBA; 2) further testing the relationship between carriers of Y-chromosome haplogroup E-V13 and Bulgaria_EIA ancestry; 3) identifying closely related populations for Albania_Medieval; 4) assessing allele-sharing between present-day Albanians and unadmixed (CEE_EarlyMedieval) and admixed (Montenegro_Medieval) East European-related proxies.

A significant advantage of f-statistics is their robustness against factors like long divergence times, small population sizes, missing data, and lower data quality. We should note that we did not use samples with less than 30k SNPs in any of our f-statistics analyses, which could have led to less reliable results.

### *qpAdm* admixture modelling

We employed *qpAdm* software from ADMIXTOOLS v.2.0.0^43^ to model the ancestry of the examined populations by using a base set of references approach^42^, with an emphasis on the samples from Albania. A model was accepted as statistically plausible if its p-value was ≥0.01, as followed by previous work^3^. Z-scores were automatically estimated by our *qpAdm* script, and only values to close to or > Z = 3 are reported. An extensive description of our *qpAdm* models and the rationale behind the chosen source and reference populations can be found in the Supplementary Material.

## Mobest analysis

The Mobest R package facilitates the spatial and temporal analysis of human genetic ancestry, employing Gaussian process regression (kriging) to generate a continuous map over both space and time [https://github.com/nevrome/mobest]^45^. It also calculates similarity probabilities within the interpolated field, which can be interpreted as origin probabilities in certain circumstances. For this reason, the use of Mobest could be useful in pinpointing the affinities of the Albania_Medieval individuals, which, unlike other contemporary samples, did not experience a shift towards the East European cline.

To this end, we used a reference dataset comprising 5,664 published individuals from the Allen Ancient DNA Resource^103^ and the samples from the Antonio et al. (2024) study^39^ that were used for the *qpAdm* analyses (Table S1). The reference dataset was filtered to only include samples covering a geographic area between 29.9° and 70° latitude and -24° and 70° longitude (in EPSG:4326 projection), to a specific chronology (150 to 5,000 years before present). The samples were subjected to a kernel size of 800 (corresponding to 800 km in space and 800 years in time) for interpolation. The predication grid was set to 50 by 50 km tiles. As genetic input we used the first two PCs of the West Eurasian PCA as well as the first two PCs of a PCA only including European individuals. Samples with less than 15k SNPs were excluded. The relative search time was set to 0, thus, the probability surface indicates the highest genetic-geographical match at the mean date of the respective individual. The observed probabilities indicate the presence or absence of genetic representation in different regions and time periods. Regions with zero probability are areas lacking sufficient reference samples, leading to lower confidence in genetic interpolation for those areas. Conversely, regions with higher probability scores represent better-supported genetic matches at the mean dates of the samples used.

### Admixture time estimation

To investigate the timing of the introduction of Yamnaya-related and East European-related ancestry into Albania_Cinamak_EBA and present-day Albanians, we utilized the Distribution of Ancestry Tracts of Evolutionary Signals (DATES) tool^44^. This approach estimates admixture timing in generations before the period during which the sample lived, converting these into years with an average generation length of 28 years (following the protocol proposed by the tool’s developers), and to absolute dates by considering the sampling age of ancient genomes.

A good model fit is indicated by a normalized root-mean-square deviation (NRMSD) of less than 0.7, a Z-score greater than 2, and an admixture time (λ) of less than 200 generations^44^. However, these criteria serve as general guidelines rather than strict cut-offs for the acceptance or rejection of admixture timing estimations.

### Uniparental marker analysis and interpretation

Ancient DNA Y-chromosome haplogroup data were aggregated from the literature (Tables S22-S23). In most cases, the terminal subclade we cite was assigned by yfull.com^64^ and the free, publicly available FamilyTreeDNA Discover Beta^TM^ (Gene by Gene Ltd), both being broadly used resources for inferring human uniparental haplogroups and their TMRCA^6,107,108^. Additionally, publicly available raw data from Albania by Lazaridis et al. (2022) ^6,13^ and Central-Western Balkan samples by Antonio et al. (2024)^39^ were manually evaluated in the present study using IGV ^109^ and were also called with SAMtools and BCFtools^110^ by Open Genomes (Ted Kandell), Open Genomes and the Society of Serbian Genealogists “Poreklo” (Milan Rajevac), and Leo R. Cooper. TMRCAs are publicly available at yfull.com and FamilyTreeDNA Discover Beta^TM^. The present-day Albanian data populating the yfull.com and FamilyTreeDNA Discover Beta^TM^ Y-chromosome and mtDNA public databases stem from direct-to-consumer whole genome sequencing tests (BigY^111^, Nebula Genomics, Dante Labs, YSEQ).

To generate Fig. 10, we assembled all Y-chromosome haplogroups from Table S18, which we used to produce a bar-chart in RStudio using ggplot2^112^. Regarding Fig. 11, which plots the TMRCAs of the principal Y-chromosome haplogroups of present-day Albanians (E-V13, J2b-L283, R1b-BY611, I-M223, and R1b-PF7562) (n = 1088), we consulted the corresponding phylogenetic trees at yfull.com^64^. We then assembled in Table S24 the TMRCAs of the Y-chromosome subclades associated with Albanians and their expansions into neighbouring regions (Greece, Bosnia, Montenegro, North Macedonia, Serbia). We did not include the TMRCAs of subclades lacking an association with Albanians (i.e. not having any Albanians in their daughter lineages). To study the Y-chromosome and mtDNA haplogroup distribution of the present-day ethnic Albanian population, we used both academic samples from the scientific literature^69–72,79^ and the publicly available datasets of rrenjet.com^73^ (n = 1816) and Yfull^64^ (Tables S18-19). We should note that the academic samples often comprise low-level STR tests, which do not provide terminal subclades for some haplogroups, especially those downstream of R1b-M269.

### Preprocessing of the present-day Albanian dataset

The data for the present-day Albanian samples were provided as analysis-ready BAM files (GC1, GC4, GNE11, GNE14) or VCF files genotyped by the Dragen/GATK4 Haplotype caller best practice analysis pipeline. Only the VCF file of sample GNE4 was genotyped against the hg19 human reference, the rest of VCFs were called against the hs37d5 (GRCh37) human reference.

For the VCF files the multiple nucleotide polymorphism (MNP) variants were split by the ***‘bcftools norm’*** [<u>10.1093/gigascience/giab008</u>] command with the *‘-m –any’* options; then filtered for high confidence SNPs by ***‘bcftools view’*** with the *‘-f PASS -v snps’* options; and finally normalised to the reference by *‘**bcftools norm’*** with the *‘-m +snps -f hs37d5.fa.gz’* options. In the case of individual GNE4, the hg19 reference FASTA file was used for normalisation and the output was converted to the GRCh37 chromosome notation style (no ‘chr’ tag before chromosomes). We restricted the variants genotyped to the chromosomes only where the hs37d5 and hg19 version of GRCh37 genome coordinates and reference are identical (except for different chromosome naming). For convenience, we created a joint VCF file from the single VCF files to contain all called GTs. Since all the VCF files were generated from high coverage shotgun WGS data, we assumed that at any position where no variants were called in a particular sample, the sample had homozygote reference alleles. Accordingly, the VCF files were merged by the **‘bcftools merge’** command with the *‘-0’* option (assume genotypes at missing sites are 0/0).

### Preparation of the 1240K marker set data of present-day Albanian individuals

The analysis-ready BAM files were genotyped by ***‘bcftools mpileup’*** with the *‘-f hs37d5.fa.gz -I -E -a FORMAT/DP -R 1240K_sites.tsv.gz’* and the ***‘bcftools call’*** commands with the *‘-Aim -C alleles -T 1240K_sites.tsv.gz’* options. The joint VCF file of present-day individuals containing GATK-genotyped data was filtered for the 1240K position by ***‘bcftools view’*** with the *‘-R 1240K_sites.tsv.gz’* option and merged to the data that were genotyped from BAM files, using the *‘bcftools merge’* command with the *‘-0’* option. The unphased present-day Albanian joint VCF dataset was imported to PLINK (version 1.9) [www.cog-genomics.org/plink/1.9/] binary format with the *‘--keep-allele-order --output-chr M --real-ref-alleles --recode vcf-iid’* options. The diploid data were pseudohaploidised by the ***‘pseudoHaploidize’*** command from the correctKin toolset [https://doi.org/10.1186/s13059-023-02882-4]. In all downstream analysis where modern samples were coanalysed with ancient data only the pseudohaploidised dataset was used.

### Preparation of the genotype data for IBD sharing analysis

The GRCh37 joint VCF file of present-day was phased by SHAPEIT5 (version 5.1.1) with the ‘***phase_common***’ tool [https://doi.org/10.1038/s41588-023-01415-w]. The phased variants were restricted to 1240K positions by ***bcftools view’*** with the *‘-R 1240K_sites.tsv.gz’* option.

The publicly available ancient samples with less than 0.4x mean genome coverage were excluded from the IBD analysis. Furthermore, only samples with the PASS quality flag (as noted in the Allen Ancient Data Repository (AADR) [https://doi.org/10.1038/s41588-020-00756-0] annotation file) were included in the analysis. Additionally, we included 8 high coverage Mesolithic-Neolithic age ancient samples (I6767.SG, R2.SG, R3.SG, R7.SG, R9.SG, R10.SG, R15.SG, Stuttgart.DG) in our dataset as a control for validating IBD filtration and for the detection of false positive IBD matches (Table S8). We assumed that no Medieval-Post-Medieval-present-day individuals would share IBD segments with the control samples.

The high coverage analysis-ready GRCh37 BAM files of present-day Albanians and the publicly available ancient BAM files were imputed by the GLIMPSE2 [https://doi.org/10.1038/s41588-020-00756-0] imputation framework. The 1KG Phase III joint VCF data were used as a reference and the imputation was restricted to the common, bi-allelic SNP variants of the autosomal chromosomes (∼78.1 M SNP marker positions). The phased/imputed GTs were restricted to 1240K marker set positions by ***‘bcftools view’*** with the *‘-R 1240K_sites.tsv.gz’* option. The 1240K restricted imputed data and the phased modern data were merged by the **‘bcftools merge’** command with the *‘-0’* option.

### IBD sharing analysis

The 1240K restricted data were lifted to hdf5 format by the ancIBD framework (version 0.6) [https://doi.org/10.1038/s41588-023-01582-w] and the raw IBDs were called using the *“min_cm=4 e_model=’haploid_gl2’, h_model=’FiveStateScaled’, t_model=’standard’”* parameters. Instead of using the default 220 SNP/cM density criterion for filtering raw IBDs, a novel approach for true IBD filtration [https://github.com/zmaroti/ancIBDfiltration] was applied. Our filtration method is based on the hypothesis that true IBD fragments are randomly distributed in the genome and higher-than-expected IBD counts at any genome location are due to the excess of false positive raw IBDs. Based on this hypothesis, the method downscores the markers at the low confidence genome regions leading to automatically masking of such regions. A detailed description of the method, including the in-depth analysis of the experimental raw IBD distribution, mathematical formulas, and comparison with the original density-based approach can be found at [https://github.com/zmaroti/ancIBDfiltration/blob/main/Method_description_and_performance_analysis.pdf].

In our dataset we included 8 Mesolithic and Neolithic samples (dated between 6246-9465 BP) to control our IBD filtration methodology. We assumed that any shared IBDs between these old samples and the coanalysed ancient or present-day data would represent false positive IBD segments. Table S9 contains the number of false positive IBDs that were shared between the 8 Mesolithic-Neolithic samples and the remaining 330 Balkan and West Eurasian-related samples.

The 4 cM IBD length threshold resulted in excessive false positive IBDs (∼20% for density and ∼8% for the score method) even with the score-based methodology (Table S9). The 5 cM data contained 61,381 raw IBD fragments between the 338 individuals, while only 17 out of the true 7,507 IBDs were between the Neolithic samples and the much more recent Balkan individuals (Table S9, S10) indicating about 0.65% of false positive IBDs at this threshold length. Based on these results, we used the distribution of the 5 cM raw IBDs to calculate the marker scores – our rationale is that the distribution of smaller fragments indicates the low confidence regions with better spatial accuracy leading to stricter IBD filtration. The low False Positive (FP) rate presented in Table S9 suggests that the vast majority of the identified IBDs above 5 cM length represent true positive matches, while the lower IBD length threshold identifies older connections between closely related populations in our datasets, which mirror our *qpAdm*, PCA and uniparental marker analyses. Accordingly, we summarised the shared IBD fragments between the analysed individuals based on the > 5, 6, 7, 8, 12 and 16 cM IBD length thresholds in Table S11.

In the downstream analyses and IBD network visualisation we only used the IBD data based on the 6 cM and 8 cM long IBD fragment thresholds. To investigate the extent of IBD sharing between the analysed populations we also calculated group statistics for the different IBD metrics (n_IBD – count of shared IBD, max_IBD – maximum length of shared IBD, and sum_IBD – sum of shared IBD) based on the 6 and 8cM IBD length data (Tables S12-S17). To account for the different number of individuals in the different groups we calculated a normalisation factor based on the number of possible combinations between the individuals to adjust the IBD statistics. Accordingly, within a group there are N*(N-1)/2 potential combinations between pairs of different individuals (where N is the number of individuals), while in case of the intra group scenario, there are N_1_*N_2_ sample pair combinations (where N_1_ and N_2_ stands for the number of individuals in group 1 and group 2). In Tables S12-S17 we included all the relevant information, namely: the number of individuals in group1 and 2 (count1, count2); the normalisation factor (norm); whether the relatedness is within or between groups (type); the given IBD statistic (n_IBD, max_IBD, sum_IBD) score that is being analysed (score); the IBD length threshold the data was based on (len_IBD); the number of individuals sharing IBD between group 1 and 2 (N); the mean of the IBD statistics (mean), that is calculated based on the IBD statistics between *only* the individuals that share IBD; the dispersion of the IBD statistics (dispersion); and lastly the normalised sum of the IBD statistics (norm_sum) that holds the sum of the analysed IBD statistic divided by the normalisation factor.

### Coalescence rate and Ne analysis

The Colate input files were precomputed using the “mode make_tmp” option with the provided SGDP “half_ne_fixed” mutation ages for all data^83^. We used the publicly available BAM files for the ancient DNA samples from Medieval-Post-Medieval Albania (I14686, I14687, I15707, I13839, I14622). For the high coverage data of the present-day Albanian dataset, we used the accessible regions of the GRCh37 reference as the target mask and the associated BCF files. The pairwise coalescence rates were estimated for all sample combinations using “mode mut” on the precompiled “colate.in” input files using the 20 bootstraps, with sample chronologies-epochs defined by bins ‘1,5,0.4’, with the appropriate relative ages (“target_age”, “reference_age”) of samples (based on the AADR annotation file and a uniform 5 year for all modern samples). The output was imported in R (version 4.4.2) by the relater package (version relater_0.0.0.9000) and visualised in ggplot2 (version 3.5.1).

### hapROH

The results of the abovementioned Colate analysis indicated that the founding Albanian population was small. To test whether the proto-Albanians were an endogamous or bottlenecked population with reduced genetic diversity, we used hapROH^85^ [https://pypi.org/project/hapROH/], which detects ROH due to recent common parental ancestry. We inferred ROHs using the pseudo haploid 1240K dataset and hapROH for all ancient and present-day individuals with >400K genotyped markers from the territory of modern Albania. The identified ROHs were plotted and used for Ne estimation using the maximum likelihood MLE method of the hapROH tool.

### Data visualisation

Graph visualization and data manipulation was performed in R (version 4.4.1). The **iGraph** R package (version 2.0.3) was used to represent and filter the IBD data as an undirected graph where each individual was represented as a node. The weighted edges of the graph were used to represent the sum of shared IBD between any two individuals. The Fruchterman Reingold [https://doi.org/10.1002/spe.4380211102] algorithm implemented in the **qgraph** R package (version 1.9.8) was used to build the IBD network layout. For consistent graph layout the 5cM IBD length data was used in the FR algorithm and the resulting graph was post filtered for max_IBD>6.0, max_IBD>8.0 and/or other population or date based criteria. The resulting IBD graphs were plotted by the **ggplot2** R package (version 3.5.1). All haplogroup plots were also created using package ggplot2. To generate vertical stacked bar plots, we additionally used the forcats package. To combine plots, we used Adobe Illustrator v. 27.4. We generated maps using the QGIS Geographic Information System, QGIS Association (http://www.qgis.org), SimpleMappr. The inset in Fig. 1B. and the entirety of Fig. 5 were created using base maps from ArcGIS® software by Esri. ArcGIS® and ArcMap™ are the intellectual property of Esri and are used herein under license. Copyright © Esri. All rights reserved. For more information about Esri® software, please visit www.esri.com.

## Supporting information

Supplementary Text and Images

## Acknowledgements

The authors are grateful to Joscha Gretzinger (Max Planck Institute for Evolutionary Anthropology, Leipzig) for undertaking the Mobest analyses. L.R.D. thanks Andreas Kyropoulos and Leonidas Embirikos for extensive discussions on Albanian history and linguistics. L.R.D. thanks Alexandros Spanos and Leo R. Cooper for assistance in compiling part of the ancient Greek Y-chromosome dataset (Table S23). The authors thank Ted Kandell (Open Genomes) and Milan Rajevac (Open Genomes, the Society of Serbian Genealogists “Poreklo”) for providing detailed Y-chromosome haplogroup determinations for part of the examined dataset.

## Author contributions

Conceptualisation: L.R.D. Data curation: L.R.D., A.L., A.A., Z.M., D.W. Formal Analysis: L.R.D., A.A., Z.M., D.W., A.H. Funding acquisition: L.R.D. Investigation: L.R.D. Methodology: L.R.D., A.L., A.A., Z.M., D.W., A.H. Project Administration: L.R.D. Validation: L.R.D., A.L., A.A., Z.M., D.W., A.H. Visualisation: L.R.D., A.L., A.A., Z.M., D.W., A.H. Writing – original draft: L.R.D. Writing – review and editing. L.R.D., A.L., A.A., Z.M., G.B., A.M., I.M., D.W., B.D.J., A.H.

## Data availability

All data needed to evaluate the conclusions in the paper are present in the paper and the Supplementary Materials. However, after consulting with the 74 ethnic Albanian participants, their sequencing data will not be deposited in a public repository and will be made available upon request to A.L. and G.B. Permission to use the data will be sought from each individual participant.

