## Supplementary Text and Images for "Ancient DNA reveals the origins of the Albanians"

**1. *qpAdm* Admixture modelling**

*1.1. Ancestry modelling for Bronze Age, Iron Age, Roman, and post-Roman Balkan populations, using Mesolithic-Neolithic sources*

We first sought to characterise the distal genetic makeup of all the examined samples (Table S3), using an Mesolithic-Neolithic model where we rotated among Yamnaya_Samara, Iran_N, Levant_N, Albania_NChL, Anatolia_N, Iron_Gates_HG. Due to the great geographic and temporal diversity of the samples, we used two different farmer-related metapopulations (Albania_NChL, Anatolia_N), as some groups differentially preferred either source, and the model failed in their absence. Even so, a subset of our samples (all samples from Bulgaria, Greece, and North Macedonia) required a different Neolithic source (Bulgaria_N, Greece_Alepotrypa_EN, North_Macedonia_N) for the model to attain significance, likely due to variation not captured by our standard farmer-related sources (Albania_NChL, Anatolia_N). The fixed references included Cameroon_SMA, Morocco_Iberomaurusian, ISR_Natufian_EpiP, RUS_West_Siberia_HG, Russia_Karelia_HG, WHG, TUR_Pinarbasi_EpiP, Russia_Boisman_MN. We should note that the proportion of Iran_N-related ancestry might be underestimated for some of the studied populations (e.g. Greece_Manika_Helladic_EBA.SG, Greece_BA_Mycenaean), as the Neolithic populations contributing to the bulk of their genome were already admixed with a Iran_N/CHG-related source^1,2^.

A group of individuals from Post-Medieval Barç (Albania_Barc_Post_Medieval_Roma) formed a cluster that was distinct from both ancient and present-day Balkan peoples (Fig. 4A), and their uniparental markers were typical of present-day Roma people (Table S22). To model their ancestry, we used the distal Yamnaya_Samara, Iran_N, Anatolia_N, Levant_N, CHG, Iron_Gates_HG, Indian_GreatAndaman_100BP.SG, with fixed references being Cameroon_SMA, Mar_Taforalt_EpiP, Mongolia_North_N, ISR_Natufian_EpiP, Russia_West_Siberia_HG, Russia_Karelia_HG, WHG, TUR_Pinarbasi_EpiP, Russia_Boisman_MN. The resulting model confirmed significant ancestry ultimately from South Asia and the Caucasus (Table S4), in agreement with the known history of the Roma people^3^ and our ADMIXTURE analyses (Figs. S2-S3).

*1.2. Ancestry modelling for the Early Bronze Age sample from* *Çinamak, Albania*

To better resolve the more recent ancestry of Albania_Çinamak_EBA we rotated among roughly contemporary sources (Russia_Steppe_Catacomb, Bulgaria_Boyanovo_EBA, Germany_CordedWare, Poland_Southeast_CordedWare.SG, Serbia_EBA_Yamnaya, Albania_NChL, Czech_CordedWare, Iron_Gates_HG), with the following base set of references: Cameroon_SMA, Yamnaya_Samara, Turkey_Arslantepe_EBA, Iran_N, Anatolia_N, Mar_Taforalt_EpiP, ISR_Natufian_EpiP, Levant_N, Russia_West_Siberia_HG, Russia_Karelia_HG, WHG, TUR_Pinarbasi_EpiP, Russia_Boisman_MN. Sources from the early Balkans (Bulgaria_EBA_Yamnaya, Serbia_EBA_Yamnaya) and individuals directly from the steppe (Russia_Steppe_Catacomb) were preferred over Corded Ware groups, suggesting that the latter did not contribute to the ancestry of the earliest steppe populations of the Balkans. The proportions of steppe ancestry under this model (70-74%; Table S4) are consistent with the one using earlier sources from the Mesolithic and Neolithic (66-70%; Table S3).

*1.3. Ancestry modelling for the Middle Bronze Age sample from Shkrel, Albania*

We rotated among geographically and temporally contemporaneous metapopulations (Albania_Çinamak_EBA, Croatia_MBA_Cetina, Croatia_BA_Bogomolje, Serbia_Mokrin_EBA_Maros, Greece_Logkas_MBA.SG) and our base set of references for post-Neolithic populations (Cameroon_SMA, Yamnaya_Samara, Turkey_Arslantepe_EBA, Iran_N, Anatolia_N, Mar_Taforalt_EpiP, ISR_Natufian_EpiP, Levant_N, Russia_West_Siberia_HG, Russia_Karelia_HG, WHG, TUR_Pinarbasi_EpiP, Russia_Boisman_MN) (Table S4). The sample could be adequately modelled as deriving its entire ancestry from sources originating from the Middle Bronze Age West Balkans and northern Greece (Croatia_MBA_Cetina, Croatia_BA_Bogomolje, Greece_Logkas_MBA.SG; Table S4) with a lower contribution of steppe ancestry (roughly 40%) compared to the preceding Early Bronze Age individual from Çinamak (66-70%; Table S4), indicating the homogenisation of European Neolithic-Chalcolithic ancestry.

*1.4. Ancestry modelling for the* *Late Bronze Age and Iron Age samples from Çinamak, Albania*

By rotating among contemporaneous metapopulations across the Balkans (Albania_MBA, Croatia_MBA_Cetina, Montenegro_MLBA, Serbia_Mokrin_EBA_Maros, North_Macedonia_BA, Bulgaria_EIA, Greece_BA_Mycenaean), the Late Bronze Age and Iron Age samples from Çinamak, Albania could be effectively modelled as deriving their entire ancestry from the neighbouring population of North_Macedonia_BA (Table S4). Two-way models combining a local West Balkan source (Albania_MBA, Croatia_MBA_Cetina, Montenegro_MLBA) and a southeastern Balkan source (Bulgaria_EIA, Greece_BA_Mycenaean) were also adequate (Table S4), suggesting subtle ancestry shifts from the preceding Middle Bronze Age.

We should note that we excluded individuals I18723, I18721, I18719 from Bezdanjača Cave in Croatia, who were archaeologically dated to the Bronze Age (1,500-800 BCE), and were interpreted in previous studies as outliers compared to the contemporary population of Croatia^4,5^. We argue that, based on insights from the PCA and their uniparental markers, these outlying individuals likely date to post-Medieval times. Individuals I18721 and I18719 are assigned to haplogroup I2-M423>I-CTS10228>I-Y3120 (Table S22), which is associated with the Slavic expansion toward southern Europe during the Migration Period and has experienced major founder effects in the South Slavic population of the Balkans^6–8^. This haplogroup is unlikely to have entered the Western Balkans in the Bronze Age, as our extensive haplogroup dataset shows that, as expected, subclades downstream of I2-M423 appear in the region primarily during the Migration Period (Fig. 10). There were occasional migrants from the Balto-Slavic and steppe nomadic world to southern Europe in the Iron Age, such as the two mercenaries from the Battle of Himera in Sicily^9^, but the Bezdanjača Cave outliers are unlikely to represent BA migrants from northern Europe, as the mitochondrial haplogroup of individual I18719 (HV0a1a1b), has a TMRCA of 225 years before present with a person from present-day Germany^10^. Furthermore, previous archaeological studies have recovered the remains of post-BA individuals in Bezdanjača Cave, which were radiocarbon-dated to the 17^th^ century CE and World War II^11^. Another sample that might be misdated is I13170 from Velika Gruda in Montenegro. Without having been radiocarbon-dated, this sample has been archaeologically assigned to the Iron Age (800-400 BCE), but it has been excluded from our analyses as it clustered with present-day South Slavs^5^.

*1.5. Ancestry modelling for Balkan-shifted Iron Age outliers from Himera, Sicily*

Two mercenaries from the Battle of Himera in Sicily (Himera_Balkan_outliers) cluster close to West Balkan populations in the Bronze Age-Iron Age PCA (Fig. 3A, B). To test whether this clustering involves shared ancestry or is instead the result of projection artefacts, we rotated among a broad range of Balkan and Alpine populations (Albania_BA_IA, Slovenia_EIA. Montenegro_MLBA, Serbia_Mokrin_EBA_Maros, Serbia_LBA, North_Macedonia_BA, Bulgaria_EIA, Greece_BA_Mycenaean) (Table S4). Only two-way models are successful for Himera_Balkan_outliers, suggesting that they either originate from a population that has not yet been sampled, or represent admixed individuals between two different populations. In distal models, Himera_Balkan_outliers are characterised by high IGHG-related ancestry (9%), which in Bronze and Iron Age Balkan populations is found in similar proportions only among samples from Early Bronze Age Serbia (11%; Table S3). Indeed, most two-way proximate models for Himera_Balkan_outliers require a source from Serbia (Serbia_Mokrin_EBA_Maros, Serbia_LBA) for the bulk of their ancestry (65-80%), with additional admixture (20-25%) from a southern proxy (North_Macedonia_BA, Bulgaria_EIA, Greece_BA_Mycenaean) (Table S4). Based on the above, Himera_Balkan_outliers likely originate from an inland location in the Central Balkans, at the junction where “Illyrian” and “Daco-Thracian” languages are thought to have been spoken (Fig. S16). Our hypothesis is further supported by IBD-sharing, which revealed connections of the Himera outliers with both West and East Balkan populations (Table S17).

*1.6. Ancestry modelling for Medieval, and Post-Roman populations within the East European cline*

Our rotated model included Roman era Balkan populations (Serbia_Roman_Svilos, West_Anatolia_Roman, Croatia_Trogir_Medieval, Croatia_Roman_Gardun, Montenegro_Doclea_Roman) and a northeast-European-related source (CEE_Medieval), with a base set of reference populations (Cameroon_SMA, Yamnaya_Samara, Turkey_Arslantepe_EBA, CHG, Iran_N, Anatolia_N, Iron_Gates_HG, Mar_Taforalt_EpiP, ISR_Natufian_EpiP, Levant_N, Russia_West_Siberia_HG, Russia_Karelia_HG, WHG, 'TUR_Pinarbasi_EpiP', Russia_Boisman_MN). Northeast-European-related admixture follows a geographical pattern among Balkan populations, with samples from Medieval-Post-Medieval Croatia scoring the highest proportions of said ancestry (75-86%), Medieval-Post-Medieval Serbia and Montenegro with slightly lower levels (ca. 65%), and North_Macedonia with the lowest (ca. 40%) (Table S4).

*1.7. Proximate ancestry modelling for Medieval, Post-Medieval, and newly sequenced present-day samples from Albania*

Based on the PCA (Fig. 4), it is evident that the Roman and Migration Periods witnessed profound demographic shifts in the ancestry of Balkan populations. Previous studies have suggested that the overall genetic structure of samples from Albania remained remarkably homogenous across time^5,12^. However, they used Mesolithic and Neolithic sources (Anatolia_N, CHG, EHG)^5,12^, which are not helpful in understanding the proximate populations that contributed to the formation of the Medieval, Post-Medieval, and present-day peoples of Albania. A more recent study^13^, employed a *qpAdm* model for modern Albanian samples from the HO dataset (only 600k SNPs) using a local source (Albania_BA_IA), as well as generalised sources for individuals rich in Anatolian-related ancestry, and CEE_Medieval. Our aim is to use *qpAdm* models to identify the likeliest populations who contributed to the ancestry of samples from Albania in each of the three periods examined (Medieval, Post-Medieval, present-day), guided by our findings from the PCA, uniparental markers, and ancIBD.

In terms of sample selection for the *qpAdm* analyses described below, we should note that among the samples from Albania, individual I13834 from Barç (Southeast, Korça Basin), radiocarbon dated to Post-Medieval times (1452-1619 calCE (385±15 BP, PSUAMS-8300)^5,12^, is intriguing. The sample clusters with Bronze Age and Iron Age samples from Albania (Fig. 5) and shares IBD segments solely with samples from that period (Fig. 6A) and accordingly lacks any of the Iran N-related ancestry present in Medieval, Post-Medieval, and present-day samples from Albania (Table S3). In contrast, sample I13839 from neighbouring Shtikë (Southeastern, Kolonja Plateau), which is radiocarbon dated to 889-989 CE^5,12^ derives part of his ancestry from an Iran N-related source (Table S3), as expected. Based on the above, individual I13834 from Barç either descends from a population that maintained an unadmixed profile for more than 1,600 years, or it might represent a case of the freshwater reservoir effect, which can cause errors in carbon dating of samples^14^. Due to these dating uncertainties, we tentatively assigned individual I13834 at least to the earlier Medieval population on the PCA (Fig. 4), and excluded him from all proximate admixture analyses.

To characterise the ancestry of Albania_Medieval (I13839, I14622), we rotated among a local Albanian Bronze and Iron Age source (Albania_BA_IA, assuming some degree of local continuity), an Eastern Balkan source (Bulgaria_EIA), and populations with full (East_Anatolia_BA_IA, Southeast_Anatolia_Roman) or predominant Anatolian ancestry [Balkans_Anatolian_o (samples admixed with Anatolians, North Africans, and Levantines^13^), West_Anatolia_Roman]. Only two-and-three-way models were feasible (Table S4), which recovered Albania_Medieval as deriving most (68-84%) of their ancestry from Albania_BA_IA, with additional admixture from either an Anatolian-admixed (16-32%) or Bulgaria_EIA-admixed (32%) source. While we cannot identify whether Anatolian or Bulgaria-EIA-related ancestry is more likely, the latter receives lower p-values (p = 0.067073), and higher Standard Errors (0.101). The proportion of Anatolian-related ancestry is lower (16-21%) if an East Anatolian proxy is used, and higher (32%) in West Anatolian Romans and Anatolian-admixed samples from the Roman Balkans are employed as sources.

We next rotated using our unadmixed East European proxy (CEE_Medieval) and slightly earlier Balkan Roman populations that cluster close to Albania_Medieval on the PCA (Montenegro_Doclea_Roman, Serbia_Roman_Naissus; Fig. 4A) (Table S4). To test linguistic theories which postulate a Daco-Thracian origin of Abanian^15–17^, we also added individuals from Roman Serbia (Serbia_Roman_Thracian_profile) that cluster with Bronze and Iron Age samples from the southeastern Balkans (Bulgaria_EIA and Greece_BA_Mycenaean; Fig. 4). A one-way Roman period West Balkan model (100% Montenegro_Doclea_Roman ) was adequate, suggesting that Daco-Thracian-speaking populations either did not contribute ancestry to Albania_Medieval, or their autosomal profile was Central-West-Balkan-like. Furthermore, Albania_Medieval lacks East European-related ancestry entirely (based on our *qpAdm* models and the absence of IBD matches), demonstrating that the Slavic migrations that transformed the genetic and linguistic landscape of the Balkans in the Early Middle Ages did not affect the sampled individuals from 8-9^th^ CE Albania.

To reveal the proximal ancestry of samples from Post-Medieval Bardhoc and Pazhok, we rotated using a local source (Albania_Medieval), an unadmixed northeastern European proxy (CEE_Medieval), and a suite of Medieval, and Post-Medieval Balkan metapopulations that fall within the East European cline of the PCA (Croatia_Mdv_PostMdv, Serbia_Mdv_PostMdv, Montenegro_Medieval; Fig. 4; Table S4). We recover Albania_Bardhoc_Post_Medieval as descending 100% from Albania_Medieval, which demonstrates the persistence of populations lacking East European-related admixture even in the 15^th^ century CE (Table S4). Two Post-Medieval outliers (Albania_Bardhoc_Post_Medieval_o, Albania_ Post_Medieval_Pazhok_o) exhibiting a shift towards the East European cline on the PCA (Fig. 4B), are characterised by moderate levels of East European-related admixture (21-32%) than their contemporaries from Bardhoc (Table S4). This finding suggests that currently unsampled populations with moderate levels of East European-related admixture already existed in the territory of present-day Albania, or the process of admixture with Slavic-speaking groups was ongoing during the 15^th^ century.

Our next step was to model the ancestry of the newly sequenced, present-day Albanian individuals. To incorporate West Balkan genetic variation that might not captured by the two Medieval samples from Shtikë and Kënetë, and to maximise the number of SNPs used, we created a new category named as *West_Balkan_Roman_Medieval*, which includes Albania_Medieval and Montenegro_Doclea_Roman – a geographically adjacent individual that plots on a similar position on the PCA (Fig. 4) and shares a large proportion of West Balkan Iron Age ancestry (Table S4). We also added a Roman era Central Balkan proxy (Serbia_Roman_Naissus) from the city of Naissus (which has been proposed as a proto-Albanian toponym^18,19^), as well as a suite of East European-related proxies (CEE_Medieval Montenegro_Medieval, Croatia_Mdv_PMdv).

We recover significant variation in East-European-related admixture in present-day Albanians. Depending on whether a largely unadmixed (CEE_Medieval) or paleo-Balkan-admixed (Montenegro_Medieval, Croatia_Mdv_PMdv) East European-related proxy is used, we recover present-day Albanians as scoring 4-16% or 8-32% of this ancestry, respectively. Furthermore, while many individuals can be modelled as 100% West_Balkan_Roman_Medieval, indicating remarkable continuity from Roman and Early Medieval times, some outliers from TN (TN1, TN2, TN7, TN10, TN11) and GNW (GNW5, GNW6, GNW7) score 23-50% CEE_Medieval or 36-73% Montenegro_Medieval – an expected finding from zones of intercultural contact and exchange. Overall, the average proportion of East European-related ancestry in our present-day Albanian dataset is 10% or 20%, depending on whether CEE_Medieval or Montenegro_Medieval is used as a proxy.

*1.8. Testing for sex bias in East-European-related admixed populations from Albania*

A significant knowledge gap in our understanding of the introduction of East-European-related ancestry in Medieval, post-Medieval, and present-day populations from Albania involves its social and gender dynamics. We do not know whether the individuals or population(s) with elevated East-European-related ancestry who admixed with the inhabitants of Albania were primarily male, female, or had an even sex ratio. To test for the presence of sex bias, we aimed to compare ancestry proportions on the X-chromosome with those of the autosomes. Our model included two potentially local sources (Albania_Medieval, Serbia_Roman_Naissus), a suite of northeast-European-related proxies (CEE_Medieval, Montenegro_Medieval, Serbia_Medieval_Post_Medieval, Croatia_Medieval_Post_Medieval), with a base set of reference populations (Cameroon_SMA, Yamnaya_Samara, Turkey_Arslantepe_EBA, CHG, Iran_N, Anatolia_N, Iron_Gates_HG, Mar_Taforalt_EpiP, ISR_Natufian_EpiP, Levant_N, Russia_West_Siberia_HG, Russia_Karelia_HG, WHG, 'TUR_Pinarbasi_EpiP', Russia_Boisman_MN). Groups with less than 15% northeast-European-related ancestry were excluded from our analysis, namely Albania_Bardhoc_PostMdv (0%) and TS (4-11%) (Table S4).

The results of our *qpAdm* analysis for sex bias are inconclusive. Central Ghegs (GC) are characterised by 100% Albania_Medieval ancestry on their X-chromosome, which may indicate a high proportion of northeast-European-related males admixing with this population. However, the autosomal ancestry of GC is also overwhelmingly local (85-100%), which may also lead to fully local ancestry on the X-chromosome in the absence of sex bias. In two other groups (Albania_Bardhoc_PostMdv_o, GS), northeast-European-related sources and local sources are equivocal (100% of either type of ancestry, only differing by p-value support). In all other groups (Albania_Pazhok_PostMdv_o, GNE, GNW, TDA, TN, TW), the ancestry on the X-chromosome is predominantly or fully northeast-European-related (Table S4), suggesting an overabundance of females introducing this type of ancestry into the indigenous populations of Albania. However, we caution that the comparative accuracy of these results is severely restricted by the low number of markers on the X-chromosome (4.6k SNPs), compared to the autosomes (1240k SNPs), which lead to very high standard errors (Table S4). Furthermore, the *qpAdm* model that yielded the highest p-value is not necessarily the best-fitting model, while the choice of left and right populations also affects the results^20^.

Due to the inconclusive nature of the X-chromosome *qpAdm* results, we sought to investigate the possibility of sex bias in the uniparental markers (Y-chromosome, mitochondrial DNA) of present-day Albanians. The results are presented in the main text.

*1.9. Modelling the ancestry of E-V13 and J2b-L283 individuals from the Roman Balkans*

Despite the high frequency of E-V13 in present-day Balkans, the origins of this haplogroup have remained unresolved. To this end, we modelled 17 E-V13 individuals from the Roman Balkans and one individual from Avar era Hungary, using an earlier eastern Balkan source rich in this haplogroup (Bulgaria_EIA), a merged West Balkan source (*West_Balkans_BA_IA*, comprising Albania_BA_IA, Croatia_BA_Bogomolje, Croatia_MBA_Cetina, Croatia_BA, Croatia_Bezdanjača_BA, Croatia_MBA, Croatia_EIA, Croatia_IA, Montenegro_MLBA), as well as Central Balkan (Serbia_Mokrin_EBA_Maros), southeast Balkan (Greece_Mycenaean_BA), Anatolian (West_Anatolia_Roman, Southeast_Anatolia_Roman), and Scythian-related (Serbia_Steppe_Nomad) sources.

We recover that 72% of all E-V13 individuals derive 24-100% of their ancestry from a Bulgaria_EIA/Greece_Mycenean_BA source, often with equal statistical support (Table S4). Given that E-V13 has never been found in BA-IA Greece, despite dense sampling (Fig. 10), we consider Bulgaria_EIA a more likely source for E-V13 in the Roman and Avar populations. Indeed, D-statistics and IBD matching suggest that Bulgaria_EIA is a more likely source of E-V13 for at least some of these individuals (Fig. S6; Table S10) and pinpoint the “Daco-Thracians” as the ethnolinguistic group that may have disseminated a significant proportion of the E-V13 diversity that we observe today in the Balkans and elsewhere. Our hypothesis is further supported by the fact that several of the E-V13 individuals modelled as 100% Bulgaria_EIA (R6756, I15504, I15518, I15490, I15495, I15554, I16750) plot within the same PCA space with Thracians from Bulgaria (Fig. 4A).

Despite the abovementioned pattern, we should note that E-V13 is not always associated with Bulgaria_EIA/Greece_Mycenean_BA-related sources, as some individuals descend fully or largely from West Balkan (R3659), Anatolian (I15513, I15525), and nomadic steppe (R3931) populations.

Haplogroup J2b-L283 is the West and Central Balkan equivalent of E-V13, with a frequency of 50-70% in this region during the BA-IA. Accordingly, all Roman era J2b-L283 individuals from the Balkans derive 50-100% of their ancestry from a West_Balkans_BA_IA or Serbia_Mokrin_EBA_Maros-related population (Fig. S17; Table S4).

**Supplementary Figures**

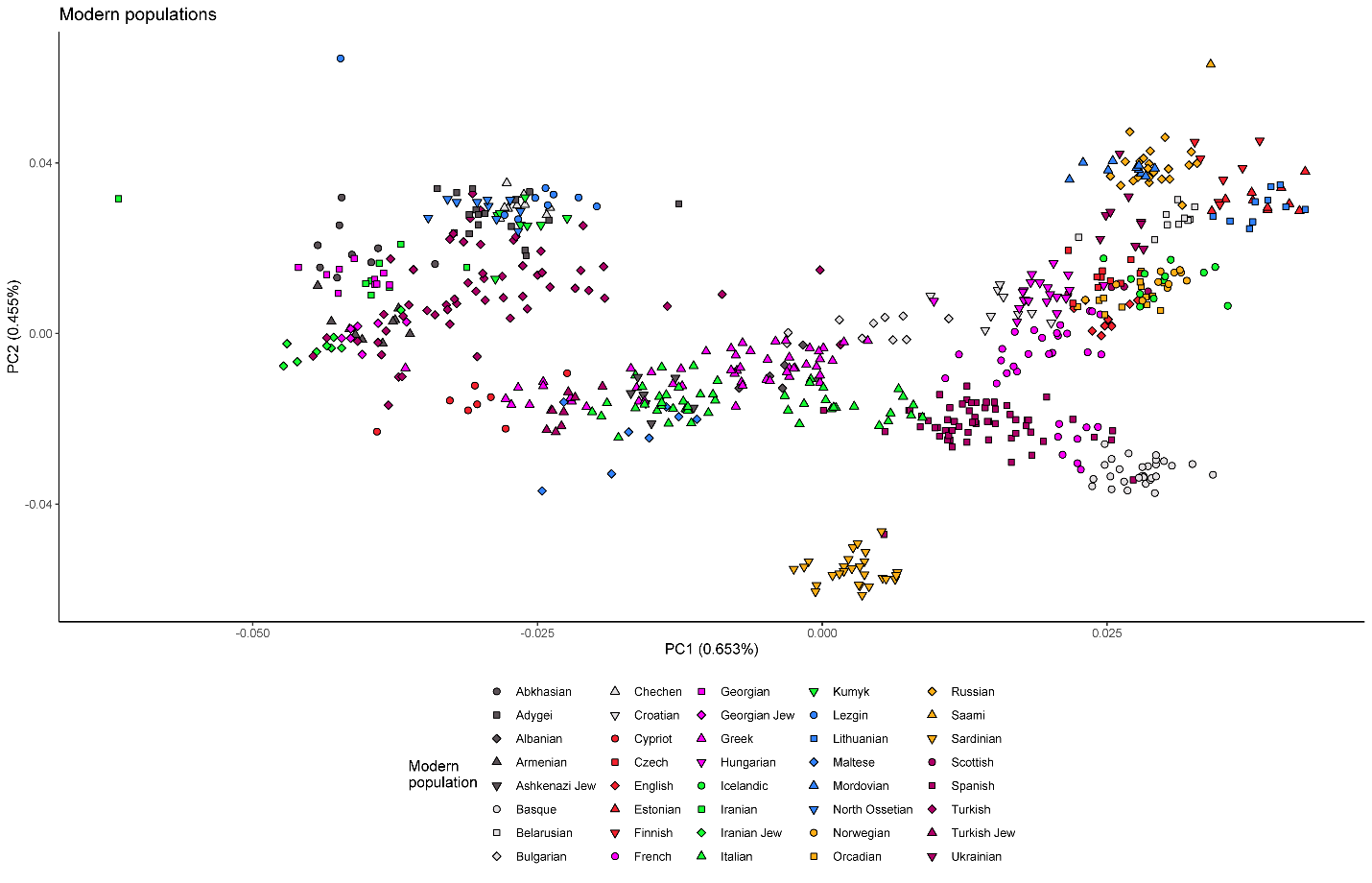

**Figure S1.** Principal Components Analysis on a subset of the HO and Reitsema et al. 2022 dataset, onto which the ancient individuals used in this study were projected upon.

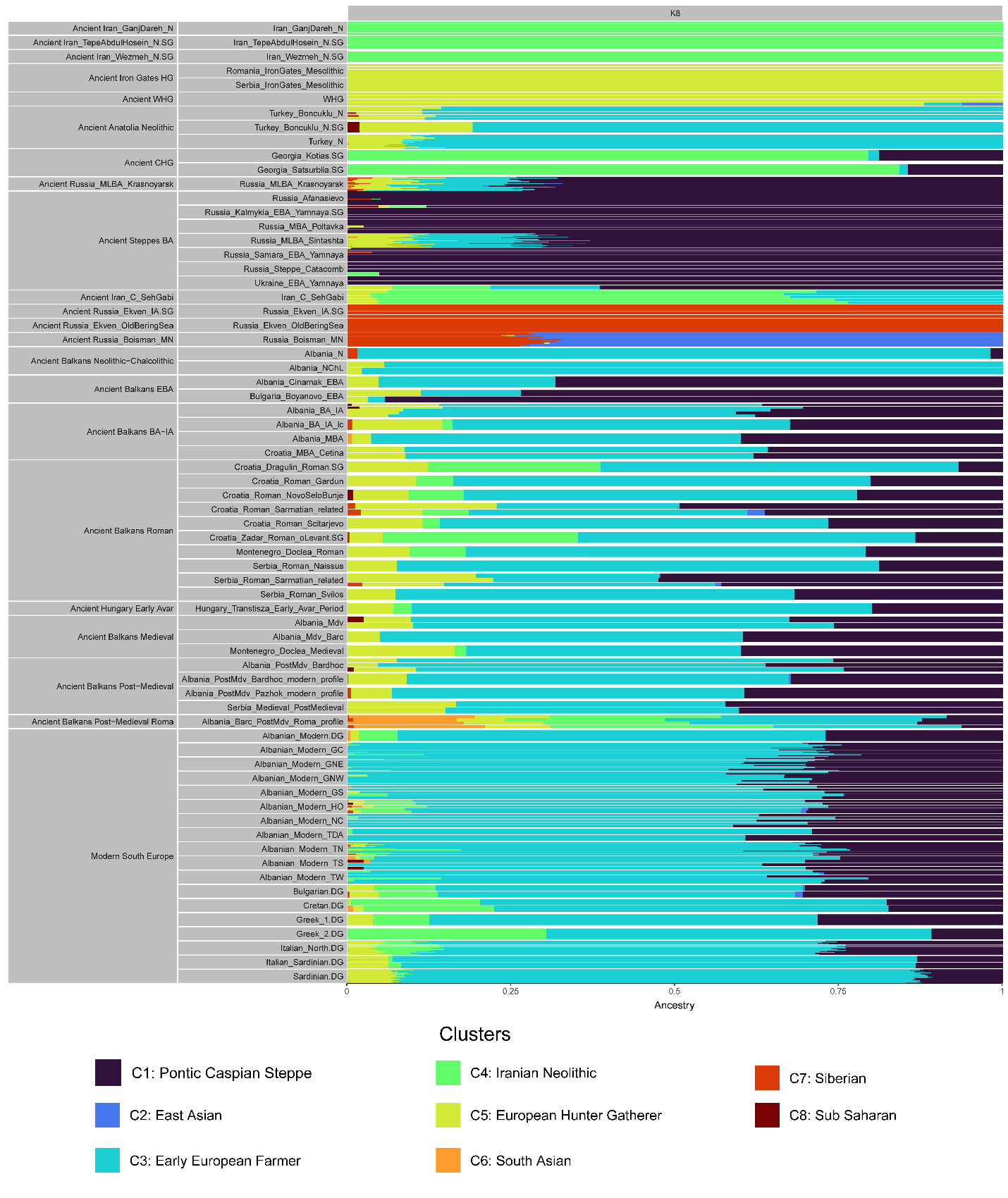

**Figure S2.** Bar graph illustrating the results of ADMIXTURE analysis, with an emphasis on the Balkans from the Mesolithic to the present-day era. Even though these results cannot be interpreted literally, as the clusters are inferred from both present-day and ancient DNA data, they broadly resemble our distal *qpAdm* analyses (Tables S3, S4), showing a series of major genetic shifts in the Balkans, especially during the Neolithic and Bronze Age, with the introduction of Yamnaya-related ancestry from the Pontic-Caspian steppe. Furthermore, several populations in the Roman era display high levels of West-Central Asian ancestry, and populations falling within the East European cline in the PCA (Fig. 4A) are characterised be higher proportions of European Mesolithic ancestry compared to preceding era. Significant levels of South Asian ancestry characterise a group of post-Medieval Albanian individuals interpreted as belonging to the Roma people based on insights from the PCA, *qpAdm*, and uniparental data.

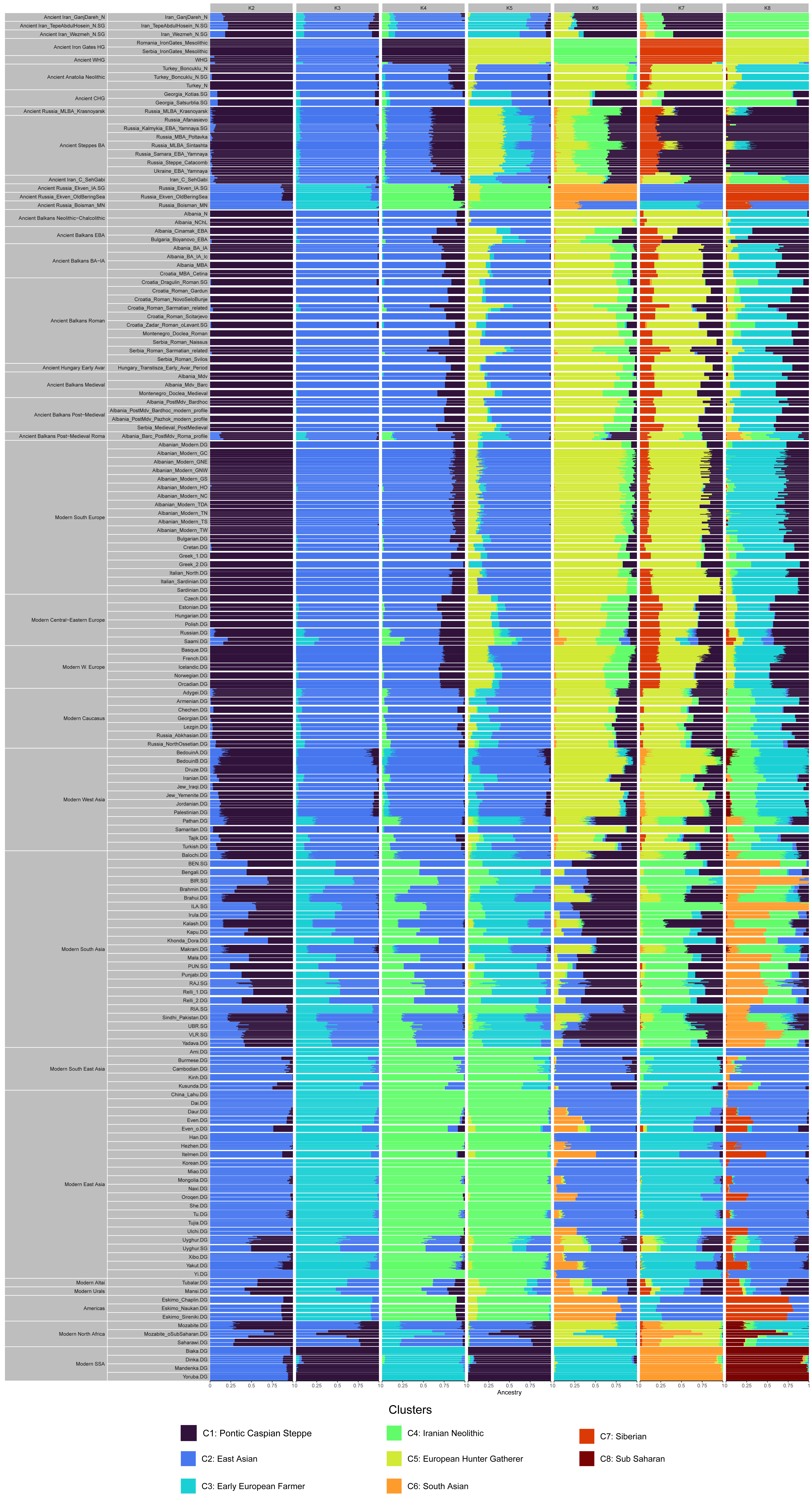

**Figure S3.** Full report of ADMIXTURE analysis on ancient and present-day individuals.

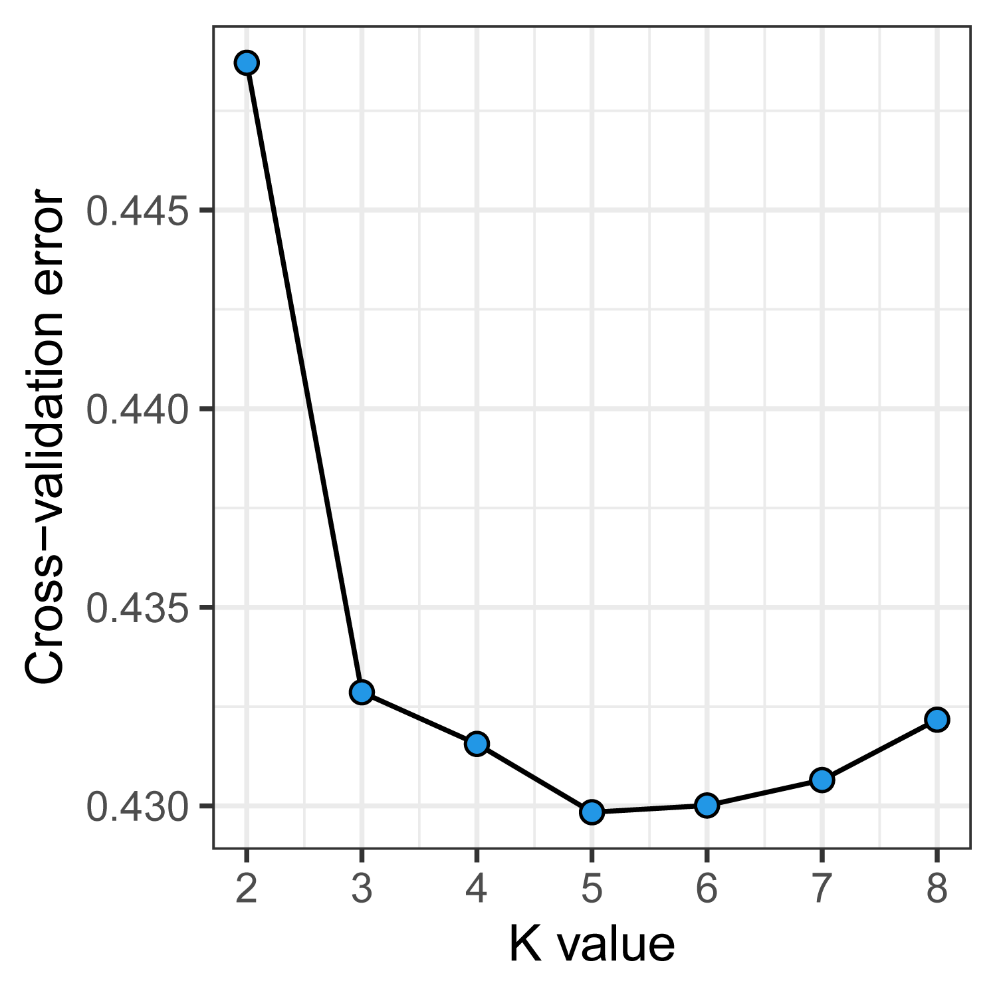

**Figure S4.** Cross-validation errors of ADMIXTURE analysis on ancient and present-day individuals.

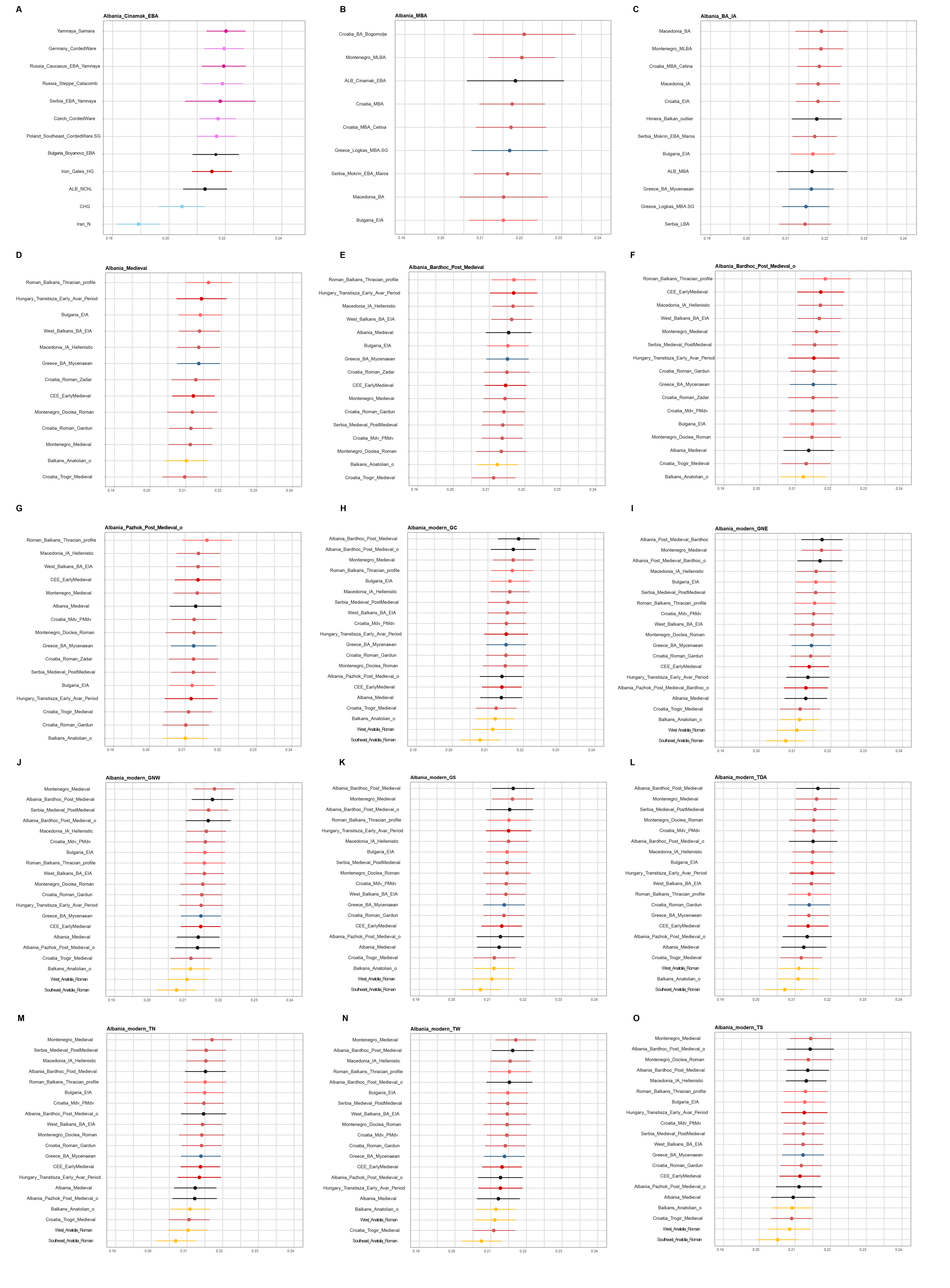

**Figure S5.** Outgroup *f3*-statistics panel of the form *f3(Cameroon_SMA; tested sample from Albania, other ancient population)* estimating shared genetic drift based on allele sharing. The further to the right the points are in each plot, the higher the allele sharing with the tested sample or metapopulation from Albania. Wider error bars indicate lower precision, as a result of a smaller number of available SNPs in the given pairwise combination.

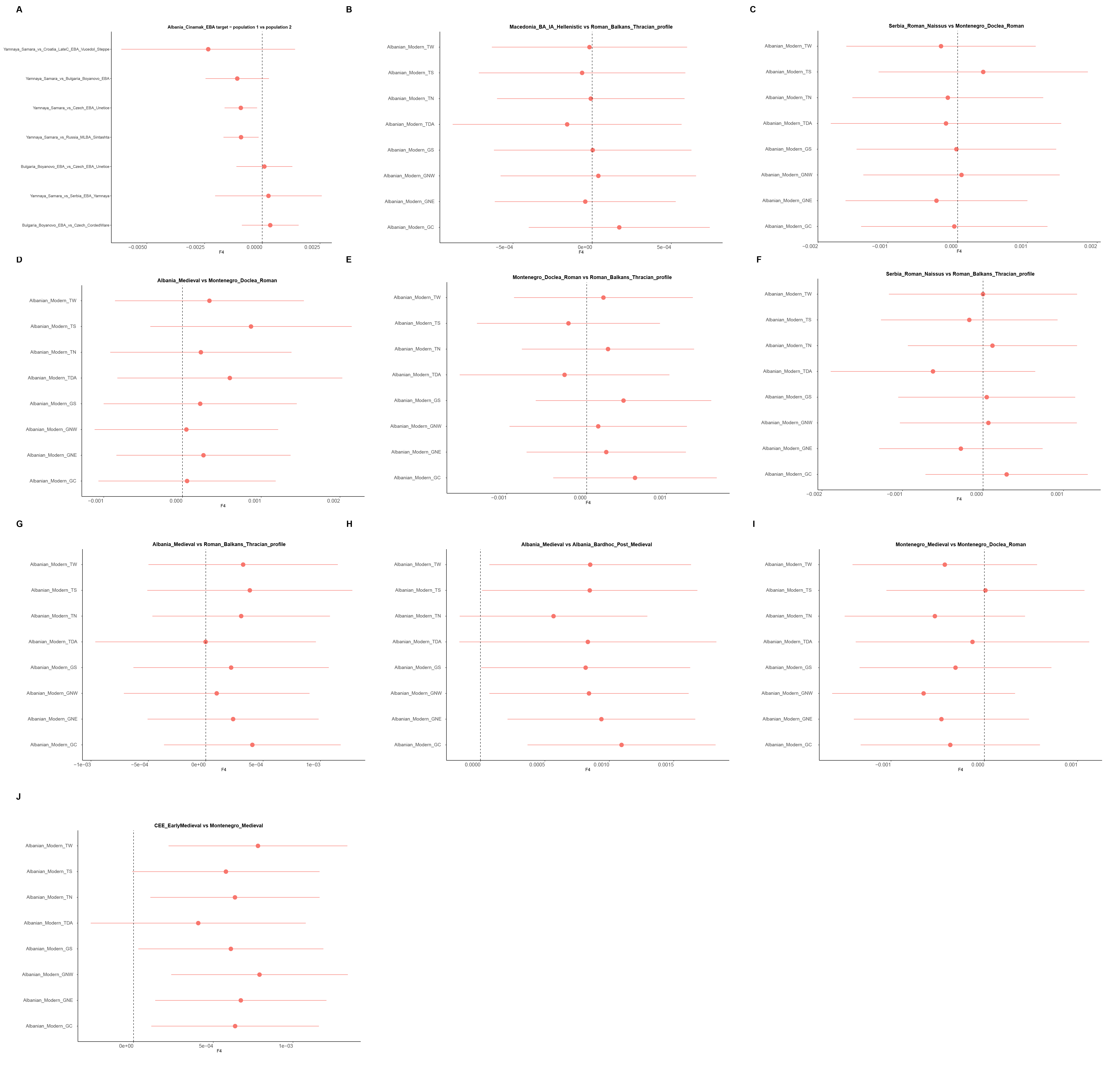

**Figure S6.** Outgroup *f4*-statistics of the form *f4(outgroup, population A; population B, population C)* estimating shared genetic affinity based on allele sharing.

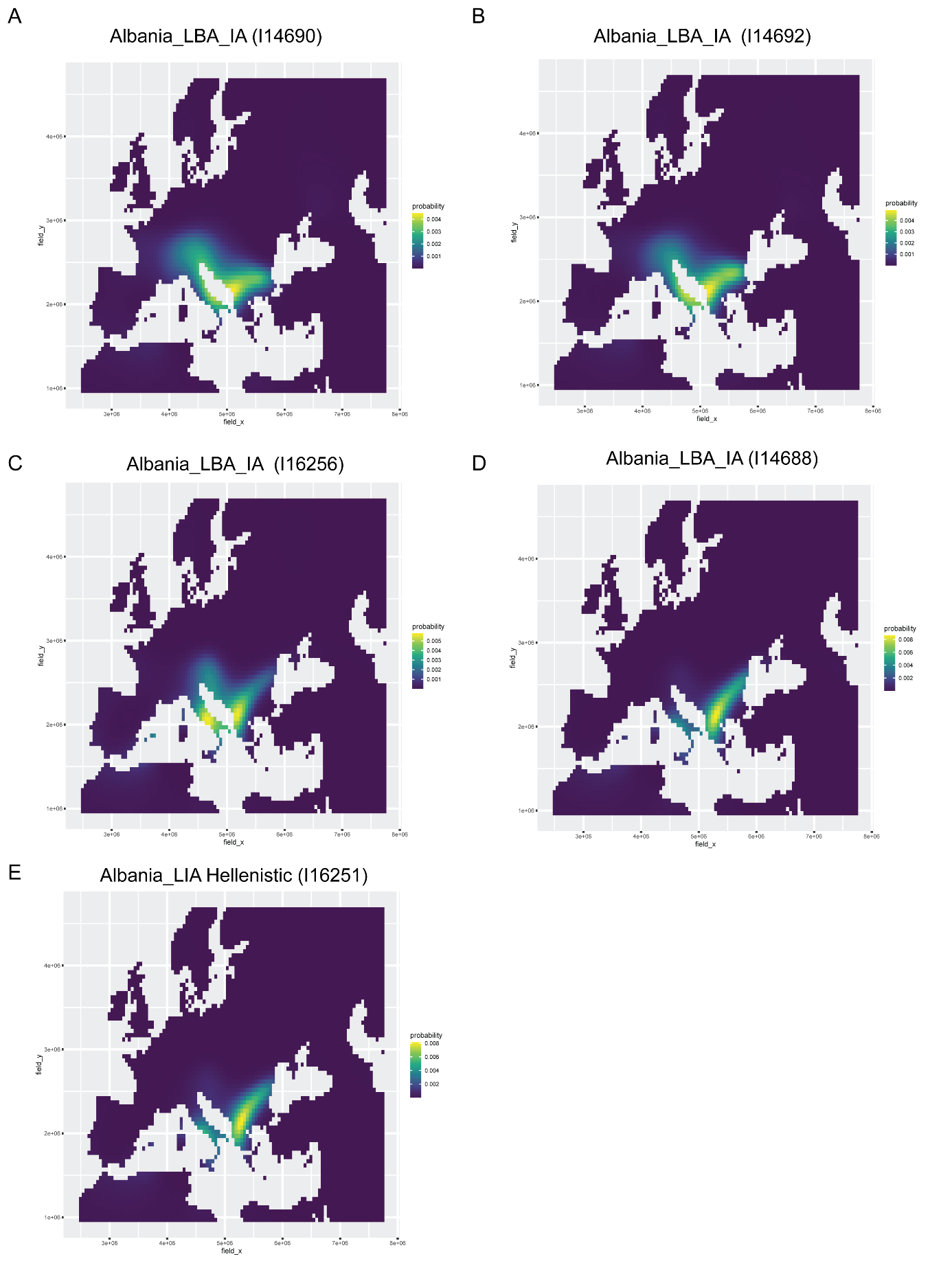

**Figure S7.** Mobest analysis of Late Bronze Age-Iron Age samples from the region of Albania, plotting the probability surface that identifies the highest genetic-geographical match at the mean date of the respective individual. The higher the probability surface (light yellow-green), the closer the genetic match. The latitude and longitude coordinates with the best fit for the examined individuals are provided in EPSG:3035 projection in Table S19.

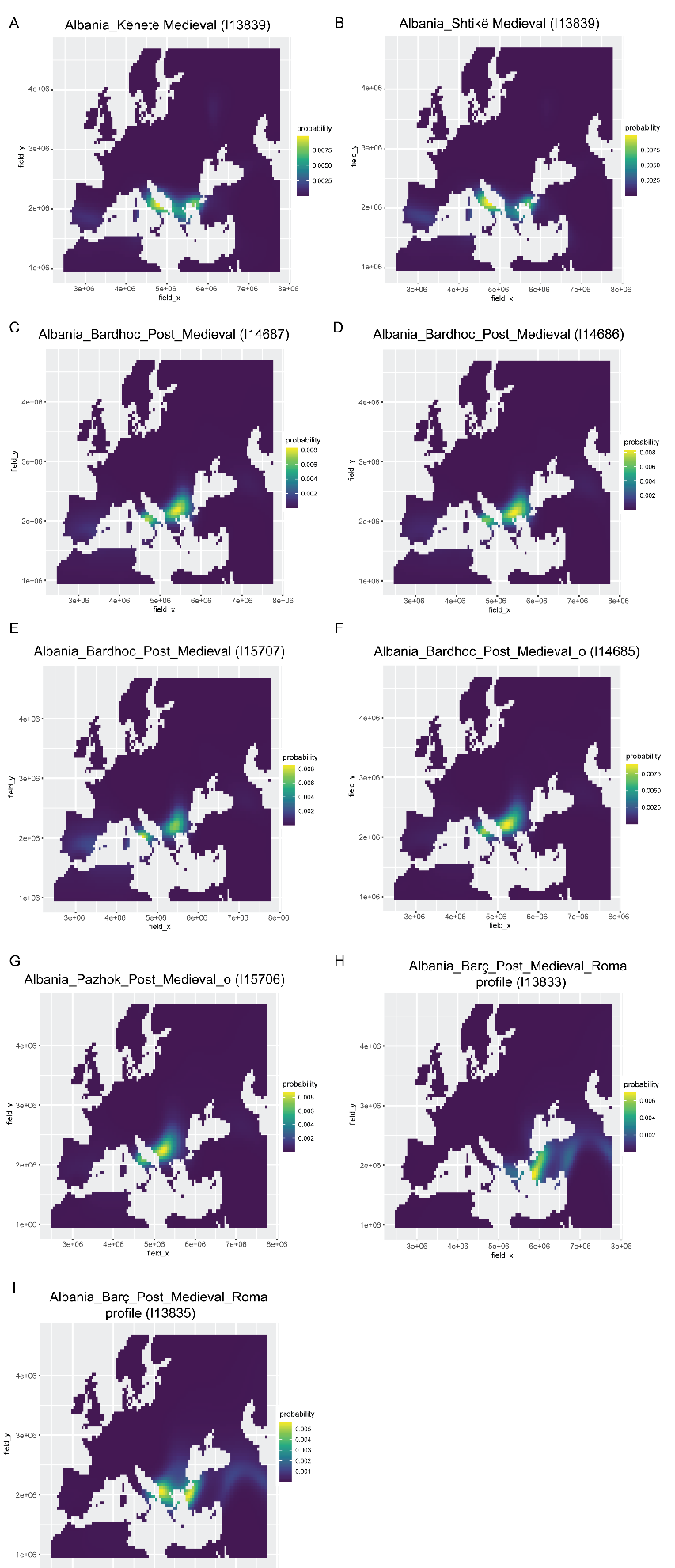

**Figure S8.** Mobest analysis of post-Medieval samples from Bardhoc, Pazhok and Barç, plotting the probability surface that identifies the highest genetic-geographical match at the mean date of the respective individual. The higher the probability surface (light yellow-green), the closer the genetic match. The latitude and longitude coordinates with the best fit for the examined individuals are provided in EPSG:3035 projection in Table S19.

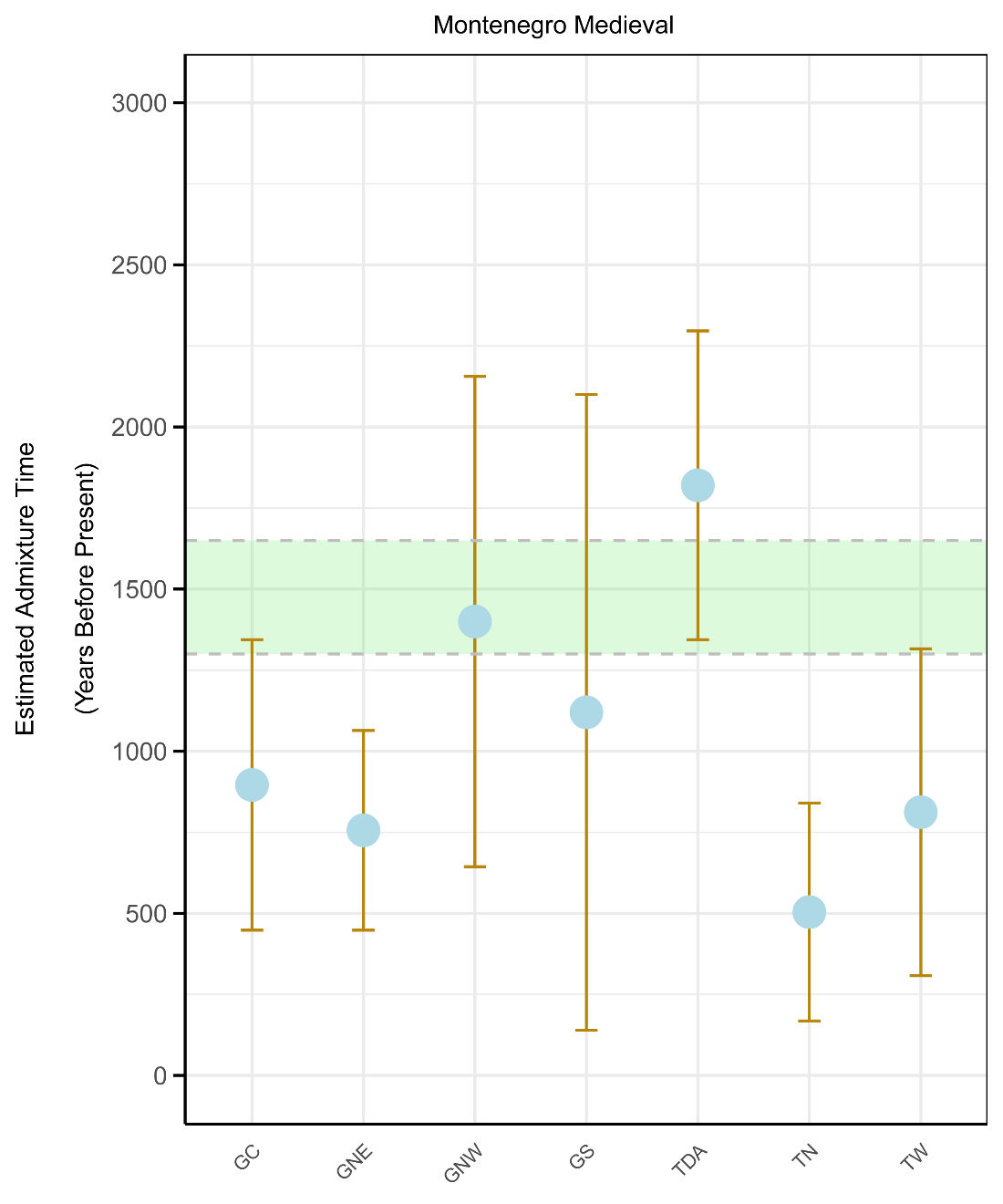

**Figure S9.** Average admixture dates of East European-related admixture (represented by Montenegro_Medieval) in present-day Albanian populations, estimated by DATES. Admixture dates for GNW, GS span a broad time range. The green dashed line represents the historically attested arrival of East European peoples in the Balkan Peninsula.

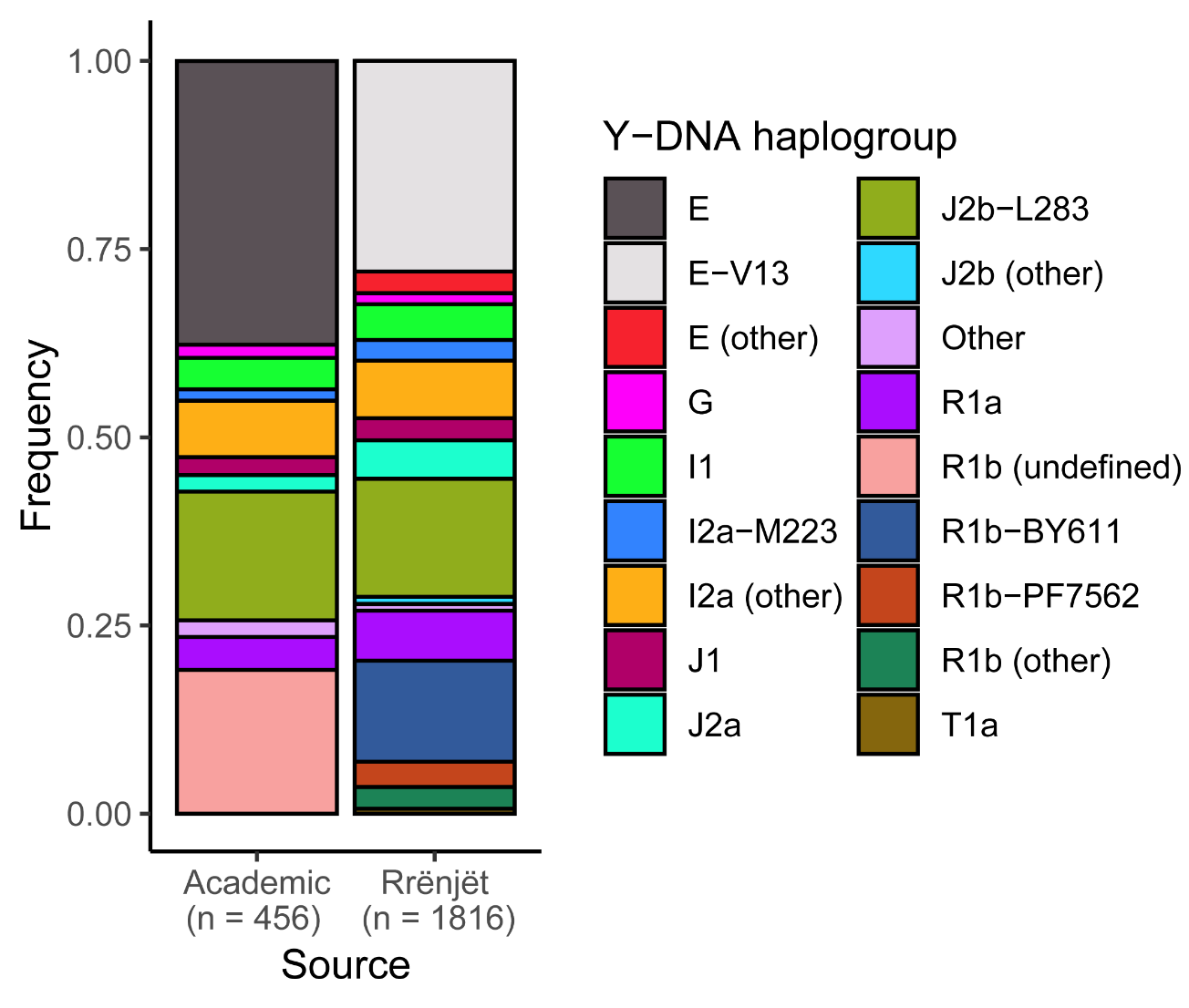

**Figure S10.** Haplogroup frequencies in present-day Albanians based on academic samples and the Rrënjët public database. Note that due to differences in STR and SNP testing, the resolution of haplogroups E and R1b-M269 in the academic samples is lower than that of Rrënjët. Migration Period haplogroups [I1, I2a (other), R1a] collectively account for 16-19% of present-day Albanian patrilines.

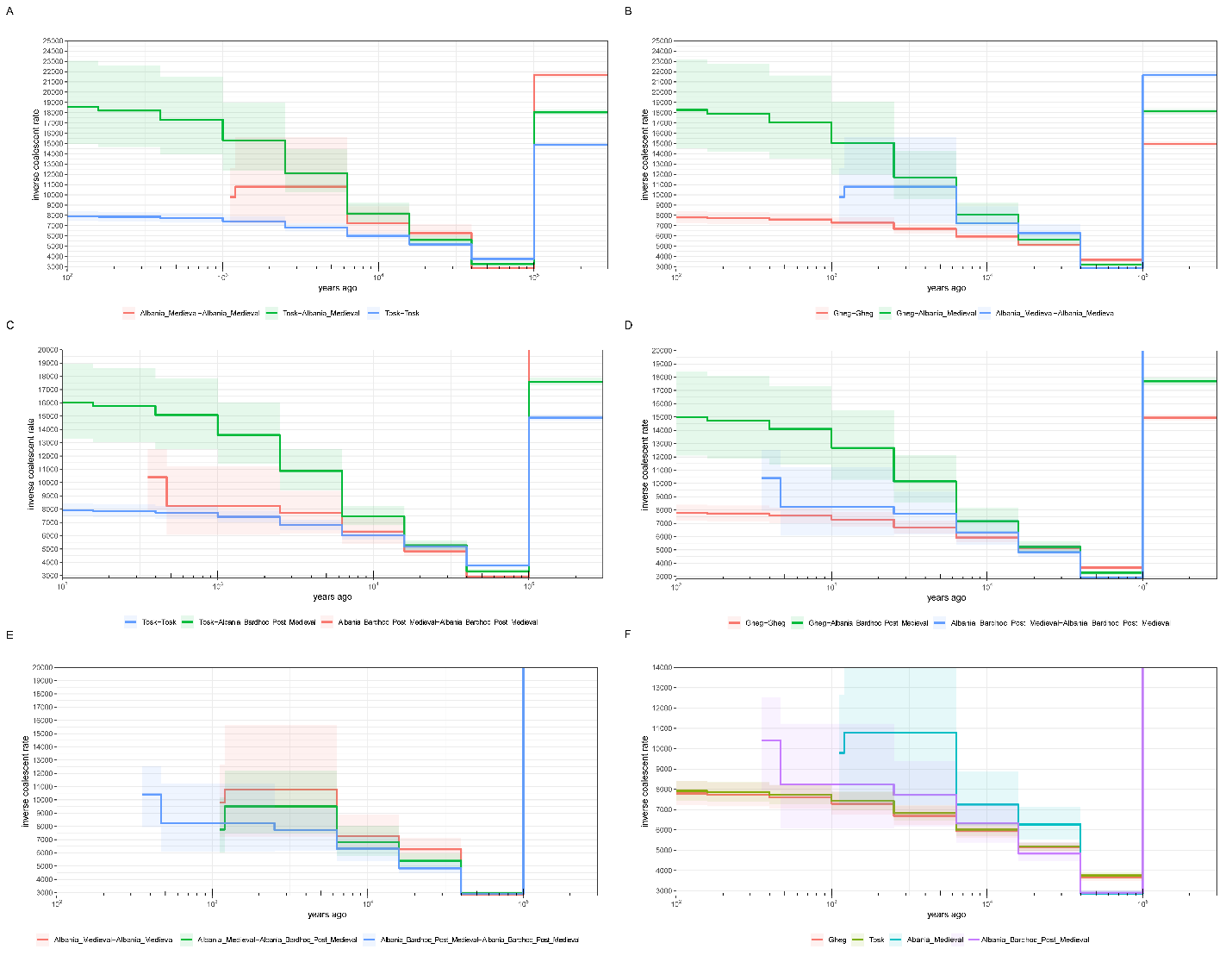

**Figure S11.** Inverse coalescence rates as estimated by Colate. A) Within-population rates for Albania_Medieval and Tosks, and between-population rates for Tosks versus Albania_Medieval. B) Same, for Albania_Medieval and Ghegs. C) Within-population rates for Albania_Bardhoc_Post_Medieval and Tosks, and between-population rates for Tosks versus Albania_Bardhoc_Post_Medieval. D) Same, for Albania_Bardhoc_Post_Medieval and Ghegs. E) Within-population rates for Albania_Medieval and Albania_Bardhoc_Post_Medieval, and between-population rates for Albania_Bardhoc_Post_Medieval versus Albania_Medieval. F) Within-population rates for Ghegs, Tosks, Albania_Bardhoc_Post_Medieval, and Albania_Medieval.

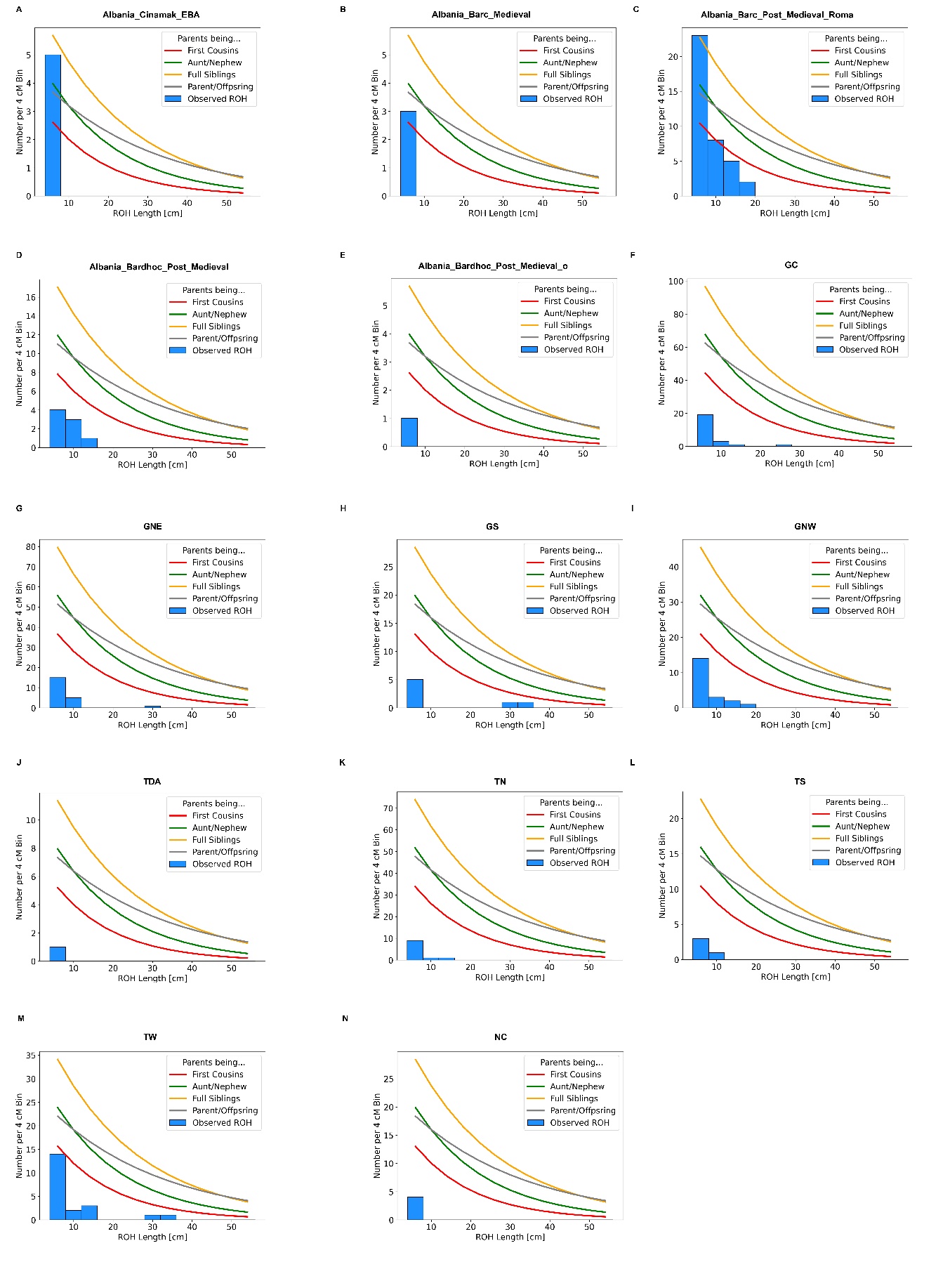
**Figure S12.** Histogram of estimated ROH lengths for ancient and present-day populations from the territory of modern Albania, alongside ROH densities as expected for varying degrees of parental relationships.

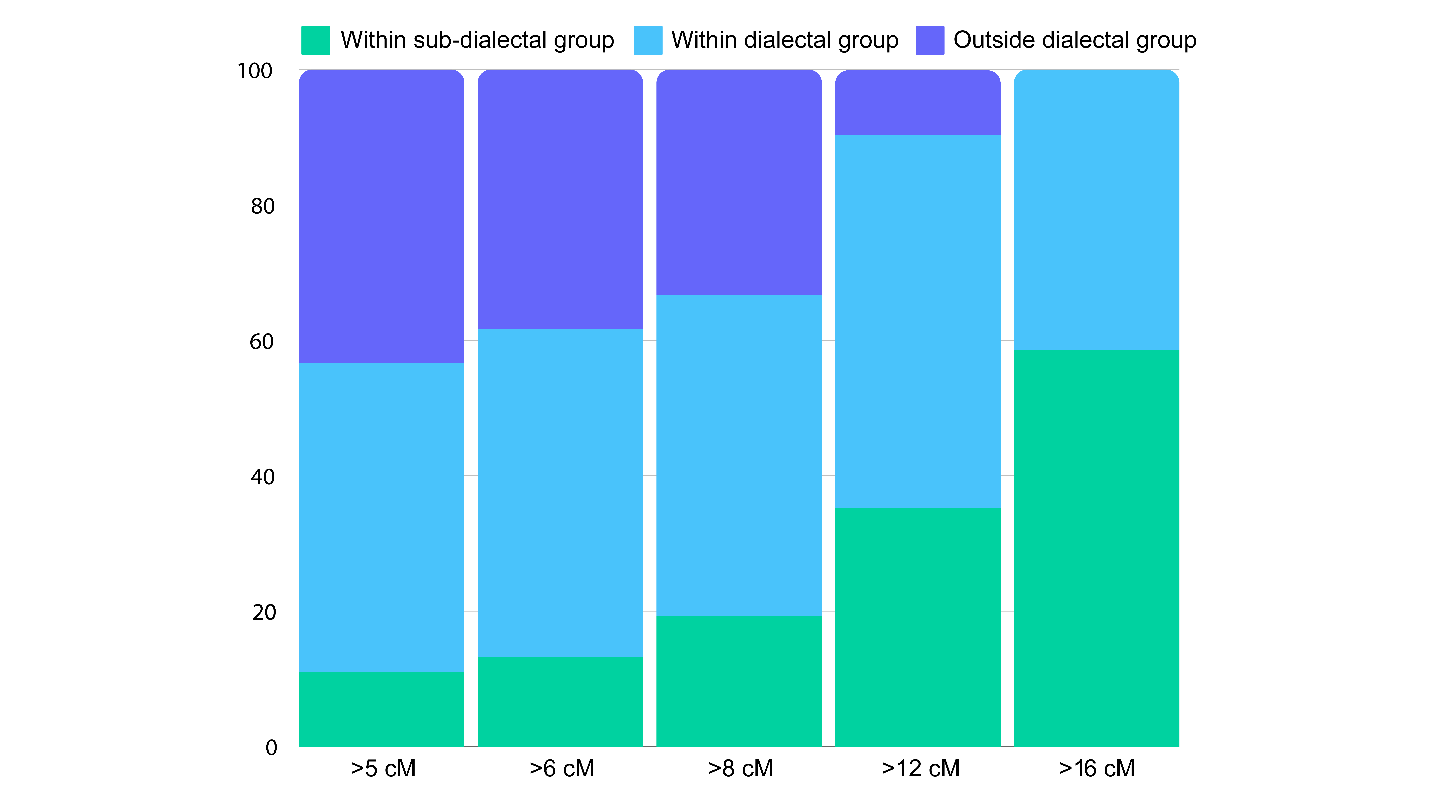

**Figure S13.** Proportion of IBD-sharing patterns within present-day Albanian sub-dialectal groups (e.g. GNE), dialectal groups (e.g. all Ghegs), and between dialectal groups (Ghegs versus Tosks), at different cM sizes. Large chromosomal blocks (>12-16 cM) show virtually no inter-dialectal IBD sharing, indicating the presence of longstanding genetic isolation between Ghegs and Tosks.

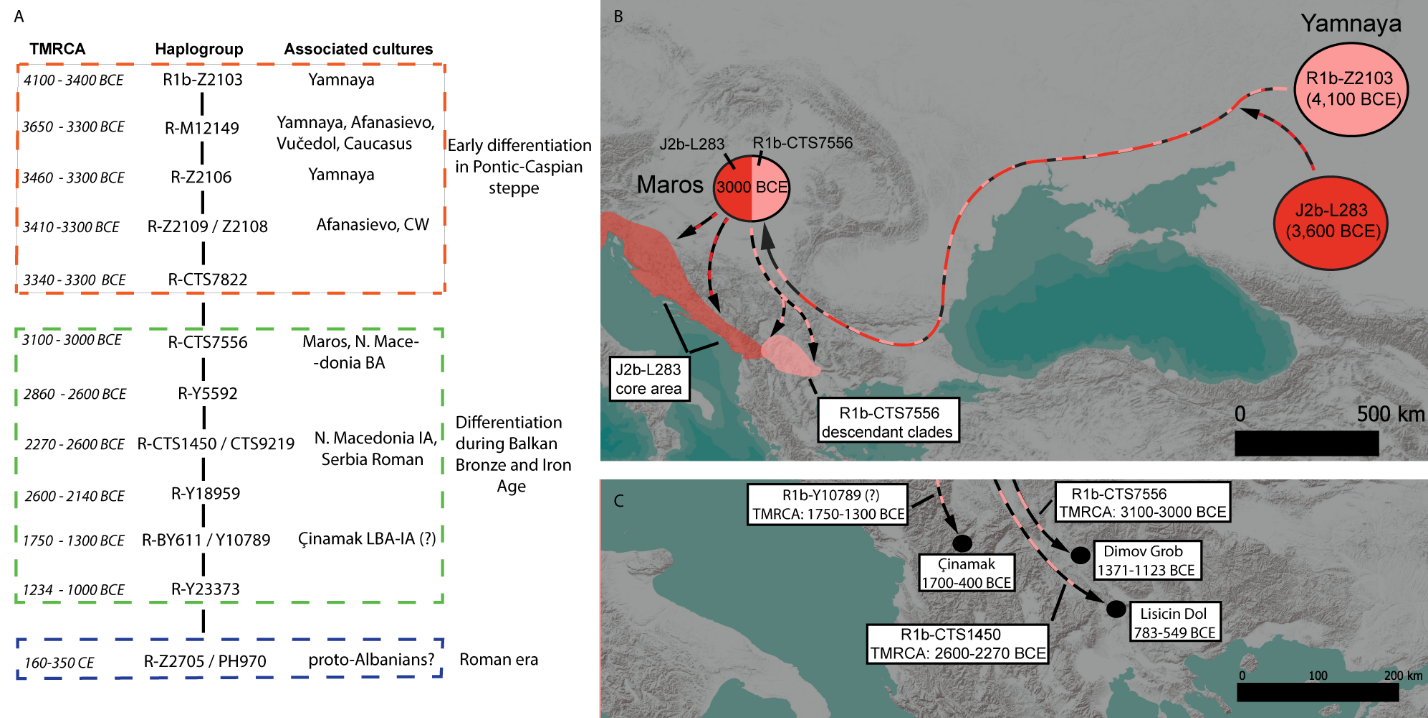

**Figure S14.** Hypothetical origin and dispersal of haplogroups R1b-CTS7566 and J2b-L283 in the Balkans during the Bronze Age. A) Schematic phylogeny of R1b-CTS7566 based on Y-full and FTDNA; B) Hypothetical reconstruction of the dispersal of R1b-CTS7566 and J2b-L283 based on phylogeny and ancient DNA samples belonging to these haplogroups. C) R1b-CTS7566 and descendant clades in the aDNA record of the Balkans.

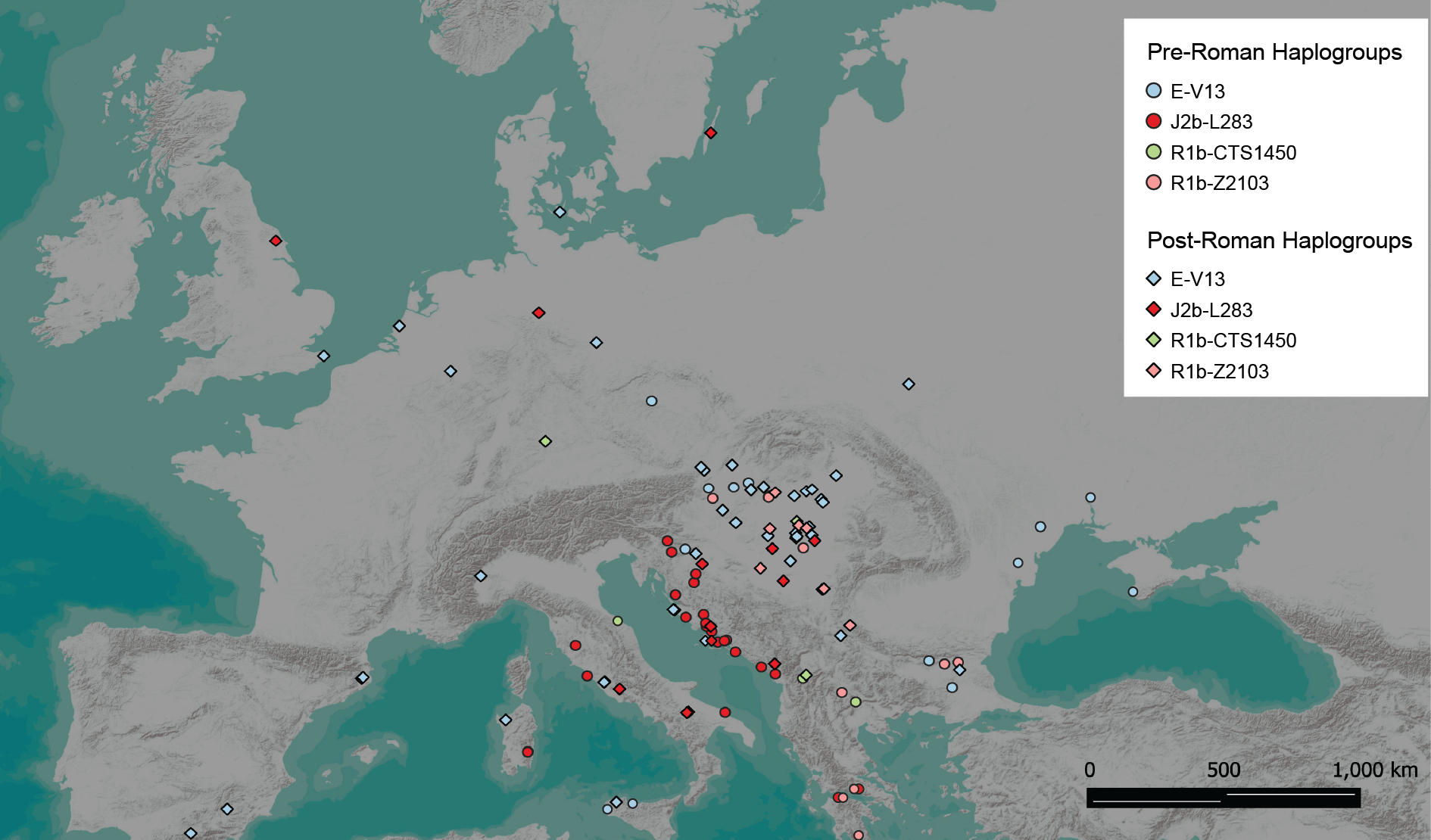

**Figure S15.** The distribution of haplogroups E-V13, J2b-L283, R1b-Z2103 and subclade R1b-CTS1450 in Europe before, during, and after the Roman period. Based on Tables S18-S26.

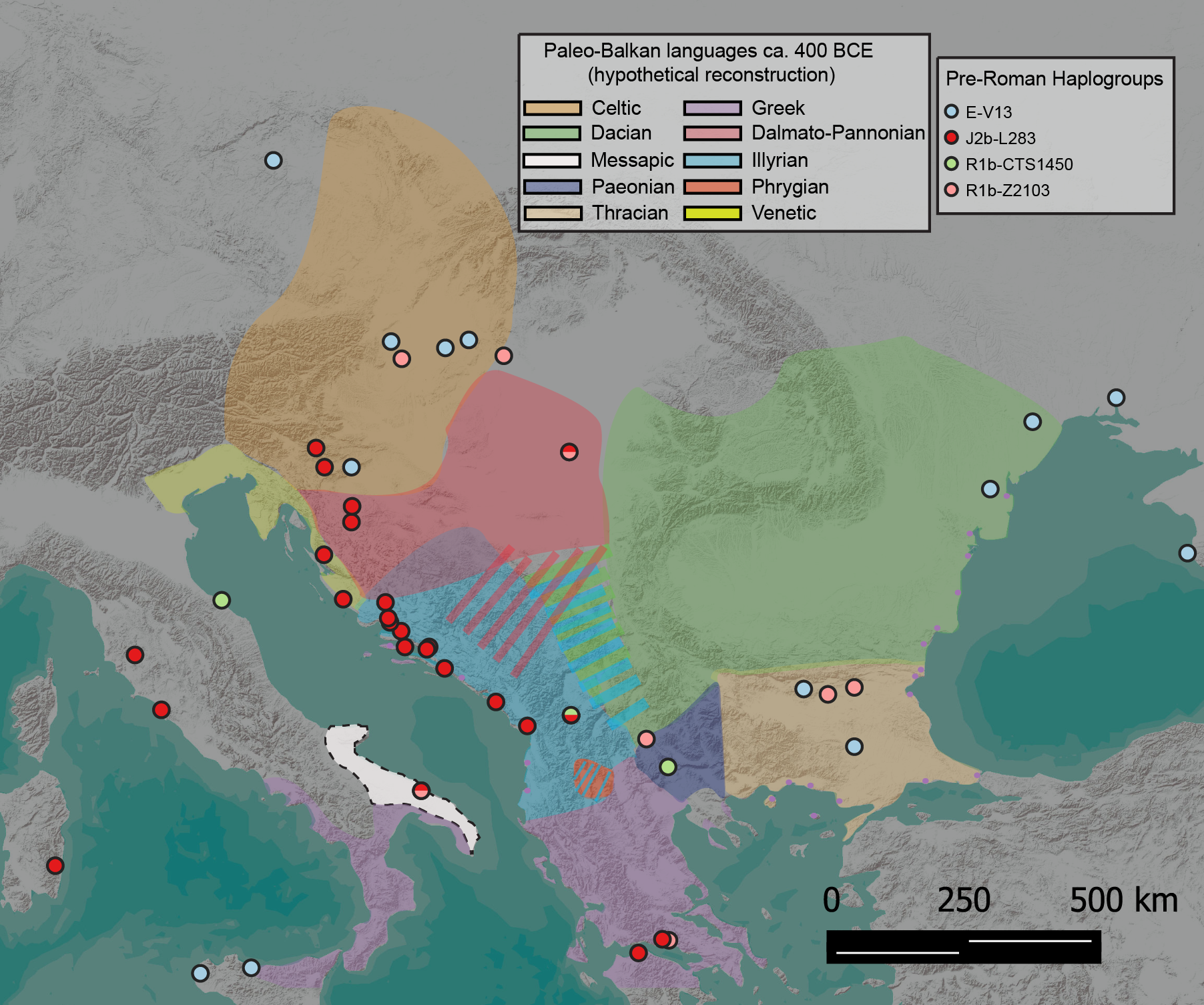

**Figure S16.** Hypothetical distribution of paleo-Balkan languages ca. 400 BCE, and their possible association with a subset of the pre-Roman haplogroups in the region. We caution that the distributional ranges provided here are based on fragmentary archaeological, historical, onomastic, and toponymic data^15,16,21–25^, and only serve as approximations. The range of Phrygian in the Balkans is poorly understood, whereas in the examined period, Chalkidiki in Macedonia is reported to have been inhabited by speakers of an undetermined language. Whether Dacian-Thracian constitute a single language, or related dialects-languages is unknown^15,16^. The same holds true for Dalmatian-Illyrian-Messapic^21,26,27^.

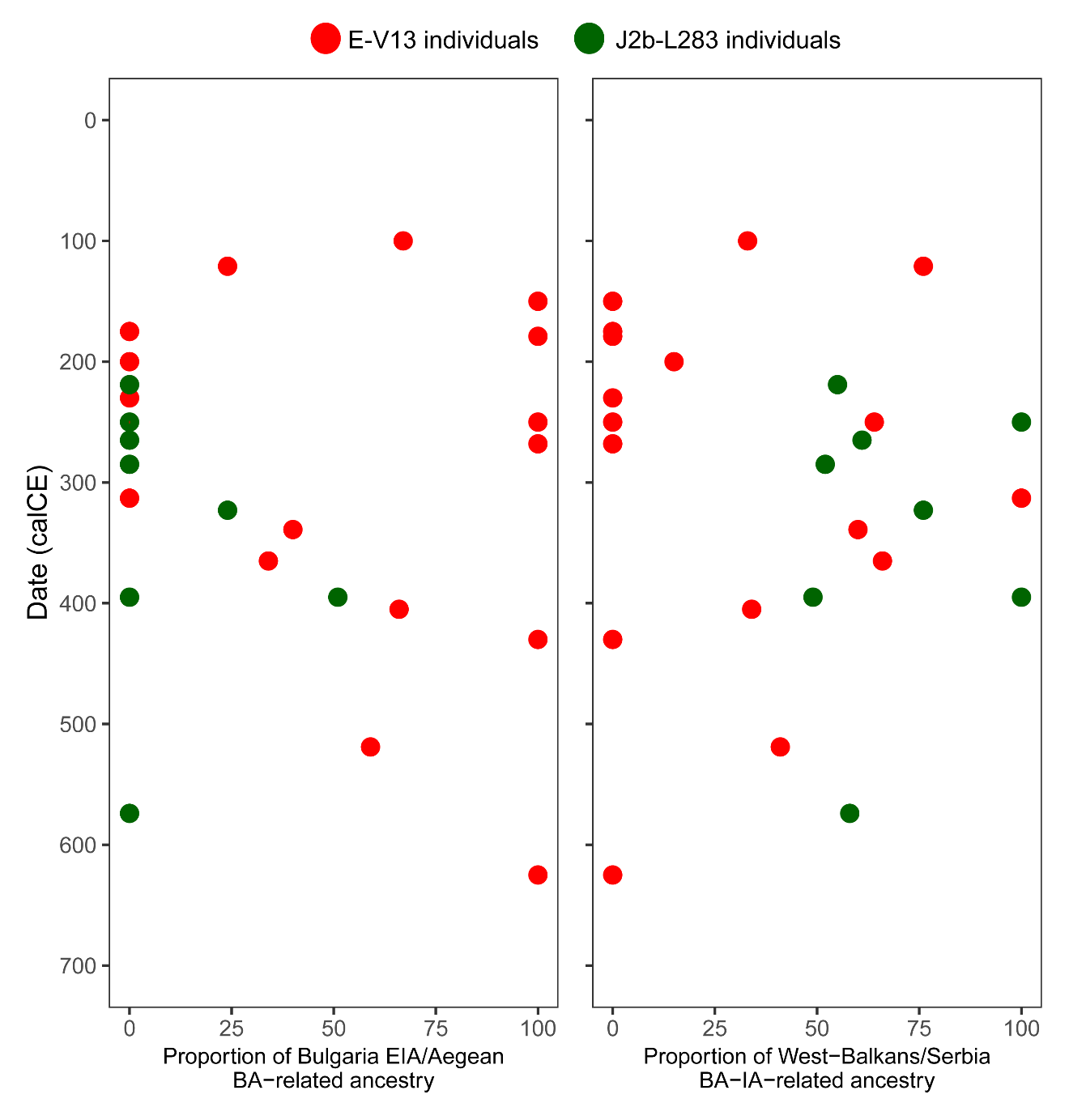

**Figure S17.** Autosomal ancestry proportions of E-V13 and J2b-L283 Roman and Avar period individuals from the Balkans and Hungary, respectively. Note that some E-V13 individuals derive none of their ancestry from either of the palaeo-Balkan sources employed here.

Supplementary Tables S1-S24 uploaded as separate .xlsx files.
